# DNA damage induces long range changes to duplex structure - a non-protein start to damage detection?

**DOI:** 10.64898/2026.03.06.709887

**Authors:** Sophie E. Fountain, Mahmoud A. S. Abdelhamid, Timothy D. Craggs

**Affiliations:** School of Mathematical and Physical Sciences, Faculty of Science, The University of Sheffield, Dainton Building, Brook Hill, S3 7HF, UK; Exciting Instruments, Block 5, Level 9 Pennine Five Campus, 18 Hawley Street, Sheffield, England, S1 4WP

## Abstract

DNA-binding proteins must quickly locate specific sites on DNA to enable replication, repair, and transcription. While sequence-specific recognition is well understood, the physical basis of structure-specific recognition remains unclear, limiting our understanding of DNA damage repair. Proteins must distinguish damaged sites within largely undamaged DNA; however, studying this is challenging due to DNA’s dynamic nature. We hypothesised that DNA damage causes changes in DNA structure, signalling protein recruitment. Using confocal single-molecule FRET, we analysed seven DNA duplexes containing modifications such as ribonucleotide, 8-oxoguanine (8–oxoG), abasic sites, nicks, and gaps, which are all involved in the base excision repair (BER) pathway. Each construct was measured with nine dye pairs in triplicate to capture changes in bending, twisting, and stretching. An automated analysis pipeline processed 162 measurements, enabling rigorous statistical comparisons. All modifications altered FRET efficiencies compared to undamaged DNA, including the subtlest change: a single oxygen difference (ribo-vs deoxyribonucleotide). Abasic sites, nicks, and gaps had the greatest effects. These findings provide direct evidence that DNA damage affects duplex structure and dynamics beyond the lesion site, suggesting DNA flexibility changes may act as an early signal for repair protein recruitment.

**GRAPHICAL ABSTRACT:** 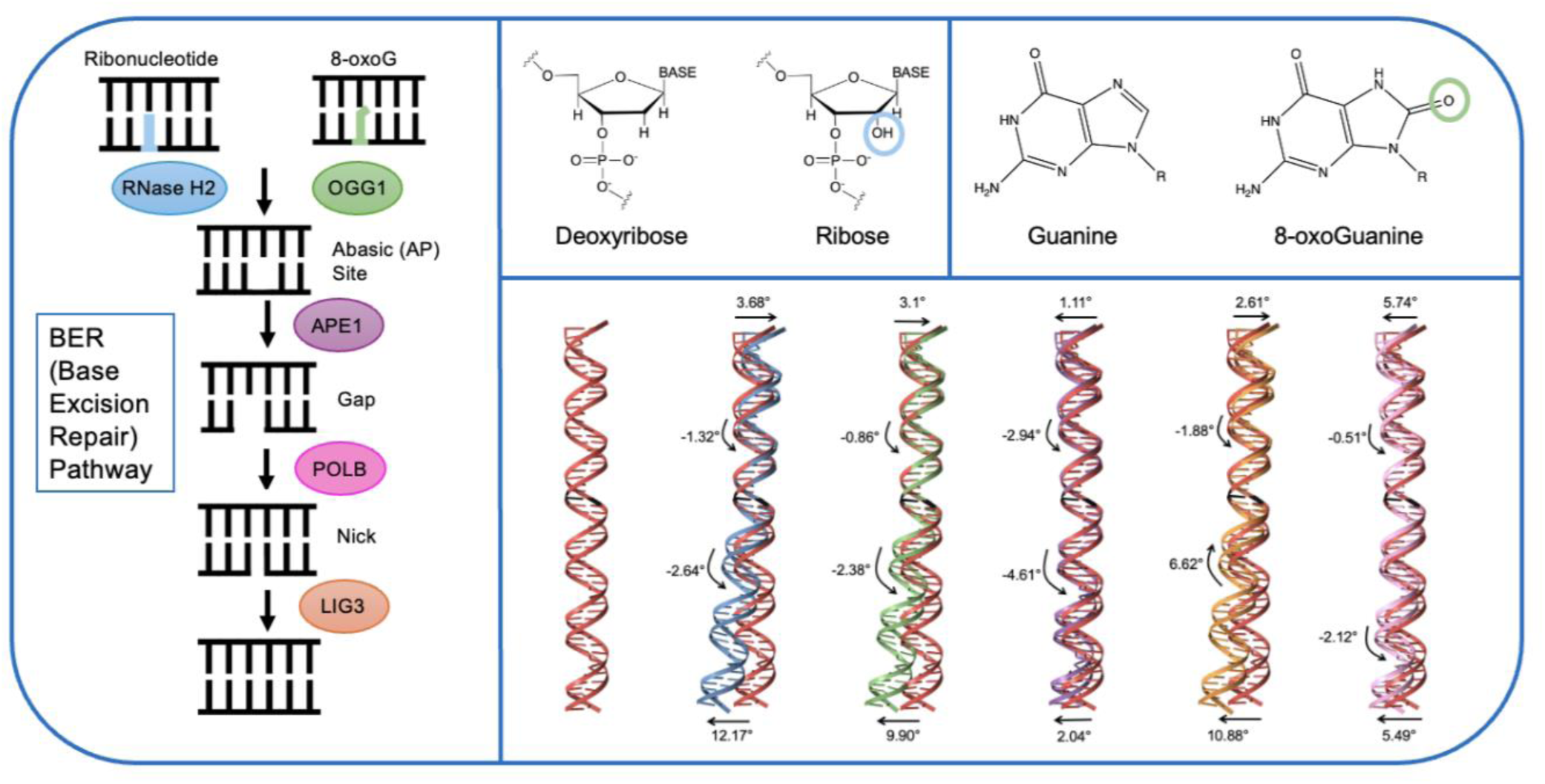

## INTRODUCTION

The integrity of DNA is crucial for the proper functioning of transcription and replication, as any damage to the nucleobases can lead to significant and potentially detrimental consequences^1^. During these processes, each base interacts with its complement, ensuring the accurate transmission of genetic information^2^. However, when a base is damaged to the extent that its hydrogen bond pairing is disrupted, it may mispair with a non-complementary base (e.g., G-A, T-C, G-G, or G-T). Such mispairing during transcription can introduce incorrect nucleotides into mRNA, which may lead to the incorporation of incorrect amino acids during translation, thereby affecting cellular function and integrity^3^. As a result, the detection and repair of damaged bases is vital for maintaining genomic stability. The primary mechanism for base repair is excision, which involves the removal of the damaged base from the DNA double helix, followed by replacement with the correct nucleotide^4^. However, the efficiency of this repair process is heavily dependent on the ability to quickly identify damaged regions within the genome^5^. To detect damage within a reasonable time frame, it is crucial to reduce the search space, ensuring that repair proteins can be directed to the damage sites efficiently. It has been proposed that a DNA segment containing a damaged base exhibits heightened molecular dynamics compared to adjacent undamaged regions, potentially triggering the response from repair proteins^6,7^. This difference in flexibility may serve as a key indicator of DNA damage, providing a more targeted approach to repair^8^. The efficiency of Uracil-DNA glycosylase (UNG) is influenced by the structural flexibility of the DNA substrate, with neighbouring bases modulating the enzyme’s activity^9^. This DNA sequence context plays a crucial role in lesion recognition and excision efficiency. To fully understand the impact of DNA damage on the structural and dynamic properties of DNA, a thorough examination of these conformational changes is required, enabling a more precise understanding of the underlying mechanisms of DNA repair^10^.

The presence of ribonucleotides in DNA can induce replication stress and genomic instability, and they have been associated with various diseases^11^. Ribonucleotides are the most abundant non-canonical nucleotides found in genomic DNA^12^. During a single round of replication in yeast, approximately 13,000 ribonucleotides may be incorporated into the genome, equating to about one ribonucleotide every 6,500 base pairs^13^. In human cells, the number of incorporated ribonucleotides per cell cycle is estimated to exceed one million^14^. This high frequency is due to the significantly higher intracellular concentrations of ribonucleotide triphosphates (rNTPs) compared to deoxyribonucleotide triphosphates (dNTPs), coupled with the imperfect selectivity of DNA polymerases^15,16^. The presence of ribonucleotides in DNA can lead to replication stress, genomic instability, and disease development^17^. They have been linked to conditions such as Aicardi-Goutières syndrome (AGS), an early-onset neuroinflammatory disorder, and Ataxia with Oculomotor Apraxia 1 (AOA1), a neurological condition associated with mutations affecting the repair of RNA-DNA junctions^18^. The accumulation of ribonucleotides in DNA can also result in chromosomal instability, gross chromosomal rearrangements, and increased mutagenesis^19^. The removal of ribonucleotides from the DNA duplex is the primary repair mechanism for most ribonucleotide lesions, making them a frequently corrected form of DNA damage^20^. Ribonucleic acid (RNA), which consists of ribose sugars, introduces RNA-like characteristics when it is incorporated into DNA^21^. Comparative studies of RNA and DNA flexibility have shown that DNA exhibits greater flexibility than RNA, further emphasising the distinct roles and properties of these nucleic acids in maintaining genomic integrity^22^.

8–oxoG is one of the most common DNA lesions resulting from reactive oxygen species (ROS), with guanine and adenine being more easily oxidised than cytosine and thymine^23,24^. The oxidation of guanine can lead to mispairing with adenine, making 8–oxoG highly mutagenic and often resulting in G-to-T transversions ^25,26^. To prevent such mutations, DNA repair glycosylases excise either 8–oxoG from 8–oxoG/C pairs or adenine from 8–oxoG/A mispairs^27^. Under normal oxidative stress conditions, an estimated 10,000 to 100,000 8–oxoG lesions occur per human cell per day^28^. Given its mutagenic potential, the accumulation of 8–oxoG is implicated in various diseases, including cancer, neurodegenerative disorders including, Alzheimer’s and Parkinson’s, and ageing-related conditions^29^. Studies indicate that impaired repair mechanisms contribute to the buildup of these lesions, leading to genomic instability and increased disease susceptibility^30^.

In DNA, the loss of a nucleobase by hydrolysis forms an abasic site. In addition to being a result of DNA damage, it is a key intermediate during base excision repair (BER)^31^. The absence of Watson-Crick bases hinders DNA polymerases from accurately identifying the appropriate nucleotide to incorporate when encountering abasic lesions^32^. As a result, multiple studies have indicated that (oxidised) abasic sites exhibit a high degree of mutagenicity^33,34^. DNA glycosylases play a critical role in the repair of damaged DNA through the BER pathway. As an intermediate step in this process, DNA glycosylases generate abasic sites^35^. These enzymes catalyse the hydrolysis of the N-glycosidic bond between the DNA base and sugar-phosphate, resulting in the release of the base. DNA glycosylases are classified into two categories: monofunctional and bifunctional. Monofunctional glycosylases remove the base, leaving behind an intact abasic site^36^. Bifunctional glycosylases additionally possess lyase activity, which allows them to cleave the DNA at the 3’ end of the abasic site. Overall, DNA glycosylases play a significant role in repairing damaged DNA through the BER pathway and exhibit remarkable conservation across various life domains^37^.

Damage to DNA can result in single-strand breaks within the double helix structure, specifically in the form of nicked, gapped, and flapped DNAs. These structural abnormalities are also essential intermediates generated during DNA replication and repair processes^38^. A nick refers to a break in the phosphodiester backbone of a double-stranded DNA molecule. It involves the breakage of the phosphodiester bond between adjacent nucleotides and can be caused by either damage or enzymatic reactions^39^. Nicks can lead to the unwinding of a DNA strand during replication. Various factors can induce nicks, including oxidative stress and ionising radiation^40^. Reactive oxygen species (ROS), including hydrogen peroxide and hydroxyl radicals, have the potential to cause damage to the deoxyribose sugar, leading to nicks within the DNA strand^41^. Nicks serve important functions in various DNA metabolism and repair processes, such as BER, nucleotide excision repair (NER), mismatch repair, removal of ribonucleoside monophosphates, and the regulation of supercoiling by topoisomerases^42^. During the nick translation process, DNA polymerase utilises a nick in the DNA strand as a reference point to identify and replace potentially damaged nucleotides. To complete the DNA repair process, DNA ligase restores the integrity of the DNA backbone after the action of DNA polymerase^43^. DNA nicks can also serve as markers for topoisomerase enzymes, which are essential for unwinding tightly packaged DNA during replication and transcription^44^. A single-stranded nick in DNA disrupts the double-stranded helical structure and reduces the molecule’s stability. Consequently, nicked DNA is more vulnerable to the physical stresses of twisting and packaging, making it more susceptible to further damage and degradation^45^.

Gaps in DNA strands arise when entire nucleotides are eliminated, resulting in an absence of bases within the strand that can form hydrogen bonds with the bases on the complementary strand. Within the context of normal cellular metabolism, DNA can undergo damage through hydrolysis or oxidation^46^. The primary repair mechanism for this type of damage involves a DNA double helix having a one-nucleotide gap. Gaps serve as crucial intermediates during DNA synthesis in BER, where the size of the gap significantly impacts the efficiency of DNA repair^47^. BER involves N-glycosylase recognising abnormal bases and breaking the glycosidic bond connecting them. Subsequently, an endonuclease cleaves the 5ʹ-side of the abnormal base, inducing alterations in the DNA structure^48^. As a result, DNA polymerase, through its exonuclease activity, excises the abnormal base^49^. A normal base is then inserted by DNA polymerase, and the region is sealed by DNA ligase. Utilising the complementary strand as a template, DNA polymerase synthesises a new strand^50^.

Base excision repair (BER) is a conserved DNA repair pathway responsible for the removal of small, chemically induced base lesions arising from deamination, oxidation or methylation (Figure 1)^56^. Substrates of BER include oxidised bases such as 8-oxoguanine (8–oxoG) and misincorporated ribonucleotides, as well as repair intermediates including abasic sites, single-strand nicks and short gaps generated during processing. BER is initiated by lesion-specific DNA glycosylases that excise the damaged base to form an abasic site, which is subsequently cleaved by an AP endonuclease to produce a single-strand break. DNA polymerase then fills the resulting gap, and DNA ligase seals the nick to restore DNA integrity^57,58^.

**Figure 1:**
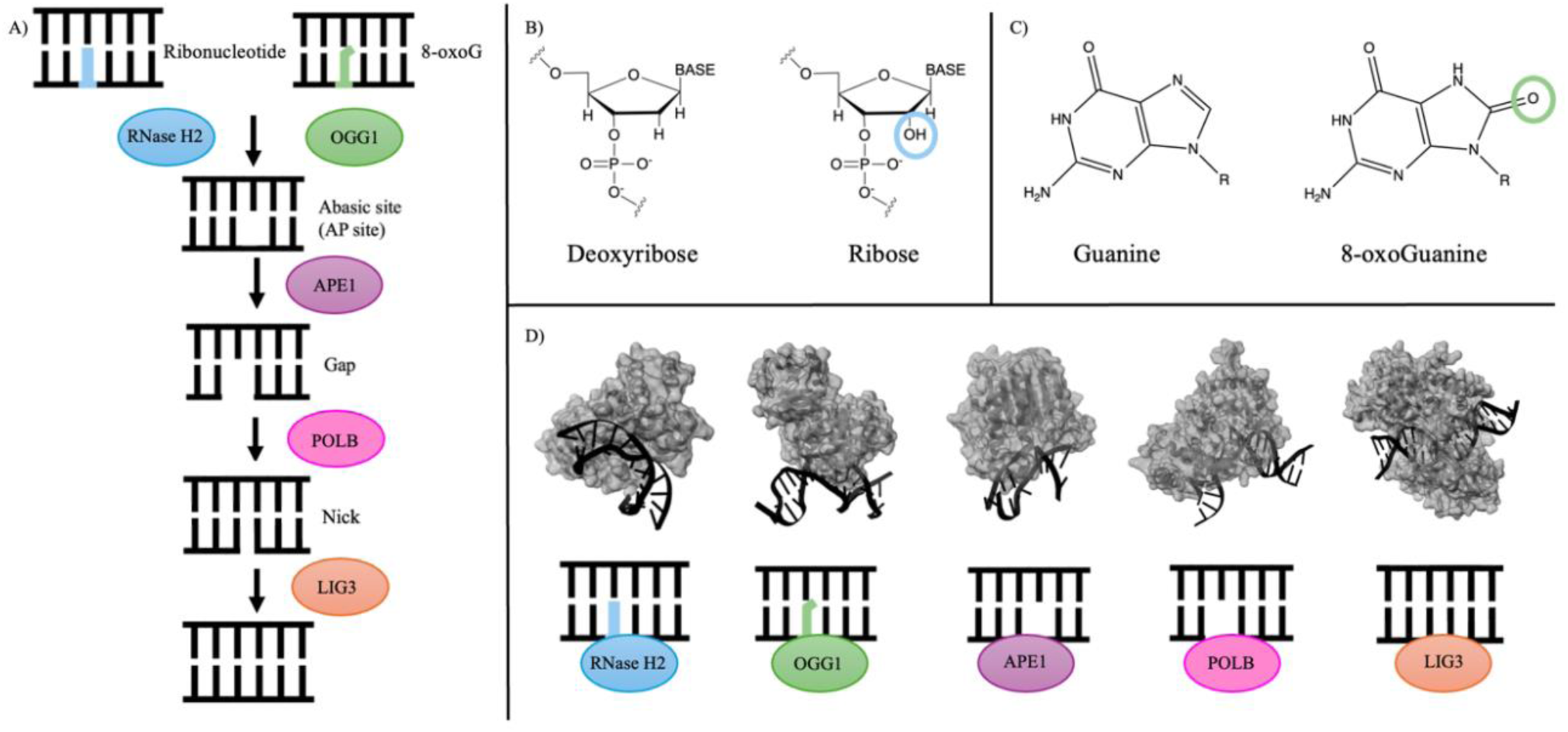
Overview of base excision repair (BER) pathways for ribonucleotide and 8–oxoG damage. (A) Schematic representation of BER, highlighting the sequential processing of ribonucleotide and 8–oxoG lesions. RNase H2 and OGG1 recognise and remove the initial lesions, leading to the formation of an abasic site (AP site), which is cleaved by APE1. The resulting gap is processed by POLB before LIG3 seals the nick to restore DNA integrity. (B) Structural comparison of deoxyribose and ribose, highlighting the 2’-hydroxyl (-OH) group in ribose. (C) Chemical structures of guanine and its oxidised form, 8–oxoG, with the oxidative modification circled. (D) Molecular representations of key BER enzymes (RNase H2 (PDB: 3PUF)^51^, OGG1 (1HU0)^52^, APE1 (1DE9)^53^, POLB (1BPX)^54^, and LIG3 (1X9N)^55^) interacting with DNA substrates, illustrating their roles in lesion recognition and repair.

**Figure 2:**
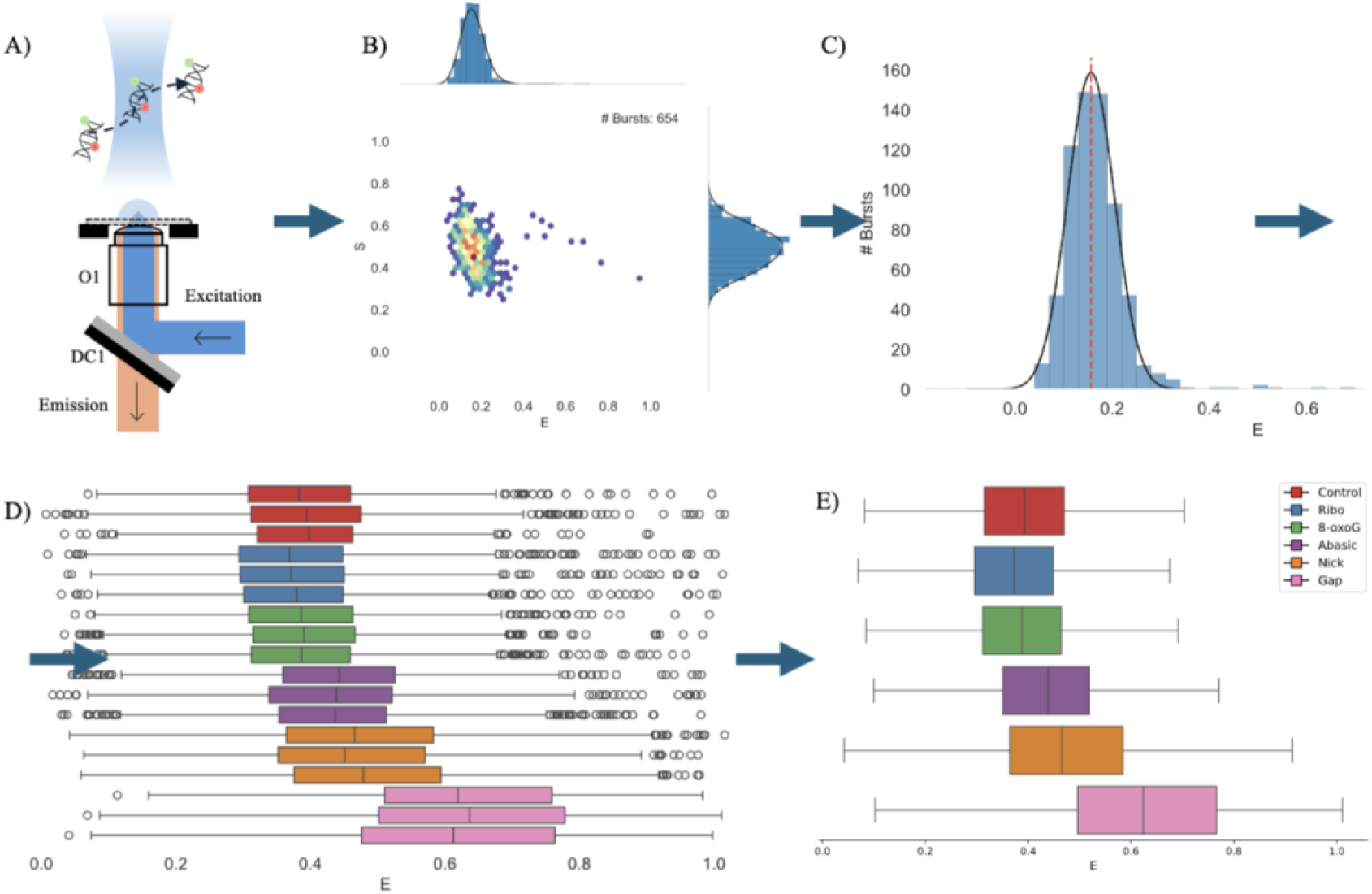
Workflow for single-molecule FRET analysis of DNA damage structures. A) Schematic of the confocal setup used for single-molecule detection B) 2D histograms of FRET efficiency (E) vs stoichiometry (S) to identify individual bursts C) 1D FRET efficiency (E) histogram and fit to extract Gaussian peak positions. D) Distribution of FRET efficiencies for various DNA constructs, including undamaged control and different lesion types (8–oxoG, ribo, abasic, nick, and gap) with repeats. E) Combined repeats for each of the damage types. Following data acquisition, the FRET of doubly labelled bursts was plotted in a E-histograms and then the E of construct was determined by a Gaussian distribution. The FRET efficiency (E) of the constructs was then plotted as box and whisker plots, and the triplicate datasets combined (Supplementary Figures 31-32).

**Figure 3:**
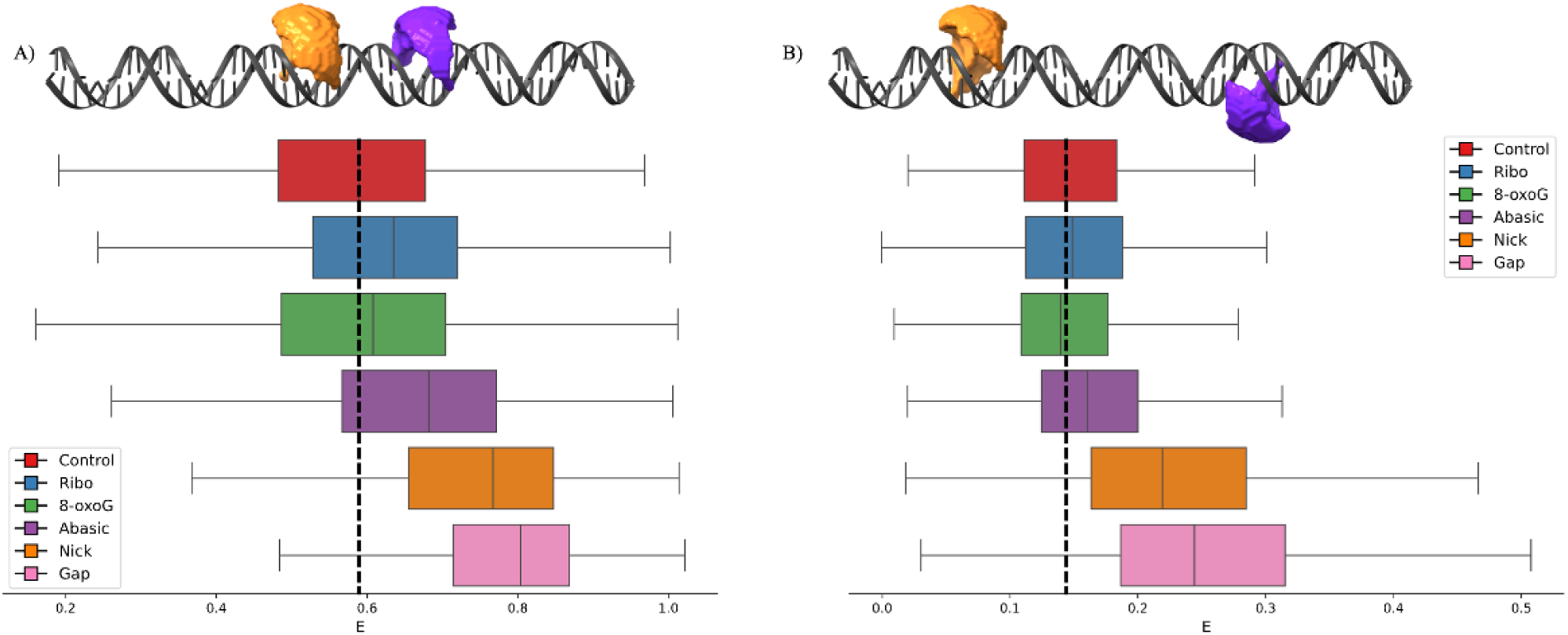
Comparison of FRET efficiency (E) for various DNA damage types at two different labelling positions. Box plot (A): Labelling position Top-7 Bottom+6 (T7B6), where the dyes, ATTO 550 (orange) and ATTO 647N (purple), are in closest proximity. At this position, FRET efficiency increases across all damage types. Box plot (B): Labelling position Top-18 Bottom+11 (T18B11). At this position, FRET efficiency also increases for most damage types when both dyes are present, except for 8–oxoG, where a decrease is observed.

In addition to BER, other repair mechanisms are vital for maintaining DNA integrity. Nucleotide excision repair (NER) plays a crucial role in repairing bulky lesions, such as those caused by UV light or chemical damage, which induce greater distortions in the DNA helix. NER removes the damaged section of DNA and replaces it with the correct nucleotides^59^. Furthermore, transcription-coupled repair (TCR), a sub-pathway of NER, specifically targets DNA lesions that stall transcription, ensuring that the transcription machinery can continue to produce RNA without disruption. These repair pathways, including BER, NER, and TCR, together contribute to maintaining genomic stability and preventing the accumulation of mutations^60^.

The objective of this study was to investigate the conformational changes in DNA structures resulting from the presence of six different types of DNA damage. Specifically, we aimed to determine whether these damages induce measurable changes in DNA flexibility, which could be detected through variations in FRET efficiency. We hypothesised that the presence of abasic sites, nicks, and gaps would lead to significant alterations in FRET efficiency due to their expected impact on DNA structure. In contrast, we anticipated more subtle changes in FRET efficiency for other types of damage, such as 8–oxoG and ribonucleotides (ribo), which are less likely to disrupt the overall conformation of the DNA duplex. By examining how DNA’s conformation responds to different damage types, this study aims to enhance our understanding of DNA damage recognition and repair mechanisms, which are crucial for maintaining genomic integrity.

## MATERIALS AND METHODS

### Sample Preparation

Oligonucleotides (TXBX series: Supplementary Table 1, NMS: Supplementary Table 2 (sequences from^61^)) were supplied by Eurogentec (Belgium), synthesised on a 200 nmol scale and purified by reverse-phase HPLC. Oligonucleotides were dissolved at approximately 100 μM in TE buffer (pH 8) and quantified using a Nanodrop. Oligonucleotides were combined to generate six different duplex constructs, each containing both a donor and an acceptor dye (Supplementary Table 3). Oligonucleotides are designated as Top (T) or Bottom (B) strands, with numerical suffixes indicating the position of the dye relative to the damage site (defined as position 0), where negative values correspond to donor dye positions upstream of the damage site on the Top strand and positive values to acceptor dye positions downstream on the Bottom strand. For each ‘TXBX’ construct, one of three donor dye positions and one of three acceptor dye positions were combined, giving nine distinct dye-pair combinations. Constituent oligonucleotides were combined stoichiometrically to yield 500 nM duplex in 10 mM Tris, 50 mM NaCl and 1 mM EDTA (pH 8). All samples were annealed by heating to 95°C, holding for 5 min, and then allowing to cool slowly to room temperature overnight.

### smFRET Measurements

smFRET measurements were performed using a home-built confocal microscope (smfBox) with alternating laser excitation (ALEX)^62^. Two diode lasers (515 nm and 635 nm, Omicron LuxX Plus) were combined into a single-mode fibre and directly modulated with a 100 µs cycle (45 µs donor excitation, 5 µs off, 45 µs acceptor excitation, 5 µs off). The output beam was collimated and cropped to 5 mm using an iris before being directed into an Olympus UPLSAPO 60× NA 1.35 oil-immersion objective via a Chroma ZT532/640rpc excitation dichroic. Fluorescence emission was collected by the same objective, focused through a 20 µm pinhole, spectrally split using a 640 nm long-pass dichroic, and detected by two avalanche photodiodes (Excelitas SPCM-AQRH-14 and SPCM-NIR-14). Photon arrival times were time-stamped using a National Instruments PCIe-6353 card, with acquisition controlled by EI-FLEX software^63,64^.

Annealed samples were diluted to ∼5 pM in observation buffer (5 mM Tris, pH 7.5; 20 mM MgCl₂; 5 mM NaCl; 0.1 mg/mL photobleached BSA), and measurements were acquired on the smfBox. For each of the 6 constructs × 9 dye-pair combinations, data were collected for 60-minutes in triplicate from three independently prepared samples using the same annealed stocks.

### Data Analysis

Data analysis was performed with Jupyter Notebooks using the FRETBursts Python module (version 0.7.1)^65^. Background in each channel was estimated by using an exponential fit of inter-photon delays. Bursts were identified by performing a dual-channel burst search (DCBS) using a photon sliding window algorithm with m = 10, F = 20 (for each channel) and DD+DA and AA thresholds = 50, to extract doubly labelled bursts from each acquisition (Supplementary Figures 1-9). Jupyter Notebooks and raw data (HDF5 files) are available at: (https://doi.org/10.5281/zenodo.17508550, https://doi.org/10.5281/zenodo.17590548, https://doi.org/10.5281/zenodo.18164375 and https://doi.org/10.5281/zenodo.18164520).

### Statistical Analysis

To determine which observed differences in FRET efficiency were significantly different (p < 0.05), the distribution of the data was assessed for normality by a combination of visual assessment of the E-histograms, quantile-quantile (QQ) plots, and the Kolmogorov-Smirnov test. All data were confirmed to deviate from a normal distribution, with the QQ plots additionally identifying that many of the measurements contained outliers, which can be expected from methods with high sensitivity to heterogeneity, such as single-molecule FRET. Consequently, the Kruskal-Wallis test was used as a nonparametric test of variance and confirmed that within all the groups, there were samples originating from different distributions. The post-hoc Dunn’s Test was then used to perform pairwise comparisons between all pairs within groups to identify which were significantly different from the others^66^. The complete results of the statistical testing can be found in the supplementary information (Supplementary Figures 10-30).

### Photophysical Characterisation

Fluorescence quantum yields (Φ) were determined for free dyes (Rhodamine B: RDB, Rhodamine 6G: RD6-G, ATTO 550, and ATTO 647N) in ethanol and for labelled DNA samples in observation buffer. UV-vis absorbance and fluorescence spectra were recorded, with measurements at five absorbance values (0.02-0.10) to avoid inner-filter effects. Quantum yields were calculated using:

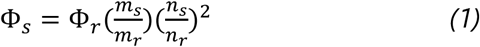

where *m* is the slope of integrated fluorescence intensity versus absorbance, *n* is the refractive index, and Φ*_r_* is the reference quantum yield (Supplementary Figure 33). All measurements were performed in triplicate (Supplementary Tables 4-6). The spectral overlap integral (J) was calculated as:

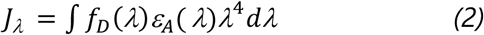

*w*here 𝑓_𝐷_ (*λ*) is the fluorescence emission of the donor and *ε*_𝐴_ (𝜆) is the acceptor molar extinction coefficient. The Förster radius (R₀) was then calculated (assuming κ^2^=2/3) using:

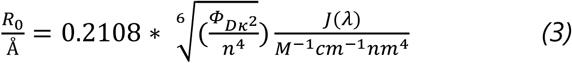

Time-resolved fluorescence lifetimes were measured for free dyes, NMS constructs, and singly labelled control duplexes (Supplementary Table 11 and Supplementary Figure 34). Fluorescence anisotropy measurements were used to analyse rotational dynamics with anisotropy calculated as:

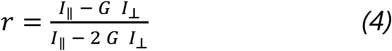

where *G* is the instrument correction factor for detection efficiencies. For high numerical aperture (NA) objectives, the anisotropy was further corrected using:

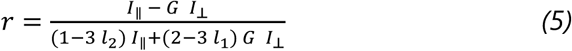

Combined restriction of donor and acceptor fluorophores (r_C,∞_) was estimated by:

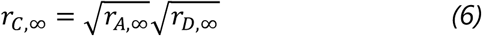

This combined anisotropy directly reflects the transfer anisotropy, as per the Soleillet theorem. The transfer anisotropy, often misrepresented, is not directly measurable because the acceptor anisotropy after donor excitation includes both the transfer and acceptor anisotropies (Supplementary Figures 35-36).

### Accessible Volume (AV) Modelling

Predicted FRET efficiencies were calculated from AV models using the software Olga (Supplementary Tables 8-10). Accessible volume (AV) clouds were generated for each of the nine different positions on the DNA where the dyes were attached. The AV clouds were calculated using the radii of the specific dye used and the linker length^67^. Orange AV clouds represent the donor dye (ATTO 550), and purple AV clouds represent the acceptor dye (ATTO 647N). The AV mean position is the central position within the AV cloud.

### Quantum Yield and Förster Radius

There were discrepancies between the predicted and measured FRET efficiencies in the control duplex (Supplementary Table 10). To assess whether these variations could be attributed to changes in the Förster radius (R₀), R₀ values were calculated for each labelling configuration using the corresponding quantum yield data (Supplementary Table 7). The measured quantum yields and spectral overlap integrals were used to determine R₀ values, which were then used to estimate distances between dyes and subsequently for rigid-body docking. Quantum yield measurements for ATTO 550 and the NMS standards (NMS 1-3) were consistent with literature values, confirming the reliability of the experimental setup (Supplementary Tables 4-6). The Top series constructs (T-7, T-12, T-18) showed slightly lower quantum yields, while the Bottom series (B+6, B+9, B+11) exhibited values of 0.53-0.54, indicating modest environmental effects on dye fluorescence.

Despite these small variations, the Förster radii determined for the NMS standards and control duplex were very similar, suggesting that differences in quantum yield cannot explain the discrepancies between predicted and measured FRET efficiencies. Recalculation of FRET efficiencies using adjusted R₀ values (Supplementary Tables 8-10) did not resolve these differences, highlighting a limitation of AV modelling to accurately predict FRET efficiencies.

### Lifetime and Anisotropy Results

The assumption of an isotropic orientation factor (κ² = 2/3) may not hold when dye mobility is restricted, and as the only unmeasured parameter in the Förster equation, κ² could significantly contribute to the observed discrepancies. Direct determination via time-resolved anisotropy would therefore be required to evaluate this effect.

Time-resolved fluorescence lifetime and anisotropy analyses provided insight into the local environments and dynamics of fluorophores across the NMS and TXBX series (Supplementary Table 11). The fluorescence lifetime of free ATTO 550 (3.6 ns) matched literature values, while labelled NMS constructs showed slightly longer lifetimes (3.73-3.82 ns), indicative of a more restricted environment. The T-series displayed modest variation (3.59-3.90 ns), reflecting subtle positional effects. ATTO 647N-labelled NMS 4 exhibited an extended lifetime (4.08 ns versus 3.5 ns for the free dye), again suggesting limited dye mobility.

Fluorescence anisotropy further revealed differences in rotational freedom (Supplementary Table 12). Free dyes showed very low residual anisotropy (ATTO 550: 0.009 ± 0.008; ATTO 647N: 0.011 ± 0.007), consistent with unrestricted rotation. In contrast, the T-series exhibited elevated values (T-7: 0.138 ± 0.006; T-12: 0.102 ± 0.003; T-18: 0.053 ± 0.011). Bottom-strand constructs showed a similar trend (B+6: 0.033 ± 0.010; B+9: 0.027 ± 0.001; B+11: 0.021 ± 0.005). However, these values are not dissimilar to the NMS residual anisotropies of 0.073 ± 0.018, 0.099 ± 0.011, 0.084 ± 0.009 and 0.027 ± 0.009 for NMS 1-4, respectively. This indicates that differences in rotational freedom are unlikely to account for the discrepancies between measured and predicted FRET efficiencies.

### Calculation of Dye Coordinates for Damage Constructs

As a result of these discrepancies, new accessible volume (AV) mean positions were determined for the six dyes in each damaged DNA construct by converting experimentally measured FRET efficiencies (E) to estimate inter-dye distances (R) using:

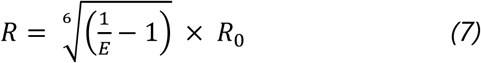

where 𝑅_0_ is the experimentally determined average Förster radius. The calculated distances were not treated as absolute, including for the control duplex (Supplementary Table 14). Rather, they were used to compute scalars by normalising each damaged construct to its corresponding undamaged control. Application of the scalars to the control-model inter-dye distances enabled generation of new inter-dye distances for the damaged constructs. All nine inter-dye distances per construct were used as constraints in a non-linear optimisation procedure (L-BFGS-B algorithm) to determine final dye coordinates in 3D space by minimising the total squared deviation between calculated and target distances through multilateration^68^.

### Rigid Body Docking

DNA bending and twisting were quantified using bespoke Python scripts built upon Biopython, employing singular value decomposition (SVD)-based structural alignment to compute optimal rotation matrices, helical axes, and bend/twist decompositions directly from PDB coordinates. Each duplex was parsed using the Bio.PDB.PDBParser() module and left and right DNA segments were defined relative to the damage site. SVD-based superposition was used to determine the principal helical axes and optimal orientation matrices for each half of the duplex. Angular deviations were calculated from shifts in the mean AV positions between control and damage-containing constructs, yielding total deformation angles with their bending and twisting components. The pivot point, corresponding to the base pair adjacent to the lesion, was used as the rotational origin for all coordinate transformations. Modified atomic coordinates were exported as new PDB files for structural visualisation. Jupyter notebooks of modelling and rigid body docking analysis are available here: https://doi.org/10.5281/zenodo.18670016.

### Burst Variance Analysis (BVA) for FRET Distributions

Burst Variance Analysis (BVA) was performed using FRETBursts to compute the standard deviation of sub-burst FRET efficiencies for each photon burst, allowing identification of static versus dynamic FRET populations by comparing observed fluctuations to those expected from the shot-noise limit. BVA looks for deviations from an expected sub-burst E variance given the E of the whole burst^69,70^. On the BVA plot, the standard deviation values clustering around the expected standard deviation indicate static FRET, whereas bursts with standard deviation values significantly above the confidence interval indicate within-burst dynamics (Supplementary Figures 39-47).

## Results

### Statistical Significance of FRET Efficiency Changes

Single-molecule analysis is used because it enables direct observation of individual biomolecules rather than population-averaged behaviour. This approach reveals distinct conformational states and their relative abundances, providing a more accurate view of structural dynamics and transitions. Because this method provides distributions rather than single averaged values, it allows differences in FRET efficiency between conditions to be evaluated with appropriate statistical tests. These tests can then determine whether the presence of DNA lesions leads to significant changes in FRET efficiency.

Dunn’s test provided robust evidence for significant and distinct changes in FRET efficiencies across various DNA damage types and labelling positions, indicating alterations in the DNA duplex conformation. For each labelling position, consistency across replicates was confirmed, with p-values consistently above 0.05 for each repeat of each labelling position across all damage types. This internal consistency increased confidence and reliability to perform comparisons between the various damage types and the undamaged control.

The abasic, nick, and gap structures consistently exhibited statistically significant differences in FRET efficiency across all tested labelling positions (p < 0.05). These damage types induced substantial structural deviations that propagated broadly through the duplex. The complete loss of a base in an abasic site and the backbone discontinuities of nicks and gaps are expected to impose significant structural alteration, altering both local and potentially global DNA dynamics.

The ribo modification showed a more nuanced effect. Significant differences in FRET efficiency were observed at positions T12B9, T12B11, T18B9, and T18B11. However, less pronounced changes were observed at T7B6, T7B9, T7B11, and T18B6, with the two constructs: T7B9 and T18B6 not reaching statistical significance. This indicates that a single ribonucleotide incorporation can induce detectable, albeit localised, conformational changes, with some regions being more sensitive to its presence. The 8–oxoG modification also exhibited significant differences at multiple positions (T7B6, T7B9, T12B6, T12B9, and T18B11), but T18B6 and T18B9 showed smaller, non-significant changes. This suggests a pattern of influence that is more restricted than abasic sites, nicks, or gaps, but more widespread than the ribonucleotide. Notably, T12B6 and T18B6 consistently showed less sensitivity to damage-induced changes across at least two different damage types, suggesting these might be less conformationally responsive regions.

### Positional Alterations in DNA Duplex Conformation

Analysis of FRET efficiencies provided a detailed picture of how different DNA modifications influenced the conformations of the duplexes. Dyes were positioned at nine distinct sites along the duplex, allowing structural changes to be monitored at multiple locations. For each damaged construct, FRET efficiencies were converted into distance estimates, which were then used to calculate scalars from the corresponding undamaged control. Transformations were then applied to the control-model dye coordinates, and new three-dimensional dye positions were determined by multilateration. Changes were assessed across three categories of dye positions: proximal, intermediate, and distal to the lesion, on both the Top (T) and Bottom (B) strands (Supplementary Table 14). These position-specific changes characterise how the different lesions alter DNA structure. The observed local and propagated structural distortions are likely to modulate lesion recognition and processing, with potential mechanistic implications for BER. Comparing responses across all positions and lesion types highlights how similar structural perturbations may affect repair outcomes in diverse genomic contexts.

### Proximal Positions (T-7 and B+6)

At the T-7 position, nick, gap, and ribonucleotide modifications consistently caused the dye to shift further from the DNA backbone, suggesting localised unwinding or increased flexibility at this site. In contrast 8–oxoG, had minimal influence on dye position, indicating only minor structural changes. The abasic site, however, resulted in the dye moving closer to the backbone, implying a more compact local conformation or helical bending around the site of base loss. Abasic sites are known to destabilise the DNA helix, disrupting base stacking and causing localised unwinding, as shown in both NMR studies and molecular dynamics simulations^71,72^. These effects likely account for the dye’s apparent movement closer to the backbone in the presence of such abasic sites.

At the B+6 position, nick and gap modifications again caused the dye to move closer to the damage site, with the nick inducing the most pronounced effect. Specifically, 8–oxoG causes bending and partial unwinding, while nicks and gaps enhance local flexibility, allowing the dye to approach the lesion site more closely^73,74^ ^75^. This disruption of the helical structure may allow the dye to move slightly away from the backbone.

Overall, T-7 and B+6 demonstrate that nicks and gaps are the most structurally disruptive, consistently reducing the dye’s distance to the damage site through enhanced flexibility and bending. Meanwhile, abasic sites and oxidative lesions primarily contribute to subtle unwinding or distortion, depending on their position and context.

### Intermediate Positions (T-12 and B+9)

Intermediate positions (T-12 and B+9) showed a similar trend. At T-12, nick and gap consistently brought the dye closer to both the backbone and the damage site, with gap having the largest effect, followed by nick, 8–oxoG, ribose, and abasic lesions. Structural effects of the nick and gap, such as increased flexibility and local unwinding, support this observation. Previous studies confirm that these discontinuities can lead to local structural changes^76, 77, 78^.

At B+9, most lesions had minimal impact, though gap and nick again shifted the dye closer to the lesion. This minimal variation could be attributed to the fact that B+9 lies at an intermediate position where the structural influence of DNA lesions is not as pronounced, as noted in other studies that have observed more subtle effects of certain lesions in regions farther from the damage site^79, 80, 81^.

Data from T-12 and B+9 show that backbone discontinuities impact DNA structure even at intermediate distances, while base and sugar lesions cause subtler effects that diminish with distance from the damage.

### Distal Positions (T-18 and B+11)

At distal positions (T-18 and B+11), lesion effects varied more. At T-18, nick, gap, and ribo brought the dye closer to the DNA backbone, whereas 8–oxoG and abasic lesions moved it further away. Nick caused the most significant shift, followed by gap, ribo, 8–oxoG, and abasic. These modest shifts suggest that, although 8–oxoG can induce local unwinding, its effects are too subtle at this distance to cause major dye displacement^82^. Literature suggests that 8–oxoG can cause local unwinding of the DNA, resulting in slight adjustments in dye position^83^.

At B+11, gap produced the largest shift toward the backbone, while abasic lesions caused a slight reduction in dye distance. Nick, 8–oxoG, and ribose induced minimal changes, but the overall trend remained: gap and nick prompted the most pronounced movements. These observations align with earlier structural studies showing that nicks and gaps substantially alter DNA flexibility, bringing dyes closer to lesion sites^84, 85^.

Across both T-18 and B+11, the consistent observation was that nick and gap lesions caused dyes to move closer to the damage site, indicative of lesion-induced DNA bending or compaction. In contrast, ribonucleotide, 8–oxoG, and abasic lesions had more variable effects, typically associated with DNA unwinding or torsional strain rather than pronounced bending (Supplementary Table 14 and Supplementary Figure 37-38).

### Overall

Across all positions, the observed dye movements illustrate how different lesions perturb DNA structure in a position-specific and lesion-specific manner. Backbone discontinuities, such as nicks and gaps, consistently induce bending, unwinding, or increased flexibility, which propagate along the duplex from proximal to distal regions, whereas base and sugar modifications (abasic sites, 8–oxoG, ribonucleotides) generally produce subtler, more localised distortions. By mapping these structural consequences across multiple positions, we obtain a detailed view of how lesions reshape the duplex beyond the immediate site of damage.

These structural insights are directly relevant to base excision repair (BER), as the ability of repair enzymes to recognise, access, and process lesions is influenced not only by the chemical nature of the damage but also by alterations to the surrounding DNA geometry. Regions of enhanced bending or flexibility may facilitate enzyme binding and strand interrogation, whereas more rigid or minimally distorted regions may impede lesion recognition^86^. Overall, integrating FRET-derived distance changes with structural mapping enables us to examine lesion-induced conformational changes.

### Bend and Twist of DNA constructs with different damages present

To assess lesion-induced structural changes beyond simple shifts in dye-labelled positions, each construct was compared with an ideal B–form control duplex (shown in red in Figure 4). Using this reference, we quantified how individual lesions perturb the DNA geometry, capturing both local distortions at the lesion site and more global changes in curvature and helical twist. By combining these structural measurements with observed dye displacements, we could directly relate lesion-induced deformations to potential effects on recognition and processing by the base excision repair (BER) machinery. This approach provides a coherent framework for interpreting the subsequent results, allowing a systematic comparison of bend, twist and dye displacement across different lesion-containing constructs and highlighting their structural and functional consequences.

**Figure 4:**
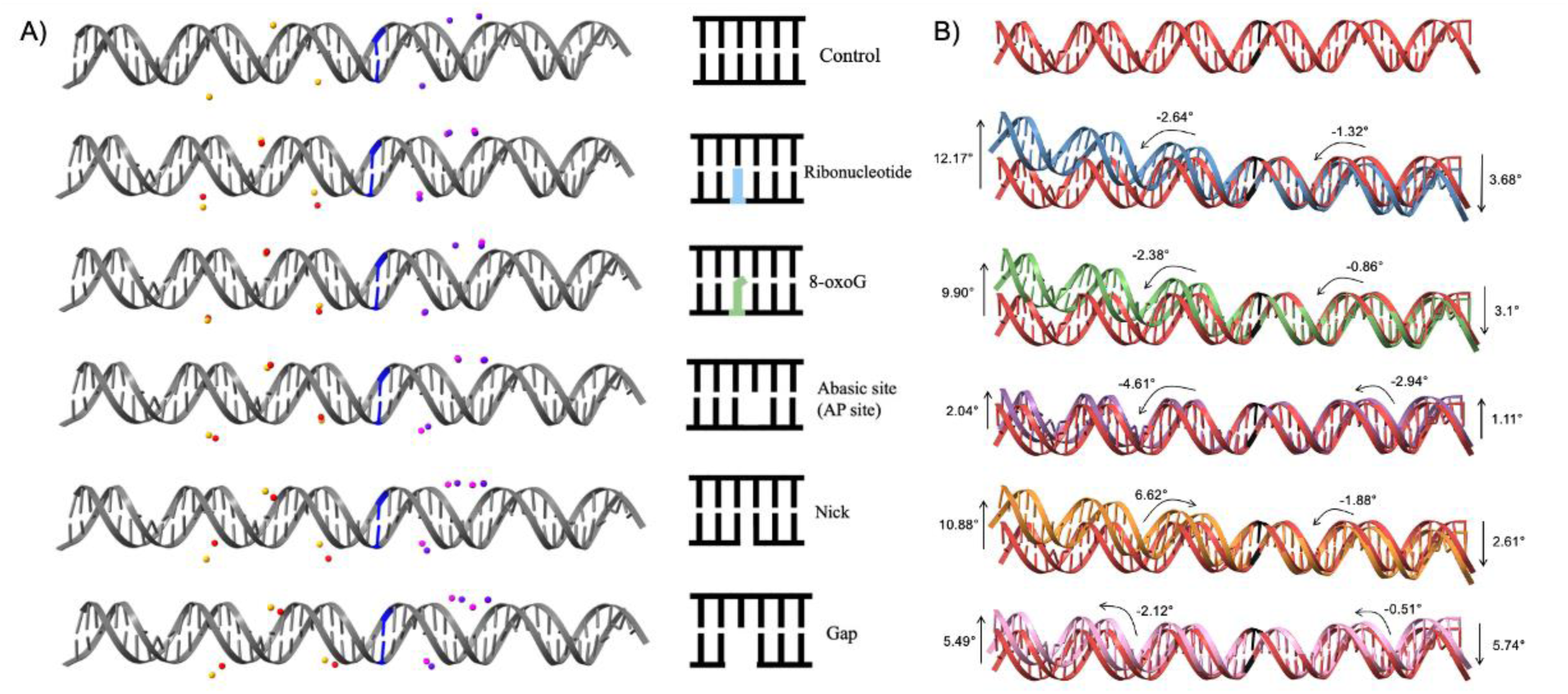
(A) Schematics illustrating different DNA damage positions (highlighted in blue) and their effects on the mean position of the AV cloud. The control position of the donor dye ATTO 550 is shown in pale orange, with its shifted position shown in dark orange. The control position of the acceptor dye ATTO 647N is shown in dark purple, with its shifted position shown in pink. (B) Models of the control duplex structure in red and the outputs of the rigid body docking for the damaged structures based on the movement of the dye AV mean positions, with angles for bend and twist of the DNA.

### Ribonucleotide

The ribonucleotide construct showed the largest bending change, with a pronounced left-handed bend of 12.17°. The helical twist was reduced on both sides of the lesion, with values of -2.64° and -1.32°, consistent with local under-twisting relative to the control. This substantial bending and unwinding suggest that the ribonucleotide strongly disrupts local base stacking and phosphate backbone geometry.

## 8–oxoG

The 8–oxoG duplex displayed a smaller left-handed bend of 9.90°, accompanied by modest under-twisting (−2.38° and -0.86°) on either side of the lesion. Although the overall deformation was less severe than that of the ribonucleotide construct, the persistence of under-twist indicates a local relaxation of the helical winding, likely arising from weakened hydrogen bonding and altered base orientation due to the oxidised guanine.

### Abasic site

The abasic site construct adopted a more relaxed conformation, with a subtle bend of 2.04° on the left and 1.11° on the right. However, the local twist values (−4.61° and -2.94°) indicate pronounced under-twisting, suggesting that base loss leads to local unstacking and increased flexibility while reducing torsional strain.

### Nick

The nicked duplex exhibited a pronounced bend, with bending angles of 10.88° on the left and 2.61° on the right. The local twist values (+6.62° and -1.88°) reveal strong over-twisting near the discontinuity and slight under-twisting on the opposite side. This torsional asymmetry indicates that strand breakage relaxes helical constraints, enabling local overwinding adjacent to the nick while maintaining moderate flexibility in neighbouring regions.

### Gap

The gap-containing duplex displayed a moderate bend, with bending angles of 5.49° on the left and 5.74° on the right, smaller than that of the nick but greater than the abasic site. The twist values (−2.12° and -0.51°) indicate mild under-twisting on both sides of the lesion, reflecting partial helical relaxation and disruption of base stacking across the missing nucleotide.

### Overall

Overall, the results reveal two distinct deformation modes within the modified duplexes. Chemical modifications such as ribo and 8–oxoG induce bending, accompanied by helical unwinding. Strand interruptions, including nicks and gaps, favour bending along the axis of the DNA with variable twist responses. The abasic site remains the least bent yet exhibits pronounced under-twisting, reflecting enhanced local flexibility and reduced torsional strain. Overall, these conformational signatures demonstrate how different lesion types uniquely perturb DNA architecture, modulating both bending direction and helical tension around the site of damage. Together, these observations illustrate how the unique structural perturbations induced by different lesion types can modulate BER efficiency and the dynamic processing of DNA damage across the genome.

### Burst Variance Analysis (BVA) for FRET Dynamics

Burst Variance Analysis (BVA) was used to gain detailed insight into the dynamic behaviour of FRET distributions, distinguishing static conformations from interconverting states within single bursts. BVA analyses photon arrival times to detect within-burst changes in E, assessing deviations from the expected sub-burst variance given the overall FRET efficiency.

BVA revealed distributions broader than expected from shot noise, indicating conformational heterogeneity. Across the TXBX series, control constructs showed the lowest variance, indicating greater rigidity compared with damaged constructs. Ribo and 8–oxoG modifications introduced modest flexibility, whereas abasic, nicked, and gapped sites produced the greatest variance, reflecting significant destabilisation and loss of structural rigidity.

## Discussion

This study presents compelling experimental evidence that various types of DNA damage induce measurable, long-range alterations to the overall structure and dynamics of DNA duplexes. Our findings, derived from extensive smFRET measurements, robust statistical analyses, and detailed conformational modelling, provide a novel perspective on how DNA itself might actively participate in the initial steps of damage recognition. The observed changes extend beyond the immediate vicinity of the damaged base, influencing distal regions of the duplex, and vary depending on the nature of the damage.

### Differential Structural Impacts of DNA Damage Types

A key finding of this study was the differential impact of various damage types on DNA conformation. Backbone discontinuities, such as nicks and gaps, consistently induced the most pronounced and widespread structural deviations. This behaviour is consistent with previous theoretical and experimental work suggesting that such breaks increase local flexibility and introduce kinks that may act as physical signals for repair machinery^87^. The observed large breadth of the Gaussian distributions for these constructs, together with the BVA analysis indicate greater conformational heterogeneity and dynamic behaviour relative to undamaged DNA.

In contrast, base and sugar modifications such as 8–oxoG, ribonucleotides, and abasic sites caused more subtle, yet still statistically significant, shifts in FRET efficiency and dye positions. Although these lesions do not involve a break in the backbone, they introduce local distortions through effects on base pairing, stacking, or sugar pucker. The abasic site, for instance, creates a void in the helix, and we observed that this consistently brought dyes closer to the backbone, suggesting local compaction^88^. Similarly, the 8–oxoG lesion, a common oxidative adduct, tends to adopt a syn-conformation, disrupting stacking and enabling mispairing^89^. Our data show that 8–oxoG induces changes in FRET that are more widespread than those caused by ribonucleotides, although less dramatic than those from backbone breaks. Ribonucleotide incorporation, despite involving only a single oxygen atom change, still resulted in measurable conformational shifts, particularly at specific labelling positions. These observations highlight the exquisite sensitivity of smFRET in detecting even minor changes to biomolecular structure.

Across all constructs, nicks and gaps consistently induced pronounced bending of the DNA helix. These lesions generated the largest deviations from the ideal B–form reference, with noticeable kinks near the damage site and more gradual distortions propagating along the duplex. In contrast, ribonucleotide, 8–oxoG, and abasic lesions produced subtler structural effects, primarily manifesting as local unwinding or alterations in helical twist. Ribonucleotides frequently introduced changes reflecting slight helical unwinding and base-stacking disruptions, while abasic sites caused modest kinking and destabilisation of the local helix consistent with the literature^90, 91^. 8–oxoG damage led to intermediate perturbations, combining minor bending with small changes in twist.

NMR spectroscopy has been used to study the structural impact of various DNA lesions. These studies showed how specific damage types alter DNA conformation, particularly in terms of flexibility and local unwinding^92, 93^. Structural studies using NMR spectroscopy further support these interpretations, showing that lesions such as 8–oxoG and abasic sites destabilise the helix, alter base stacking, and increase backbone flexibility^94, 95^. These perturbations align with the structural changes observed in our experiments, indicating how local DNA damage alters duplex conformation along the helix^96^.

### Long-Range Propagation of Conformational Changes

The propagation of these conformational changes to distal sites is particularly noteworthy. Our results demonstrate that the influence of damage is not strictly localised to the immediate vicinity of the lesion but can propagate along the helical axis. This long-range effect suggests a mechanism by which repair proteins might detect damage without necessitating the performance of a precise, base-by-base search along the entire DNA molecule. Instead, they could sense a broader region of altered flexibility or geometry, thereby significantly reducing their search space. This concept aligns with proposed mechanisms of DNA damage sensing, where proteins might scan DNA through non-specific interactions, detecting global changes in DNA architecture before engaging in sequence-specific recognition. For instance, the increased flexibility around a lesion could lead to altered diffusion rates of proteins on the DNA or present a more open or accessible conformation that facilitates binding.

### Implications for DNA Damage Recognition and Repair

The implications of these findings for DNA damage recognition and repair are profound. In the context of the BER, where enzymes like DNA glycosylases must efficiently locate and excise damaged bases, the observed conformational changes could serve as critical pre-recognition signals. Instead of relying solely on direct chemical recognition of the damaged base, which might be a slow and energy-intensive process, BER enzymes could first identify regions of altered DNA flexibility or geometry. For example, the heightened flexibility and distorted conformation induced by nicks and gaps could act as universal flags for repair enzymes that are responsible for processing these crucial intermediates. This structural signalling mechanism could significantly enhance the efficiency of the initial damage detection step, ensure rapid repair and minimise the accumulation of mutagenic lesions.

Furthermore, the differential conformational responses to distinct damage types suggest that different lesions might elicit unique structural signatures, potentially allowing for a refined level of damage discrimination by repair pathways. While our study focuses on the DNA component, it is plausible that these intrinsic DNA conformational changes can influence the binding kinetics and mechanisms of DNA-binding proteins, guiding them to the site of damage. Future studies could investigate how these observed DNA structural changes translate into altered protein-DNA interaction dynamics and specificity.

## Conclusion

This work provides experimental evidence that various types of DNA damage, ranging from a single oxygen atom addition (ribonucleotide vs. deoxyribonucleotide) to larger structural changes (nick and gap), result in measurable alterations to the overall structure and dynamics of DNA duplexes. We observed that different types of damage not only impact the local flexibility of DNA but also influence regions distal to the damage site. Our experimental design and the developed analysis workflow, together with the sensitivity of single-molecule FRET, enabled us to measure the changes in structure and flexibility arising from the various types of damage tested.

Notably, nick and gap modifications induced the most significant alterations in dye positions, affecting both their proximity to the DNA backbone and their distance from the damage site. This suggests that these disruptions create the most pronounced structural deviations. Conversely, 8–oxoG, ribonucleotide, and abasic sites caused more subtle changes, indicating that these damages, while detectable, may rely on additional factors for efficient protein recruitment.

These findings provide a deeper understanding of how damaged DNA substrates involved in the BER pathway may serve as signals for repair proteins. The conformational changes observed suggest that DNA damage can modulate the physical properties of the DNA itself, creating a readout of structural changes that could facilitate recognition by BER enzymes. This work supports the hypothesis that damage detection could be initiated within the DNA molecule itself, before direct binding by repair proteins, offering new insights into the initial steps of BER and the role of structural dynamics in damage recognition.

## ACKNOWLEDGEMENTS

Thank you to Dr Steven Quinn for allowing us time on the FluoTime 300 (PicoQuant) spectrophotometer to conduct lifetime and anisotropy measurements.

## AUTHOR CONTRIBUTIONS

Sophie E. Fountain: Conceptualisation, Experiments, Analysis, Validation, Writing-original draft. Mahmoud A. S. Abdelhamid: Conceptualisation, Analysis, Validation, Writing-review & editing. Timothy D. Craggs: Conceptualisation, Writing-review & editing.

## SUPPLEMENTARY DATA

Supplementary Data are available at NAR online.

## FUNDING

BBSRC New Investigator Grant: BB/T008032/1 (T.D.C)

BBSRC White Rose DTP Studentship: 2741120 (to S.E.F and T.D.C)

## DATA AVAILABILITY

Raw HDF5 files and analysis Jupyter notebooks stored here: https://doi.org/10.5281/zenodo.17508550, https://doi.org/10.5281/zenodo.17590548, https://doi.org/10.5281/zenodo.18164520, https://doi.org/10.5281/zenodo.18164375 and https://doi.org/10.5281/zenodo.18670016

