## Supplementary data for "DNA damage induces long range changes to duplex structure - a non-protein start to damage detection?"

| **Supplementary material** | **Location** |
|---|---|
| Supplementary Table 1: Damage DNA sequences | p2 |
| Supplementary Table 2: NMS DNA sequences | p3 |
| Supplementary Table 3: Annealing Combinations | P4 |
| Supplementary Table 4-6: Quantum Yield | p5-8 |
| Supplementary Table 7: J-integral | P9 |
| Supplementary Table 8-10: Predicted E | p10 |
| Supplementary Table 11: Lifetime | p11 |
| Supplementary Table 12: Anisotropy | p12 |
| Supplementary Table 13: Summary of quantum yield, J-integral and anisotropy | p13 |
| Supplementary Table 14: Distance Changes in AV mean position | p14-15 |

| **Supplementary material** | **Location** |
|---|---|
| Supplementary Figure 1-9: 2D Histograms | p16-33 |
| Supplementary Figure 10-18: QQ plots, Kolmogorov-Smirnov and Kruskal-Wallis | p34-42 |
| Supplementary Figure 19: Dunn Tests | p43 |
| Supplementary Figure 20-28: QQ plots, Kolmogorov-Smirnov and Kruskal-Wallis | p44-52 |
| Supplementary Figure 29-30: Dunn Tests | p53-54 |
| Supplementary Figure 31-32: Box Plots | p55-56 |
| Supplementary Figure 33: Quantum Yield example calculation | p57 |
| Supplementary Figure 34: Lifetimes | p58-60 |
| Supplementary Figure 35-36: Anisotropies | p60-62 |
| Supplementary Figure 37-38: Dye movement figure | p63-64 |
| Supplementary Figure 39-47: BVA plots | p65-82 |

Supplementary Table 1: "T" strands were labelled with the donor dye ATTO 550 and "B" strands were labelled with the acceptor dye ATTO 647N. The labelling sites of the donor and acceptor are shown in orange and in purple on the sequence respectively. The damage site is labelled in blue.

| **Name** | **Sequence** |
|---|---|
| Top -7 | 5′-CCTCATTCTTCGTCCCATTACCATACATCCCTAGAGAGTAGAGCCTGCTTCGTGG-3′ |
| Top -12 | 5′-CCTCATTCTTCGTCCCATTACCATACATCCCTAGAGAGTAGAGCCTGCTTCGTGG-3′ |
| Top -18 | 5′-CCTCATTCTTCGTCCCATTACCATACATCCCTAGAGAGTAGAGCCTGCTTCGTGG-3′ |
| Bottom +6 | 5′-CCACGAAGCAGGCTCTACTCTCTAGGGATGTATGGTAATGGGACGAAGAATGAGG-3′ |
| Bottom +9 | 5′-CCACGAAGCAGGCTCTACTCTCTAGGGATGTATGGTAATGGGACGAAGAATGAGG-3′ |
| Bottom +11 | 5′-CCACGAAGCAGGCTCTACTCTCTAGGGATGTATGGTAATGGGACGAAGAATGAGG-3′ |
| Half Top -7 | 5′-CCTCATTCTTCGTCCCATTACCATACATCC-3′ |
| Half Top -12 | 5′-CCTCATTCTTCGTCCCATTACCATACATCC-3′ |
| Half Top -18 | 5′-CCTCATTCTTCGTCCCATTACCATACATCC-3′ |
| Bottom +6 (Ribo, 8-oxoG, abasic) | 5′-CCACGAAGCAGGCTCTACTCTCTAGGGATGTATGGTAATGGGACGAAGAATGAGG-3′ |
| Bottom +9 (Ribo, 8-oxoG, abasic) | 5′-CCACGAAGCAGGCTCTACTCTCTAGGGATGTATGGTAATGGGACGAAGAATGAGG-3′ |
| Bottom +11 (Ribo, 8-oxoG, abasic) | 5′-CCACGAAGCAGGCTCTACTCTCTAGGGATGTATGGTAATGGGACGAAGAATGAGG-3′ |
| Half Top (Nick) | 5′-CTAGAGAGTAGAGCCTGCTTCGTGG-3′ |
| Half Top (Gap) | 5′-TAGAGAGTAGAGCCTGCTTCGTGG-3′ |

Supplementary Table 2: NMS1,2,3 strands were labelled with the donor dye ATTO 550 and NMS4 strands were labelled with the acceptor dye ATTO 647N. The labelling sites of the donor and acceptor are shown in orange and in purple on the sequence respectively. The damage site is labelled in blue.

| Name | Sequence |
|---|---|
| NMS1 | 5′-GAGCTGAAAGTGTCGAGTTTGTTTGAGTGTTTGTCTGG-3′ |
| NMS2 | 5′-GAGCTGAAAGTGTCGAGTTTGTTTGAGTGTTTGTCTGG-3′ |
| NMS3 | 5′-GAGCTGAAAGTGTCGAGTTTGTTTGAGTGTTTGTCTGG-3′ |
| NMS4 | 5′-CCAGACAAACACTCAAACAAACTCGACACTTTCAGCTC-3′ |
| NMS4 (Ribo, 8-oxoG, abasic) | 5′- CCAGACAAACACTCAAACAAACTCGACACTTTCAGCTC -3′ |

Supplementary Table 3: Duplex constructs from combinations of constituent oligonucleotides where (X) indicates the respective labelling position from Table S1 and 2 above.

| **Construct** | **Donor Dye** | **Acceptor Dye** |
|---|---|---|
| Full length duplex (Control) | Top(X) | Bottom(X) |
| Swapping of deoxyribose to ribose sugar (Ribo) | | Bottom(X) (Ribo) |
| Addition of oxygen on guanine base (8-oxoG) | | Bottom(X) (8-oxoG) |
| Removal of base (Abasic) | | Bottom(X) (Abasic) |
| Break in the backbone (Nick) | Half Top(X) + Half Top (Nick) | Bottom(X) |
| Removal of a nucleotide (Gap) | Half Top(X) + Half Top (Gap) | Bottom(X) |

| **Construct** | **Donor Dye** | **Acceptor Dye** |
|---|---|---|
| Full length duplex (Control) | NMS1,2,3 (X) | NMS4(X) |
| Swapping of deoxyribose to ribose sugar (Ribo) | | NMS4 (X) (Ribo) |
| Addition of oxygen on guanine base (8-oxoG) | | NMS4 (X) (8-oxoG) |
| Removal of base (Abasic) | | NMS4 (X) (Abasic) |

Supplementary Table 4: Quantum Yield measurements showing the excitation wavelength (Ex), reference sample (Ref.), quantum yield of reference sample (Qr) and refractive index of reference sample (nr) and measured samples' quantum yield (Qs) and refractive index (ns) with confidence values ($R^2$).

| Ex | Ref. | Qr | nr | $R^2$(Ref.) | Unknown | Qs | ns | $R^2$ (Unk.) |
|---|---|---|---|---|---|---|---|---|
| **550** | RDB - T1 | 0.7 | 1.33 | 0.999907305 | ATTO 550 - T1 | 0.804132455 | 1.33 | 0.999764886 |
| **550** | ATTO 550 - T3 | 0.8 | 1.33 | 0.999732951 | ATTO 550 - T2 | 0.795370245 | 1.33 | 0.999556273 |
| **550** | NMS1 - T2 | 0.76 | 1.4 | 0.995850854 | ATTO 550 - T2 | 0.794693752 | 1.33 | 0.999556273 |
| **550** | NMS2 - T2 | 0.77 | 1.4 | 0.998037073 | ATTO 550 - T2 | 0.783686177 | 1.33 | 0.999556273 |
| **550** | NMS3 - T2 | 0.76 | 1.4 | 0.998092052 | ATTO 550 - T2 | 0.803539565 | 1.33 | 0.999556273 |
| **550** | RDB - T2 | 0.7 | 1.33 | 0.991981701 | ATTO 550 - T2 | 0.837038937 | 1.33 | 0.999556273 |
| **550** | RDB - T3 | 0.7 | 1.33 | 0.999678604 | ATTO 550 - T2 | 0.902796585 | 1.33 | 0.999556273 |
| **550** | ATTO 550 - T2 | 0.8 | 1.33 | 0.999556273 | ATTO 550 - T3 | 0.804656705 | 1.33 | 0.999732951 |
| **550** | NMS1 - T2 | 0.76 | 1.4 | 0.995850854 | ATTO 550 - T3 | 0.79931957 | 1.33 | 0.999732951 |
| **550** | NMS2 - T2 | 0.77 | 1.4 | 0.998037073 | ATTO 550 - T3 | 0.788247921 | 1.33 | 0.999732951 |
| **550** | NMS3 - T2 | 0.76 | 1.4 | 0.998092052 | ATTO 550 - T3 | 0.808216873 | 1.33 | 0.999732951 |
| **550** | RDB - T2 | 0.7 | 1.33 | 0.991981701 | ATTO 550 - T3 | 0.841911241 | 1.33 | 0.999732951 |
| **550** | RDB - T3 | 0.7 | 1.33 | 0.999678604 | ATTO 550 - T3 | 0.908051656 | 1.33 | 0.999732951 |
| **550** | NMS2 - T1 | 0.77 | 1.4 | 0.999351269 | NMS1 - T1 | 0.811843308 | 1.4 | 0.998396555 |
| **550** | NMS3 - T1 | 0.76 | 1.4 | 0.999473103 | NMS1 - T1 | 0.8197872 | 1.4 | 0.998396555 |
| **550** | ATTO 550 - T2 | 0.8 | 1.33 | 0.999556273 | NMS1 - T2 | 0.765074595 | 1.4 | 0.995850854 |
| **550** | ATTO 550 - T3 | 0.8 | 1.33 | 0.999732951 | NMS1 - T2 | 0.760646959 | 1.4 | 0.995850854 |
| **550** | NMS2 - T2 | 0.77 | 1.4 | 0.998037073 | NMS1 - T2 | 0.74947298 | 1.4 | 0.995850854 |
| **550** | NMS3 - T2 | 0.76 | 1.4 | 0.998092052 | NMS1 - T2 | 0.768459634 | 1.4 | 0.995850854 |
| **550** | RDB - T2 | 0.7 | 1.33 | 0.991981701 | NMS1 - T2 | 0.800496532 | 1.4 | 0.995850854 |
| **550** | RDB - T3 | 0.7 | 1.33 | 0.999678604 | NMS1 - T2 | 0.863383414 | 1.4 | 0.995850854 |
| **550** | NMS2 - T3 | 0.77 | 1.4 | 0.998459128 | NMS1 - T3 | 0.738493975 | 1.4 | 0.996338118 |
| **550** | NMS3 - T3 | 0.76 | 1.4 | 0.999717452 | NMS1 - T3 | 0.767643962 | 1.4 | 0.996338118 |
| **550** | NMS1 - T1 | 0.76 | 1.4 | 0.998396555 | NMS2 - T1 | 0.720828754 | 1.4 | 0.999351269 |
| **550** | NMS3 - T1 | 0.76 | 1.4 | 0.999473103 | NMS2 - T1 | 0.777534455 | 1.4 | 0.999351269 |
| **550** | ATTO 550 - T2 | 0.8 | 1.33 | 0.999556273 | NMS2 - T2 | 0.786028921 | 1.4 | 0.998037073 |
| **550** | ATTO 550 - T3 | 0.8 | 1.33 | 0.999732951 | NMS2 - T2 | 0.781480019 | 1.4 | 0.998037073 |
| **550** | NMS1 - T2 | 0.76 | 1.4 | 0.995850854 | NMS2 - T2 | 0.78081534 | 1.4 | 0.998037073 |
| **550** | NMS3 - T2 | 0.76 | 1.4 | 0.998092052 | NMS2 - T2 | 0.789506671 | 1.4 | 0.998037073 |
| **550** | RDB - T2 | 0.7 | 1.33 | 0.991981701 | NMS2 - T2 | 0.822421016 | 1.4 | 0.998037073 |
| **550** | RDB - T3 | 0.7 | 1.33 | 0.999678604 | NMS2 - T2 | 0.887030282 | 1.4 | 0.998037073 |
| **550** | NMS1 - T3 | 0.76 | 1.4 | 0.996338118 | NMS2 - T3 | 0.792423527 | 1.4 | 0.998459128 |
| **550** | NMS3 - T3 | 0.76 | 1.4 | 0.999717452 | NMS2 - T3 | 0.8003936 | 1.4 | 0.998459128 |
| **550** | NMS1 - T1 | 0.76 | 1.4 | 0.998396555 | NMS3 - T1 | 0.704573089 | 1.4 | 0.999473103 |
| **550** | NMS2 - T1 | 0.77 | 1.4 | 0.999351269 | NMS3 - T1 | 0.752635457 | 1.4 | 0.999473103 |
| **550** | ATTO 550 - T2 | 0.8 | 1.33 | 0.999556273 | NMS3 - T2 | 0.756652225 | 1.4 | 0.998092052 |
| **550** | ATTO 550 - T3 | 0.8 | 1.33 | 0.999732951 | NMS3 - T2 | 0.752273332 | 1.4 | 0.998092052 |
| **550** | NMS1 - T2 | 0.76 | 1.4 | 0.995850854 | NMS3 - T2 | 0.751633495 | 1.4 | 0.998092052 |
| **550** | NMS2 - T2 | 0.77 | 1.4 | 0.998037073 | NMS3 - T2 | 0.741222362 | 1.4 | 0.998092052 |
| **550** | RDB - T2 | 0.7 | 1.33 | 0.991981701 | NMS3 - T2 | 0.791684218 | 1.4 | 0.998092052 |
| **550** | RDB - T3 | 0.7 | 1.33 | 0.999678604 | NMS3 - T2 | 0.853878806 | 1.4 | 0.998092052 |
| **550** | NMS1 - T3 | 0.76 | 1.4 | 0.996338118 | NMS3 - T3 | 0.752432154 | 1.4 | 0.999717452 |
| **550** | NMS2 - T3 | 0.77 | 1.4 | 0.998459128 | NMS3 - T3 | 0.731140279 | 1.4 | 0.999717452 |
| **550** | NMS1 - T1 | 0.76 | 1.4 | 0.998396555 | Top -12 - T1 | 0.687232066 | 1.4 | 0.997708725 |
| **550** | NMS2 - T1 | 0.77 | 1.4 | 0.999351269 | Top -12 - T1 | 0.734111518 | 1.4 | 0.997708725 |
| **550** | NMS3 - T1 | 0.76 | 1.4 | 0.999473103 | Top -12 - T1 | 0.741294804 | 1.4 | 0.997708725 |
| **550** | ATTO 550 - T2 | 0.8 | 1.33 | 0.999556273 | Top -12 - T2 | 0.744499768 | 1.4 | 0.996475725 |
| **550** | ATTO 550 - T3 | 0.8 | 1.33 | 0.999732951 | Top -12 - T2 | 0.740191203 | 1.4 | 0.996475725 |
| **550** | NMS1 - T2 | 0.76 | 1.4 | 0.995850854 | Top -12 - T2 | 0.739561642 | 1.4 | 0.996475725 |
| **550** | NMS2 - T2 | 0.77 | 1.4 | 0.998037073 | Top -12 - T2 | 0.729317721 | 1.4 | 0.996475725 |
| **550** | NMS3 - T2 | 0.76 | 1.4 | 0.998092052 | Top -12 - T2 | 0.747793775 | 1.4 | 0.996475725 |
| **550** | RDB - T2 | 0.7 | 1.33 | 0.991981701 | Top -12 - T2 | 0.778969118 | 1.4 | 0.996475725 |
| **550** | RDB - T3 | 0.7 | 1.33 | 0.999678604 | Top -12 - T2 | 0.84016481 | 1.4 | 0.996475725 |
| **550** | NMS1 - T3 | 0.76 | 1.4 | 0.996338118 | Top -12 - T3 | 0.74359809 | 1.4 | 0.99386293 |

| | | | | | | | | |
|---|---|---|---|---|---|---|---|---|
| **550** | NMS2 - T3 | 0.77 | 1.4 | 0.998459128 | Top -12 - T3 | 0.722556196 | 1.4 | 0.99386293 |
| **550** | NMS3 - T3 | 0.76 | 1.4 | 0.999717452 | Top -12 - T3 | 0.751077084 | 1.4 | 0.99386293 |
| **550** | NMS1 - T1 | 0.76 | 1.4 | 0.998396555 | Top -18 - T1 | 0.707095204 | 1.4 | 0.999083843 |
| **550** | NMS2 - T1 | 0.77 | 1.4 | 0.999351269 | Top -18 - T1 | 0.755329618 | 1.4 | 0.999083843 |
| **550** | NMS3 - T1 | 0.76 | 1.4 | 0.999473103 | Top -18 - T1 | 0.762720523 | 1.4 | 0.999083843 |
| **550** | ATTO 550 - T2 | 0.8 | 1.33 | 0.999556273 | Top -18 - T2 | 0.728682996 | 1.4 | 0.999414902 |
| **550** | ATTO 550 - T3 | 0.8 | 1.33 | 0.999732951 | Top -18 - T2 | 0.724465966 | 1.4 | 0.999414902 |
| **550** | NMS1 - T2 | 0.76 | 1.4 | 0.995850854 | Top -18 - T2 | 0.72384978 | 1.4 | 0.999414902 |
| **550** | NMS2 - T2 | 0.77 | 1.4 | 0.998037073 | Top -18 - T2 | 0.713823489 | 1.4 | 0.999414902 |
| **550** | NMS3 - T2 | 0.76 | 1.4 | 0.998092052 | Top -18 - T2 | 0.731907022 | 1.4 | 0.999414902 |
| **550** | RDB - T2 | 0.7 | 1.33 | 0.991981701 | Top -18 - T2 | 0.762420051 | 1.4 | 0.999414902 |
| **550** | RDB - T3 | 0.7 | 1.33 | 0.999678604 | Top -18 - T2 | 0.82231565 | 1.4 | 0.999414902 |
| **550** | NMS1 - T3 | 0.76 | 1.4 | 0.996338118 | Top -18 - T3 | 0.730820344 | 1.4 | 0.998974109 |
| **550** | NMS2 - T3 | 0.77 | 1.4 | 0.998459128 | Top -18 - T3 | 0.710140027 | 1.4 | 0.998974109 |
| **550** | NMS3 - T3 | 0.76 | 1.4 | 0.999717452 | Top -18 - T3 | 0.738170821 | 1.4 | 0.998974109 |
| **550** | NMS1 - T1 | 0.76 | 1.4 | 0.998396555 | Top -7 - T1 | 0.691377728 | 1.4 | 0.999062676 |
| **550** | NMS2 - T1 | 0.77 | 1.4 | 0.999351269 | Top -7 - T1 | 0.738539977 | 1.4 | 0.999062676 |
| **550** | NMS3 - T1 | 0.76 | 1.4 | 0.999473103 | Top -7 - T1 | 0.745766595 | 1.4 | 0.999062676 |
| **550** | ATTO 550 - T2 | 0.8 | 1.33 | 0.999556273 | Top -7 - T2 | 0.723404389 | 1.4 | 0.998304516 |
| **550** | ATTO 550 - T3 | 0.8 | 1.33 | 0.999732951 | Top -7 - T2 | 0.719217907 | 1.4 | 0.998304516 |
| **550** | NMS1 - T2 | 0.76 | 1.4 | 0.995850854 | Top -7 - T2 | 0.718606185 | 1.4 | 0.998304516 |
| **550** | NMS2 - T2 | 0.77 | 1.4 | 0.998037073 | Top -7 - T2 | 0.708652525 | 1.4 | 0.998304516 |
| **550** | NMS3 - T2 | 0.76 | 1.4 | 0.998092052 | Top -7 - T2 | 0.72660506 | 1.4 | 0.998304516 |
| **550** | RDB - T2 | 0.7 | 1.33 | 0.991981701 | Top -7 - T2 | 0.756897051 | 1.4 | 0.998304516 |
| **550** | RDB - T3 | 0.7 | 1.33 | 0.999678604 | Top -7 - T2 | 0.816358765 | 1.4 | 0.998304516 |
| **550** | NMS1 - T3 | 0.76 | 1.4 | 0.996338118 | Top -7 - T3 | 0.733526535 | 1.4 | 0.999315897 |
| **550** | NMS2 - T3 | 0.77 | 1.4 | 0.998459128 | Top -7 - T3 | 0.71276964 | 1.4 | 0.999315897 |
| **550** | NMS3 - T3 | 0.76 | 1.4 | 0.999717452 | Top -7 - T3 | 0.740904231 | 1.4 | 0.999315897 |

Supplementary Table 5: Quantum Yield measurements showing the excitation wavelength (Ex), reference sample (Ref.), quantum yield of reference sample (Qr) and refractive index of reference sample (nr) and measured samples' quantum yield (Qs) and refractive index (ns) with confidence values ($R^2$).

| Ex | Ref. | Qr | Nr | $R^2$ (Ref.) | Unk. | Qs | Ns | $R^2$ (Unk.) |
|---|---|---|---|---|---|---|---|---|
| **646** | ATTO 647N - T1 | 0.65 | 1.33 | 0.999999666 | ATTO 647N - T2 | 0.637195275 | 1.33 | 0.99884106 |
| **646** | ATTO 647N - T1 | 0.65 | 1.33 | 0.999999666 | ATTO 647N - T3 | 0.658075988 | 1.33 | 0.99959802 |
| **646** | ATTO 647N - T1 | 0.65 | 1.33 | 0.999999666 | Bottom +11 - T1 | 0.539973545 | 1.4 | 0.999931515 |
| **646** | ATTO 647N - T1 | 0.65 | 1.33 | 0.996215287 | Bottom +11 - T3 | 0.50486727 | 1.4 | 0.999231646 |
| **646** | ATTO 647N - T1 | 0.65 | 1.33 | 0.999999666 | Bottom +6 - T1 | 0.552265222 | 1.4 | 0.999339418 |
| **646** | ATTO 647N - T1 | 0.65 | 1.33 | 0.996215287 | Bottom +6 - T3 | 0.492425515 | 1.4 | 0.997963665 |
| **646** | ATTO 647N - T1 | 0.65 | 1.33 | 0.999999666 | Bottom +9 - T1 | 0.528427847 | 1.4 | 0.998569767 |
| **646** | ATTO 647N - T1 | 0.65 | 1.33 | 0.996215287 | Bottom +9 - T3 | 0.513297417 | 1.4 | 0.999291052 |
| **646** | ATTO 647N - T1 | 0.65 | 1.33 | 0.999999666 | NMS4 - T1 | 0.57295812 | 1.4 | 0.999913668 |
| **646** | ATTO 647N - T1 | 0.65 | 1.33 | 0.996215287 | NMS4 - T3 | 0.538354994 | 1.4 | 0.99878214 |
| **646** | ATTO 647N - T2 | 0.65 | 1.33 | 0.99884106 | ATTO 647N - T1 | 0.663062042 | 1.33 | 0.999999666 |
| **646** | ATTO 647N - T2 | 0.65 | 1.33 | 0.99884106 | ATTO 647N - T3 | 0.67130032 | 1.33 | 0.99959802 |
| **646** | ATTO 647N - T2 | 0.65 | 1.33 | 0.999421881 | Bottom +11 - T2 | 0.532648855 | 1.4 | 0.999886686 |
| **646** | ATTO 647N - T2 | 0.65 | 1.33 | 0.99884106 | Bottom +11 - T1 | 0.550824556 | 1.4 | 0.999931515 |
| **646** | ATTO 647N - T2 | 0.65 | 1.33 | 0.99884106 | Bottom +6 - T1 | 0.56336324 | 1.4 | 0.999339418 |
| **646** | ATTO 647N - T2 | 0.65 | 1.33 | 0.999421881 | Bottom +6 - T2 | 0.541131398 | 1.4 | 0.999553447 |
| **646** | ATTO 647N - T2 | 0.65 | 1.33 | 0.999421881 | Bottom +9 - T2 | 0.530704257 | 1.4 | 0.99903735 |
| **646** | ATTO 647N - T2 | 0.65 | 1.33 | 0.99884106 | Bottom +9 - T1 | 0.539046842 | 1.4 | 0.998569767 |
| **646** | ATTO 647N - T2 | 0.65 | 1.33 | 0.999421881 | NMS4 - T2 | 0.556320655 | 1.4 | 0.999484075 |
| **646** | ATTO 647N - T2 | 0.65 | 1.33 | 0.99884106 | NMS4 - T1 | 0.58447197 | 1.4 | 0.999913668 |
| **646** | ATTO 647N - T3 | 0.65 | 1.33 | 0.99959802 | ATTO 647N - T1 | 0.642023121 | 1.33 | 0.999999666 |
| **646** | ATTO 647N - T3 | 0.65 | 1.33 | 0.99959802 | ATTO 647N - T2 | 0.629375538 | 1.33 | 0.99884106 |
| **646** | ATTO 647N - T3 | 0.65 | 1.33 | 0.99959802 | Bottom +11 - T1 | 0.533346925 | 1.4 | 0.999931515 |
| **646** | ATTO 647N - T3 | 0.65 | 1.33 | 0.99959802 | Bottom +6 - T1 | 0.545487757 | 1.4 | 0.999339418 |
| **646** | ATTO 647N - T3 | 0.65 | 1.33 | 0.99959802 | Bottom +9 - T1 | 0.521942917 | 1.4 | 0.998569767 |
| **646** | ATTO 647N - T3 | 0.65 | 1.33 | 0.99959802 | NMS4 - T1 | 0.565926708 | 1.4 | 0.999913668 |

Supplementary Table 6: Quantum Yield values compared to literature values.

| Construct | Qs | Literature value |
|---|---|---|
| **ATTO 550** | 0.82±0.04 | 0.8 |
| **NMS 1** | 0.78±0.04 | 0.76±0.015 |
| **NMS 2** | 0.79±0.04 | 0.77±0.015 |
| **NMS 3** | 0.76±0.04 | 0.76±0.015 |
| **Top -7** | 0.73±0.03 | - |
| **Top -12** | 0.75±0.03 | - |
| **Top -18** | 0.74±0.03 | - |
| **ATTO 647N** | 0.65±0.02 | 0.65 |
| **NMS 4** | 0.57±0.02 | - |
| **Bottom +6** | 0.54±0.03 | - |
| **Bottom +9** | 0.53±0.01 | - |
| **Bottom +11** | 0.53±0.02 | - |

Supplementary Table 7: Table showing Epsilon, quantum yield of donor (Qd), refractive index (n), Kappa squared ($K^2$), J Integral (J) and Forster radius ($R_0$).

| **Donor** | **Acceptor** | **Epsilon** | **Qd** | **n** | **$K^2$** | **J** | **$R_0$** |
|---|---|---|---|---|---|---|---|
| ATTO 550 | ATTO 647N | 150000 | 0.82 | 1.33 | 0.66666667 | 3.70E+15 | 62.0 |
| NMS1 | NMS4 | 150000 | 0.78 | 1.4 | 0.66666667 | 4.11E+15 | 60.4 |
| NMS2 | NMS4 | 150000 | 0.79 | 1.4 | 0.66666667 | 4.09E+15 | 60.5 |
| NMS3 | NMS4 | 150000 | 0.76 | 1.4 | 0.66666667 | 4.11E+15 | 60.2 |
| Top -7 | Bottom +6 | 150000 | 0.73 | 1.4 | 0.66666667 | 4.15E+15 | 59.9 |
| Top -7 | Bottom +9 | 150000 | 0.73 | 1.4 | 0.66666667 | 3.95E+15 | 59.4 |
| Top -7 | Bottom +11 | 150000 | 0.73 | 1.4 | 0.66666667 | 4.45E+15 | 60.6 |
| Top -12 | Bottom +6 | 150000 | 0.75 | 1.4 | 0.66666667 | 4.14E+15 | 60.1 |
| Top -12 | Bottom +9 | 150000 | 0.75 | 1.4 | 0.66666667 | 3.94E+15 | 59.6 |
| Top -12 | Bottom +11 | 150000 | 0.75 | 1.4 | 0.66666667 | 4.44E+15 | 60.8 |
| Top -18 | Bottom +6 | 150000 | 0.74 | 1.4 | 0.66666667 | 4.09E+15 | 59.9 |
| Top -18 | Bottom +9 | 150000 | 0.74 | 1.4 | 0.66666667 | 3.89E+15 | 59.4 |
| Top -18 | Bottom +11 | 150000 | 0.74 | 1.4 | 0.66666667 | 4.39E+15 | 60.6 |

Supplementary Table 8: Dimensions of the linkers with different dyes.

| | Linker Length [Å] | Linker Width [Å] | $R_1$ [Å] | $R_2$ [Å] | $R_3$ [Å] |
|---|---|---|---|---|---|
| dT-C6-ATTO 550 | 16.43 | 5.9 | 8.6 | 5.37 | 4.62 |
| dT-C6-ATTO 647N | 15.02 | 4.49 | 8.62 | 5.45 | 5.02 |

Supplementary Table 9: Predicted FRET efficiencies and distances for control duplex with different labelling positions.

| | Top -7 Bottom +6 | Top -7 Bottom +9 | Top -7 Bottom +11 | Top -12 Bottom +6 | Top -12 Bottom +9 | Top -12 Bottom +11 | Top -18 Bottom +6 | Top -18 Bottom +9 | Top -18 Bottom +11 |
|---|---|---|---|---|---|---|---|---|---|
| Predicted FRET efficiencies | 0.96 | 0.81 | 0.65 | 0.65 | 0.52 | 0.45 | 0.39 | 0.19 | 0.12 |
| Predicted Mean Distance [Å] | 35.13 | 47.45 | 56.12 | 56.12 | 56.12 | 64.84 | 68.20 | 80.60 | 87.36 |

Supplementary Table 10: Differences in predicted and measured mean FRET efficiencies.

| | Top -7 Bottom +6 | Top -7 Bottom +9 | Top -7 Bottom +11 | Top -12 Bottom +6 | Top -12 Bottom +9 | Top -12 Bottom +11 | Top -18 Bottom +6 | Top -18 Bottom +9 | Top -18 Bottom +11 |
|---|---|---|---|---|---|---|---|---|---|
| Förster radius | 59.9 | 59.4 | 60.6 | 60.1 | 59.6 | 60.8 | 59.9 | 59.4 | 60.6 |
| Predicted Mean FRET efficiencies | 0.95 | 0.77 | 0.61 | 0.60 | 0.45 | 0.41 | 0.33 | 0.15 | 0.11 |
| Measured Mean FRET efficiencies | 0.60 | 0.49 | 0.39 | 0.38 | 0.27 | 0.25 | 0.22 | 0.16 | 0.14 |
| % Difference | 45.2 | 44.4 | 44.0 | 44.9 | 50.0 | 48.5 | 40 | 6.5 | 24 |

Supplementary Table 11: Lifetime values with standard deviation.

| | **Lifetime (ns)** | **Sd (ns)** |
|---|---|---|
| ATTO 550 | 3.60 | 0.0033 |
| NMS 1 | 3.73 | 0.0088 |
| NMS 2 | 3.82 | 0 |
| NMS 3 | 3.73 | 0.0033 |
| T-7 | 3.90 | 0 |
| T-12 | 3.81 | 0.0058 |
| T-18 | 3.59 | 0.0058 |
| NMS 1+4 | 3.15 | 0.01 |
| NMS 2+4 | 2.59 | 0.0333 |
| NMS 3+4 | 2.63 | 0.04 |
| T-7 B+6 | 2.08 | 0.023 |
| T-7 B+9 | 2.45 | 0.012 |
| T-7 B+11 | 2.59 | 0.012 |
| T-12 B+6 | 2.40 | 0.017 |
| T-12 B+9 | 2.88 | 0.006 |
| T-12 B+11 | 2.90 | 0.012 |
| T-18 B+6 | 2.86 | 0 |
| T-18 B+9 | 3.18 | 0 |
| T-18 B+11 | 3.25 | 0.006 |
| ATTO 647N | 3.58 | 0.0033 |
| NMS 4 | 4.08 | 0.0033 |
| B+6 | 4.06 | 0.006 |
| B+9 | 4.24 | 0.0033 |
| B+11 | 4.08 | 0.006 |

Supplementary Table 12: Residual anisotropy results.

| | Residual | | | Residual |
|---|---|---|---|---|
| ATTO 550 T1 | 0.007 | | ATTO 647N T1 | 0.019 |
| ATTO 550 T2 | 0.002 | | ATTO 647N T2 | 0.006 |
| ATTO 550 T3 | 0.017 | | ATTO 647N T3 | 0.007 |
| Average | 0.009 | | Average | 0.011 |
| NMS 1 T1 | 0.085 | | NMS 4 T1 | 0.018 |
| NMS 1 T2 | 0.052 | | NMS 4 T2 | 0.035 |
| NMS 1 T3 | 0.081 | | NMS 4 T3 | 0.027 |
| Average | 0.073 | | Average | 0.027 |
| NMS 2 T1 | 0.091 | | Bottom +6 T1 | 0.023 |
| NMS 2 T2 | 0.111 | | Bottom +6 T2 | 0.043 |
| NMS 2 T3 | 0.094 | | Bottom +6 T3 | 0.033 |
| Average | 0.099 | | Average | 0.033 |
| NMS 3 T1 | 0.074 | | Bottom +9 T1 | 0.027 |
| NMS 3 T2 | 0.091 | | Bottom +9 T2 | 0.028 |
| NMS 3 T3 | 0.087 | | Bottom +9 T3 | 0.027 |
| Average | 0.084 | | Average | 0.027 |
| T-7 T1 | 0.144 | | | |
| T-7 T2 | 0.132 | | Bottom +11 T1 | 0.021 |
| T-7 T3 | 0.138 | | Bottom +11 T2 | 0.016 |
| Average | 0.138 | | Bottom +11 T3 | 0.026 |
| | | | Average | 0.021 |
| T-12 T1 | 0.101 | | | |
| T-12 T2 | 0.100 | | | |
| T-12 T3 | 0.105 | | | |
| Average | 0.102 | | | |
| T-18 T1 | 0.064 | | | |
| T-18 T2 | 0.042 | | | |
| T-18 T3 | 0.054 | | | |
| Average | 0.053 | | | |

| Construct | Residual Anisotropy |
|---|---|
| ATTO 550 | 0.009 ± 0.0076 |
| NMS 1 | 0.073 ± 0.018 |
| NMS 2 | 0.099 ± 0.0108 |
| NMS 3 | 0.084 ± 0.0089 |
| T-7 | 0.138 ± 0.006 |
| T-12 | 0.102 ± 0.0026 |
| T-18 | 0.053 ± 0.011 |
| ATTO 647N | 0.011 ± 0.007 |
| NMS 4 | 0.027 ± 0.009 |
| Bottom +6 | 0.033 ± 0.010 |
| Bottom +9 | 0.027 ± 0.001 |
| Bottom +11 | 0.021 ± 0.005 |

Supplementary Table 13: Summary results.

| | Top -7 | Bottom +6 | Top -7 | Bottom +9 | Top -7 | Bottom +11 | Top -12 | Bottom +6 |
|---|---|---|---|---|---|---|---|---|
| Dye Position | T24 (C6), T Strand | T19 (C6), B Strand | T24 (C6), T Strand | T16 (C6), B Strand | T24 (C6), T Strand | T14 (C6), B Strand | T19 (C6), T Strand | T19 (C6), B Strand |
| Residual anisotropy | 0.138 ± 0.006 | 0.033 ± 0.010 | 0.138 ± 0.006 | 0.027 ± 0.001 | 0.138 ± 0.006 | 0.021 ± 0.005 | 0.102 ± 0.0026 | 0.033 ± 0.010 |
| Combined anisotropy | 0.068±0.015 | | 0.061±0.006 | | 0.054±0.015 | | 0.058±0.011 | |
| Lifetime | 3.9±0 | 4.06±0.006 | 3.9±0 | 4.24±0.0033 | 3.9±0 | 4.08±0.006 | 3.81±0.0058 | 4.06±0.006 |
| Fluorescence quantum yield | 0.73±0.03 | 0.54±0.03 | 0.73±0.03 | 0.53±0.01 | 0.73±0.03 | 0.53±0.02 | 0.75±0.03 | 0.54±0.03 |

| Top -12 | Bottom +9 | Top -12 | Bottom +11 | Top -18 | Bottom +6 | Top -18 | Bottom +9 | Top -18 | Bottom +11 |
|---|---|---|---|---|---|---|---|---|---|
| T19 (C6), T Strand | T16 (C6), B Strand | T19 (C6), T Strand | T14 (C6), B Strand | T13 (C6), T Strand | T19 (C6), B Strand | T13 (C6), T Strand | T16 (C6), B Strand | T13 (C6), T Strand | T14 (C6), B Strand |
| 0.102 ± 0.0026 | 0.027 ± 0.001 | 0.102 ± 0.0026 | 0.021 ± 0.005 | 0.053 ± 0.011 | 0.033 ± 0.010 | 0.053 ± 0.011 | 0.027 ± 0.001 | 0.053 ± 0.011 | 0.021 ± 0.005 |
| 0.053±0.009 | | 0.046±0.014 | | 0.039±0.008 | | 0.027±0.005 | | 0.026±0.008 | |
| 3.81±0.0058 | 4.24±0.0033 | 3.81±0.0058 | 4.08±0.006 | 3.59±0.0058 | 4.06±0.006 | 3.59±0.0058 | 4.24±0.0033 | 3.59±0.0058 | 4.08±0.006 |
| 0.75±0.03 | 0.53±0.01 | 0.75±0.03 | 0.53±0.02 | 0.74±0.03 | 0.54±0.03 | 0.74±0.03 | 0.53±0.01 | 0.74±0.03 | 0.53±0.02 |

| | NMS 1 | NMS 4 | NMS 2 | NMS 4 | NMS 3 | NMS 4 |
|---|---|---|---|---|---|---|
| Dye Position | T31 (C6), D-strand | T31 (C6), A-strand | T23 (C6), D-strand | T31 (C6), A-strand | T19 (C6), D-strand | T31 (C6), A-strand |
| Residual anisotropy | 0.073 ± 0.018 | 0.027 ± 0.009 | 0.099 ± 0.0108 | 0.027 ± 0.009 | 0.084 ± 0.0089 | 0.027 ± 0.009 |
| Combined anisotropy | 0.044±0.013 | | 0.052±0.011 | | 0.048±0.010 | |
| Lifetime | 3.73±0.0088 | 4.08±0.0033 | 3.82±0 | 4.08±0.0033 | 3.73±0.0033 | 4.08±0.0033 |
| Fluorescence quantum yield | 0.78±0.04 | 0.57±0.02 | 0.79±0.04 | 0.57±0.02 | 0.76±0.04 | 0.57±0.02 |

Supplementary Table 14: Estimated distances for each labelling position and damage type and distance changes in AV mean position.

| TX | BX | Construct | R (Å) |
|---|---|---|---|
| T7 | B6 | Control | 57.21 |
| T7 | B6 | O6-Me | 54.84 |
| T7 | B6 | Ribo | 55.66 |
| T7 | B6 | 8-oxoG | 56.62 |
| T7 | B6 | abasic | 54.00 |
| T7 | B6 | Nick | 50.68 |
| T7 | B6 | Gap | 48.94 |
| T7 | B9 | Control | 60.80 |
| T7 | B9 | O6-Me | 60.00 |
| T7 | B9 | Ribo | 60.96 |
| T7 | B9 | 8-oxoG | 61.83 |
| T7 | B9 | abasic | 60.12 |
| T7 | B9 | Nick | 56.92 |
| T7 | B9 | Gap | 54.45 |
| T7 | B11 | Control | 64.39 |
| T7 | B11 | O6-Me | 65.29 |
| T7 | B11 | Ribo | 65.07 |
| T7 | B11 | 8-oxoG | 64.68 |
| T7 | B11 | abasic | 62.58 |
| T7 | B11 | Nick | 60.62 |
| T7 | B11 | Gap | 55.31 |
| T12 | B6 | Control | 64.84 |
| T12 | B6 | O6-Me | 65.09 |
| T12 | B6 | Ribo | 64.72 |
| T12 | B6 | 8-oxoG | 65.62 |
| T12 | B6 | abasic | 62.88 |
| T12 | B6 | Nick | 59.46 |
| T12 | B6 | Gap | 58.36 |
| T12 | B9 | Control | 69.97 |
| T12 | B9 | O6-Me | 70.68 |
| T12 | B9 | Ribo | 70.41 |
| T12 | B9 | 8-oxoG | 70.93 |
| T12 | B9 | abasic | 68.64 |
| T12 | B9 | Nick | 64.60 |
| T12 | B9 | Gap | 62.62 |
| T12 | B11 | Control | 70.87 |
| T12 | B11 | O6-Me | 73.42 |
| T12 | B11 | Ribo | 70.25 |
| T12 | B11 | 8-oxoG | 70.72 |
| T12 | B11 | abasic | 69.72 |
| T12 | B11 | Nick | 64.67 |
| T12 | B11 | Gap | 63.15 |
| T18 | B6 | Control | 73.48 |
| T18 | B6 | O6-Me | 72.60 |
| T18 | B6 | Ribo | 73.59 |
| T18 | B6 | 8-oxoG | 73.20 |
| T18 | B6 | abasic | 69.08 |
| T18 | B6 | Nick | 69.04 |
| T18 | B6 | Gap | 68.33 |
| T18 | B9 | Control | 77.55 |

| | | | |
|---|---|---|---|
| T18 | B9 | O6-Me | 76.19 |
| T18 | B9 | Ribo | 76.22 |
| T18 | B9 | 8-oxoG | 77.54 |
| T18 | B9 | abasic | 75.95 |
| T18 | B9 | Nick | 71.15 |
| T18 | B9 | Gap | 71.00 |
| T18 | B11 | Control | 79.47 |
| T18 | B11 | O6-Me | 81.58 |
| T18 | B11 | Ribo | 79.10 |
| T18 | B11 | 8-oxoG | 80.33 |
| T18 | B11 | abasic | 78.12 |
| T18 | B11 | Nick | 73.11 |
| T18 | B11 | Gap | 71.47 |

| Construct | T7 (Å) | B6 (Å) |
|---|---|---|
| Ribo | 4.264 | 1.517 |
| 8-oxoG | 3.350 | 0.666 |
| Abasic | 2.707 | 2.477 |
| Nick | 5.208 | 2.611 |
| Gap | 3.598 | 2.306 |

| Construct | T12 (Å) | B9 (Å) |
|---|---|---|
| Ribo | 2.044 | 1.814 |
| 8-oxoG | 2.346 | 1.238 |
| Abasic | 1.284 | 0.682 |
| Nick | 2.965 | 2.902 |
| Gap | 3.483 | 3.397 |

| Construct | T18 (Å) | B11 (Å) |
|---|---|---|
| Ribo | 3.566 | 1.233 |
| 8-oxoG | 3.401 | 1.031 |
| Abasic | 2.958 | 1.073 |
| Nick | 5.587 | 3.356 |
| Gap | 4.305 | 5.124 |

| **Construct** | **Mean Difference:** | **Mean Relative Difference:** | **Standard Deviation of Differences:** |
|---|---|---|---|
| | **(Å)** | | |
| **Ribo** | 1.44E-05 | 2.52E-09 | 5.46E-06 |
| **8-oxoG** | 2.44E-06 | 4.66E-08 | 1.52E-06 |
| **Abasic** | 1.56E-05 | 2.66E-07 | 9.40E-06 |
| **Nick** | 6.84E-06 | 1.08E-07 | 5.53E-06 |
| **Gap** | 5.67E-06 | 1.21E-07 | 3.61E-06 |

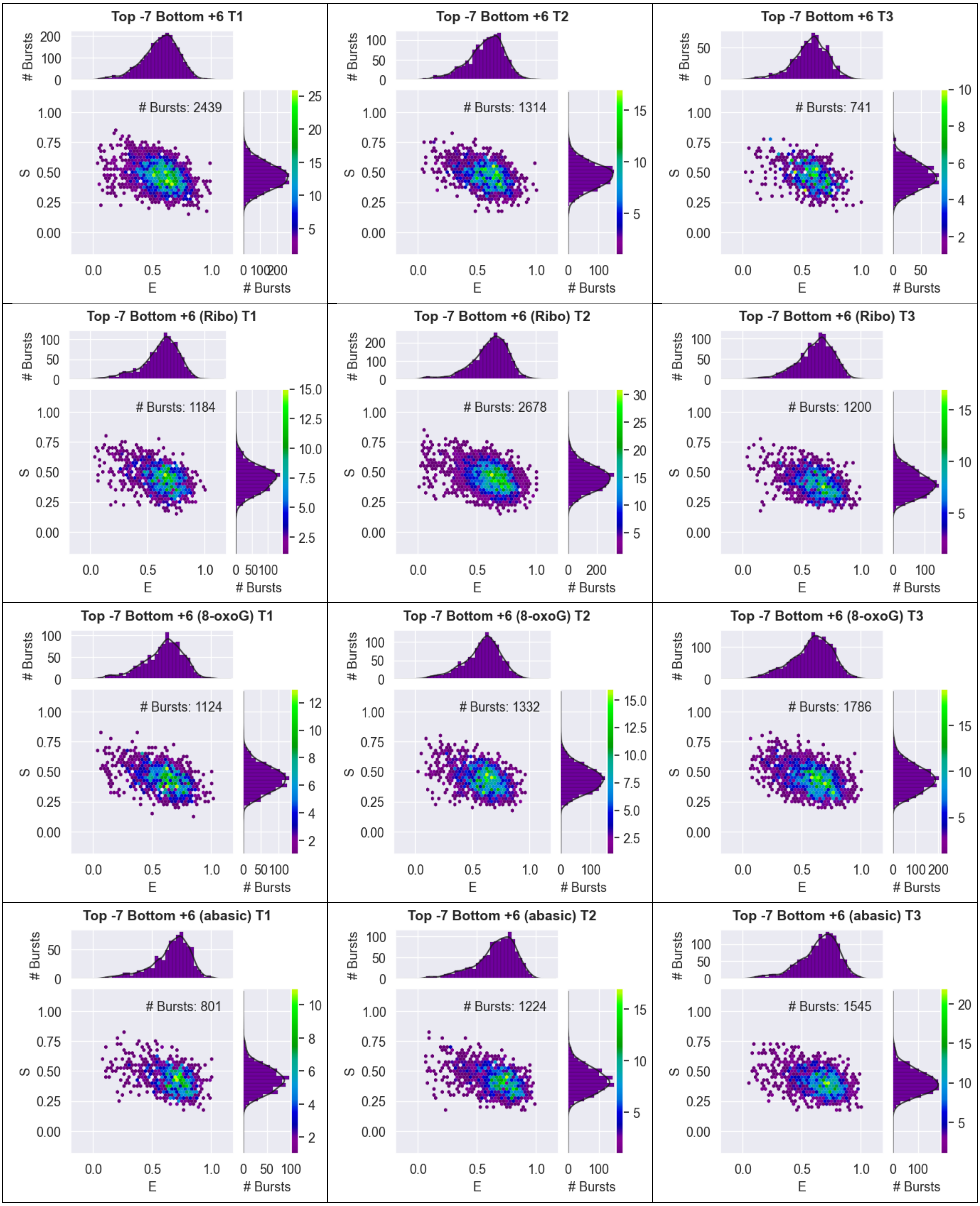
Top -7 Bottom +6 T1
### Bursts: 2439
Top -7 Bottom +6 T2
### Bursts: 1314
Top -7 Bottom +6 T3
### Bursts: 741
Top -7 Bottom +6 (Ribo) T1
### Bursts: 1184
Top -7 Bottom +6 (Ribo) T2
### Bursts: 2678
Top -7 Bottom +6 (Ribo) T3
### Bursts: 1200
Top -7 Bottom +6 (8-oxoG) T1
### Bursts: 1124
Top -7 Bottom +6 (8-oxoG) T2
### Bursts: 1332
Top -7 Bottom +6 (8-oxoG) T3
### Bursts: 1786
Top -7 Bottom +6 (abasic) T1
### Bursts: 801
Top -7 Bottom +6 (abasic) T2
### Bursts: 1224
Top -7 Bottom +6 (abasic) T3
### Bursts: 1545
E
S
### Bursts

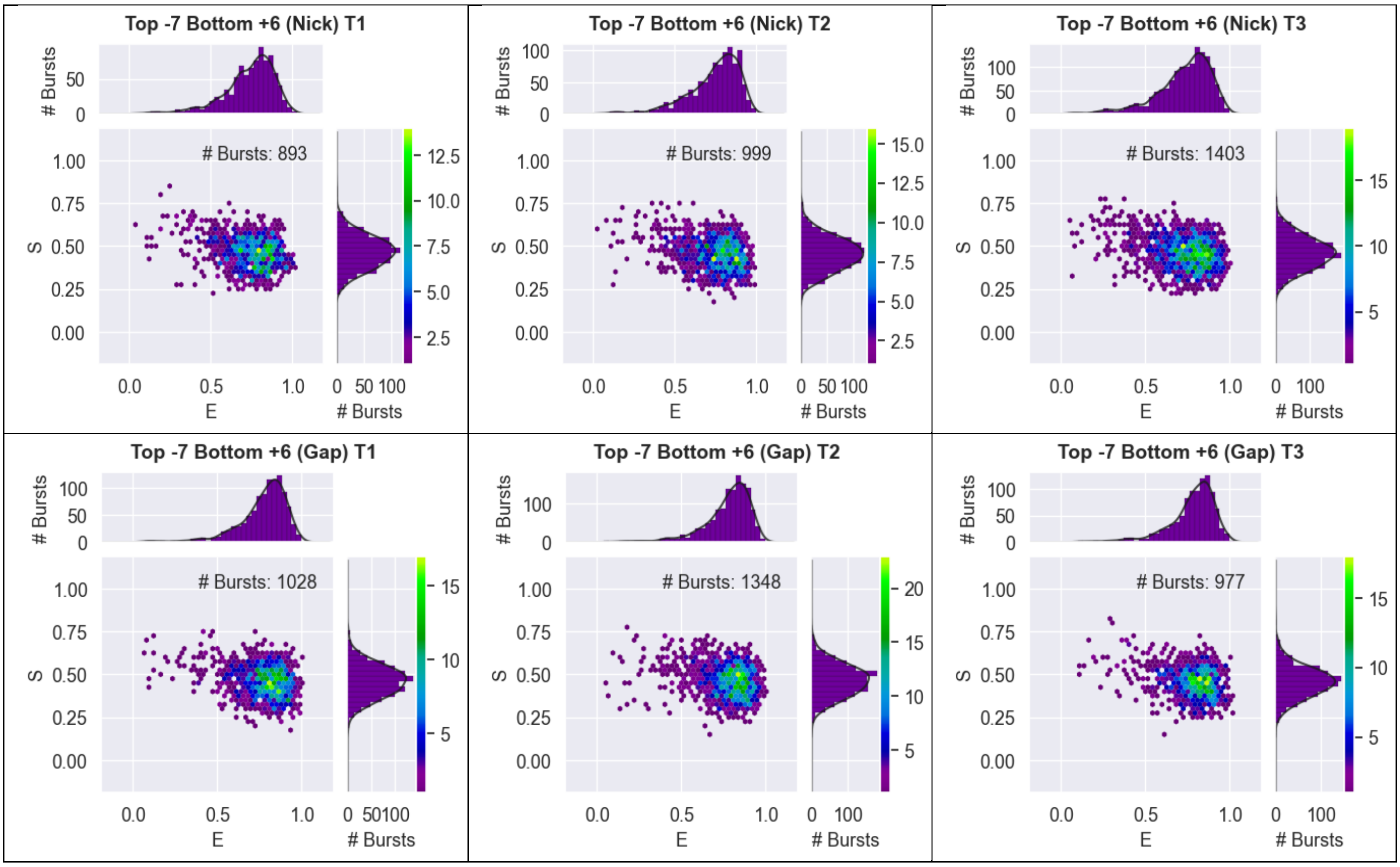

Supplementary Figure 1: 2D ES histograms from DCBS analysis showing FRET efficiency (E) and stoichiometry (S) for each independent repeat of the seven duplex constructs for the dye-pair Top -7 Bottom +6.

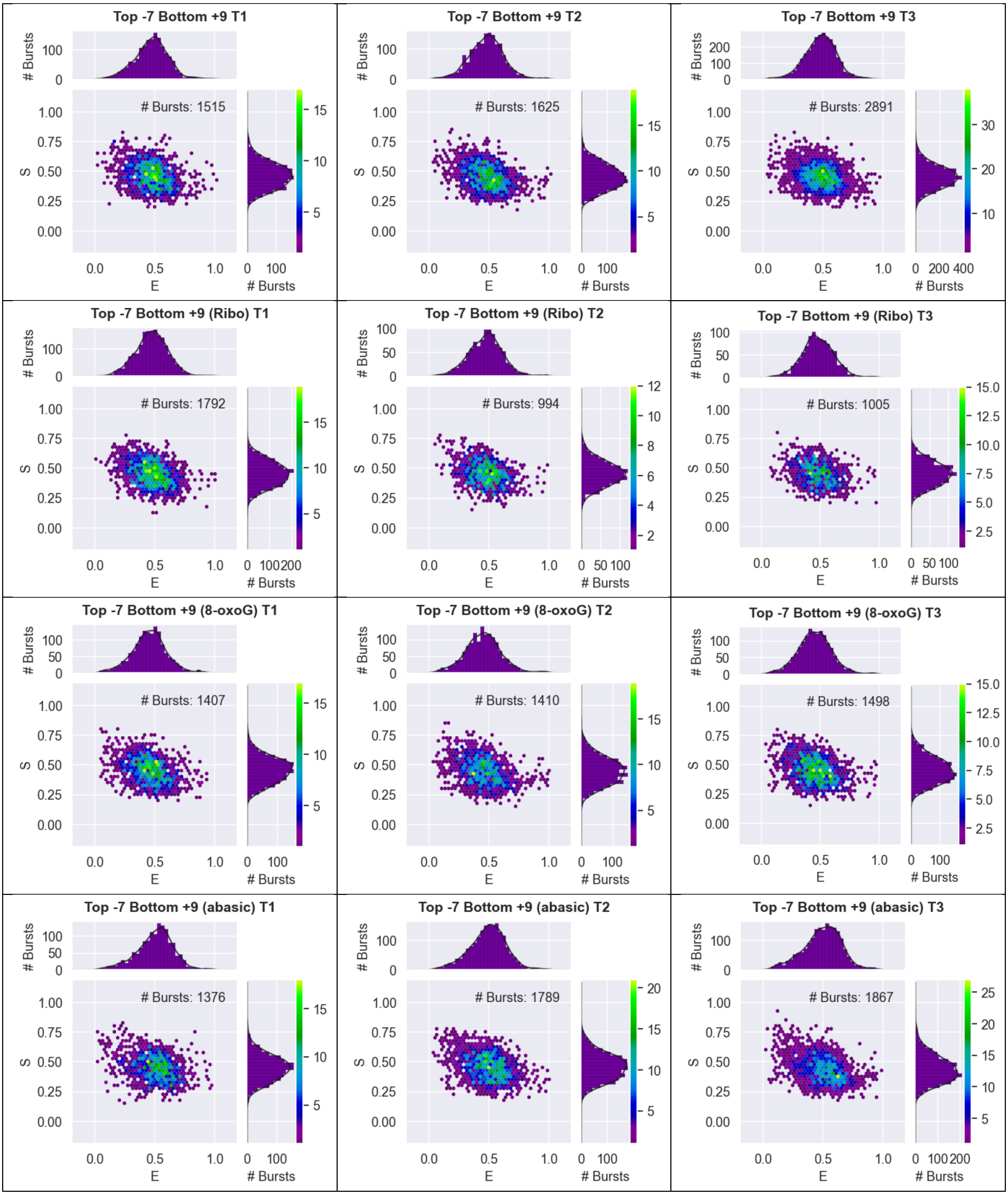

Top -7 Bottom +9 T1
### Bursts: 1515
Top -7 Bottom +9 T2
### Bursts: 1625
Top -7 Bottom +9 T3
### Bursts: 2891
Top -7 Bottom +9 (Ribo) T1
### Bursts: 1792
Top -7 Bottom +9 (Ribo) T2
### Bursts: 994
Top -7 Bottom +9 (Ribo) T3
### Bursts: 1005
Top -7 Bottom +9 (8-oxoG) T1
### Bursts: 1407
Top -7 Bottom +9 (8-oxoG) T2
### Bursts: 1410
Top -7 Bottom +9 (8-oxoG) T3
### Bursts: 1498
Top -7 Bottom +9 (abasic) T1
### Bursts: 1376
Top -7 Bottom +9 (abasic) T2
### Bursts: 1789
Top -7 Bottom +9 (abasic) T3
### Bursts: 1867
E
S
### Bursts

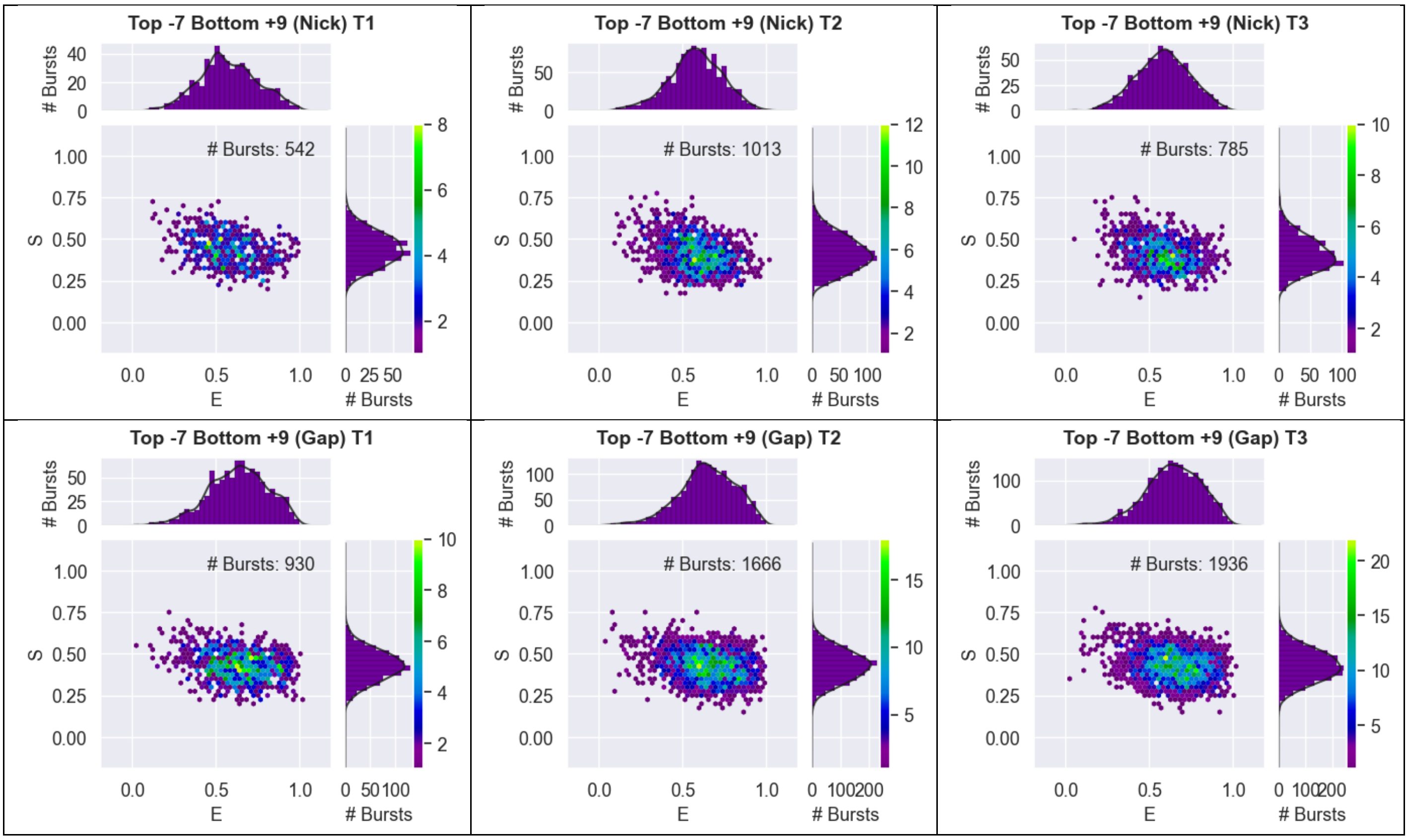

Supplementary Figure 2: 2D ES histograms from DCBS analysis showing FRET efficiency (E) and stoichiometry (S) for each independent repeat of the seven duplex constructs for the dye-pair Top -7 Bottom +9.

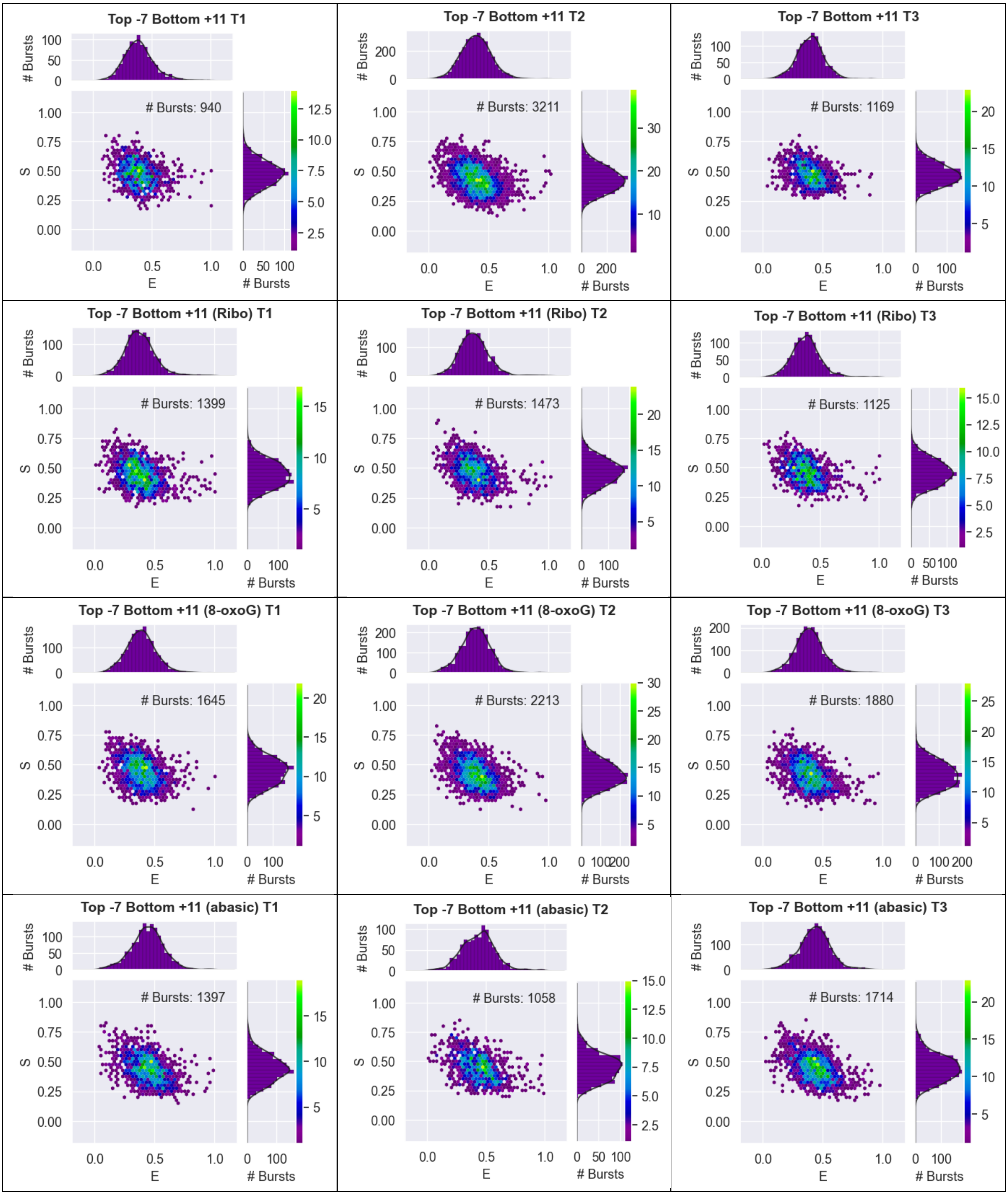
Top -7 Bottom +11 T1
### Bursts: 940
Top -7 Bottom +11 T2
### Bursts: 3211
Top -7 Bottom +11 T3
### Bursts: 1169
Top -7 Bottom +11 (Ribo) T1
### Bursts: 1399
Top -7 Bottom +11 (Ribo) T2
### Bursts: 1473
Top -7 Bottom +11 (Ribo) T3
### Bursts: 1125
Top -7 Bottom +11 (8-oxoG) T1
### Bursts: 1645
Top -7 Bottom +11 (8-oxoG) T2
### Bursts: 2213
Top -7 Bottom +11 (8-oxoG) T3
### Bursts: 1880
Top -7 Bottom +11 (abasic) T1
### Bursts: 1397
Top -7 Bottom +11 (abasic) T2
### Bursts: 1058
Top -7 Bottom +11 (abasic) T3
### Bursts: 1714
S
E
### Bursts

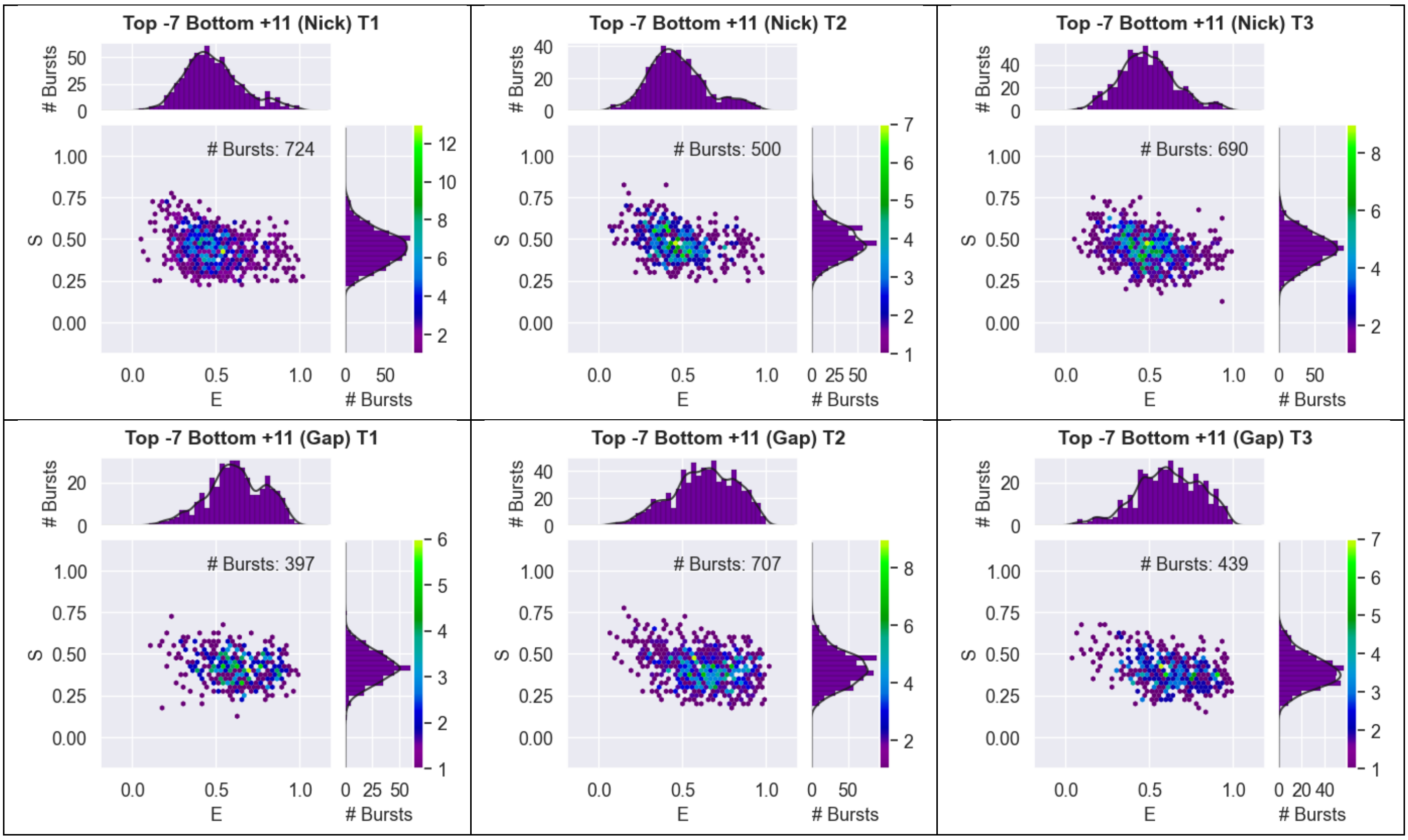

Supplementary Figure 3: 2D ES histograms from DCBS analysis showing FRET efficiency (E) and stoichiometry (S) for each independent repeat of the seven duplex constructs for the dye-pair Top -7 Bottom +11.

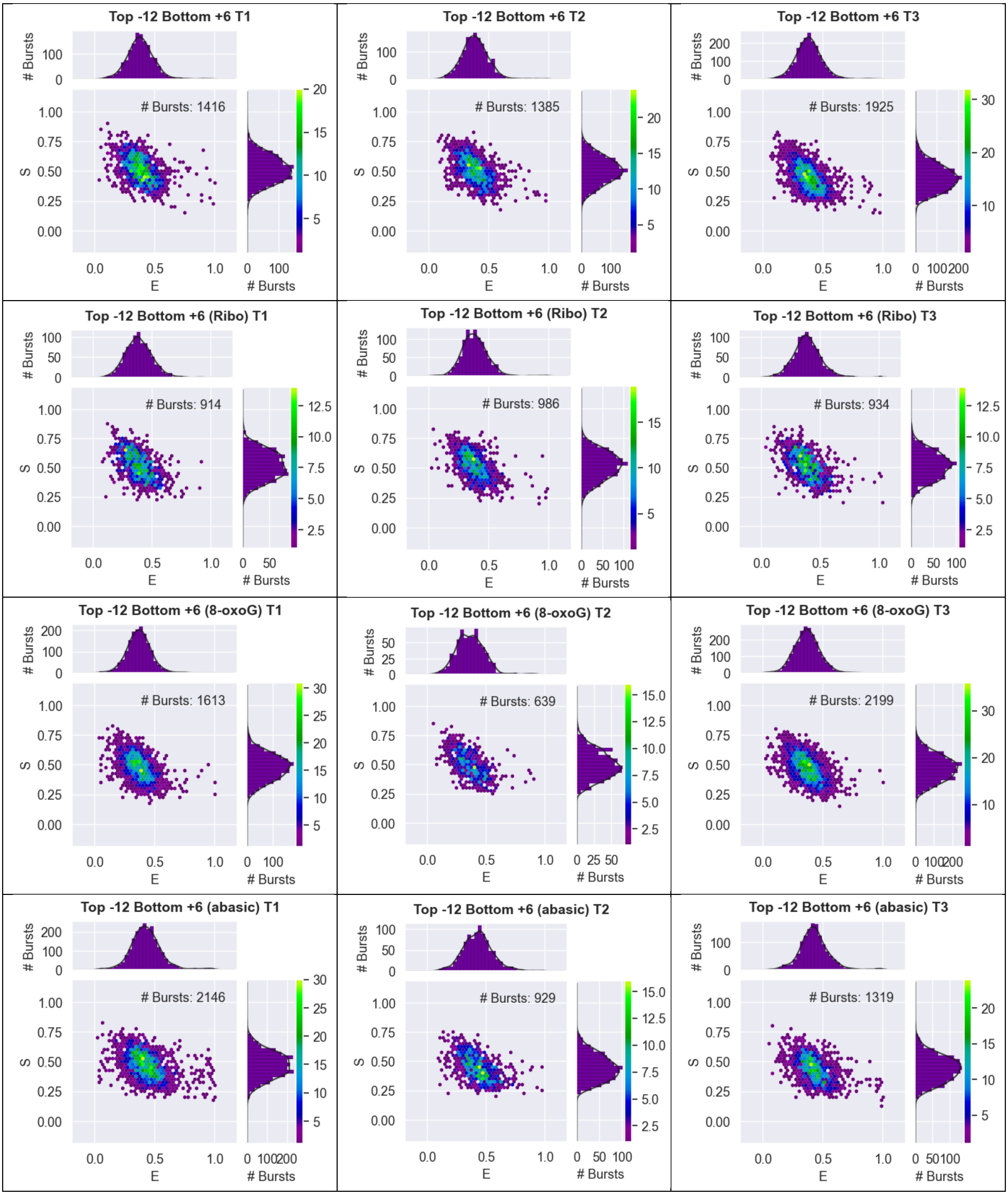
Top -12 Bottom +6 T1
### Bursts: 1416
Top -12 Bottom +6 T2
### Bursts: 1385
Top -12 Bottom +6 T3
### Bursts: 1925
Top -12 Bottom +6 (Ribo) T1
### Bursts: 914
Top -12 Bottom +6 (Ribo) T2
### Bursts: 986
Top -12 Bottom +6 (Ribo) T3
### Bursts: 934
Top -12 Bottom +6 (8-oxoG) T1
### Bursts: 1613
Top -12 Bottom +6 (8-oxoG) T2
### Bursts: 639
Top -12 Bottom +6 (8-oxoG) T3
### Bursts: 2199
Top -12 Bottom +6 (abasic) T1
### Bursts: 2146
Top -12 Bottom +6 (abasic) T2
### Bursts: 929
Top -12 Bottom +6 (abasic) T3
### Bursts: 1319
S
E
### Bursts

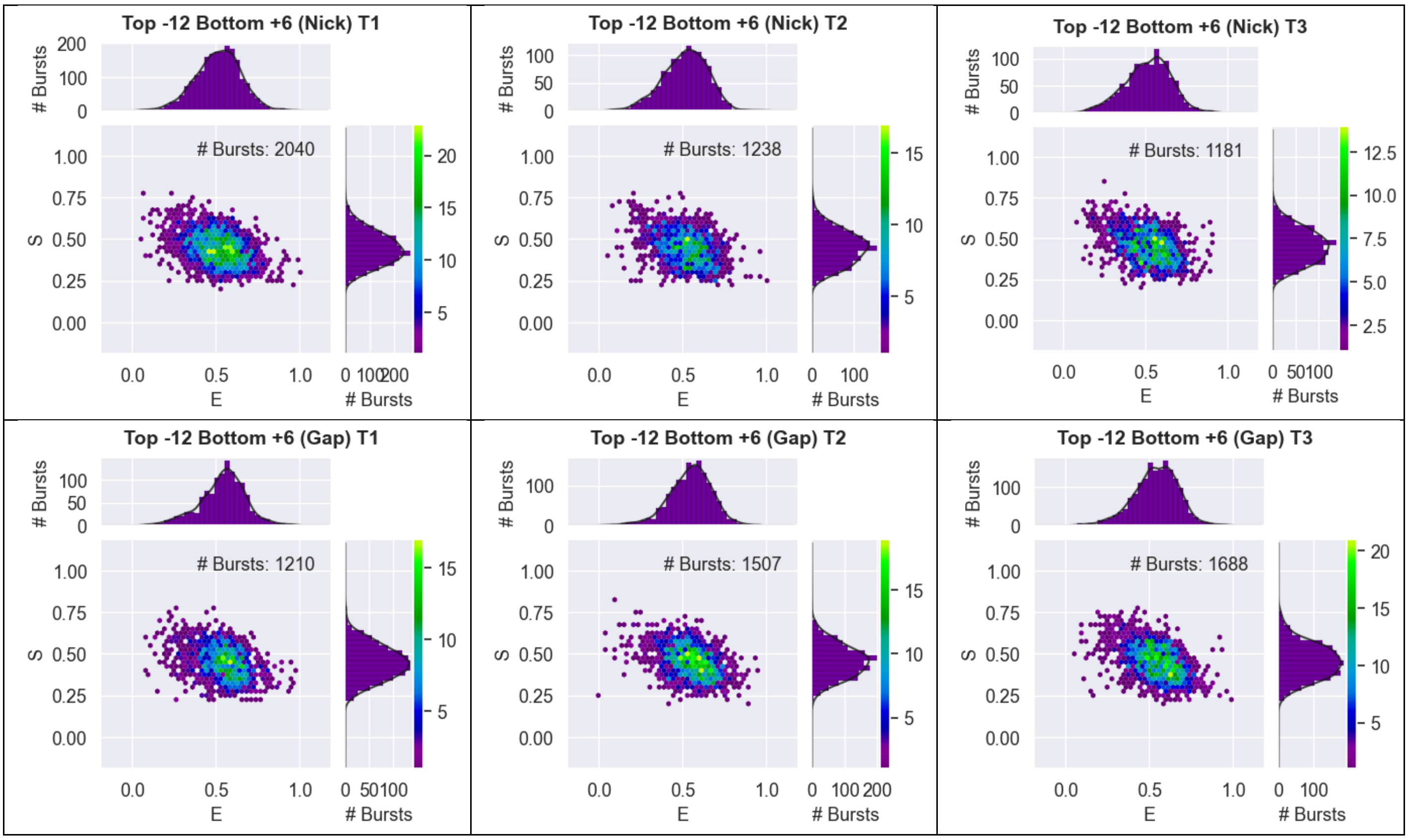

Supplementary Figure 4: 2D ES histograms from DCBS analysis showing FRET efficiency (E) and stoichiometry (S) for each independent repeat of the seven duplex constructs for the dye-pair Top -12 Bottom +6.

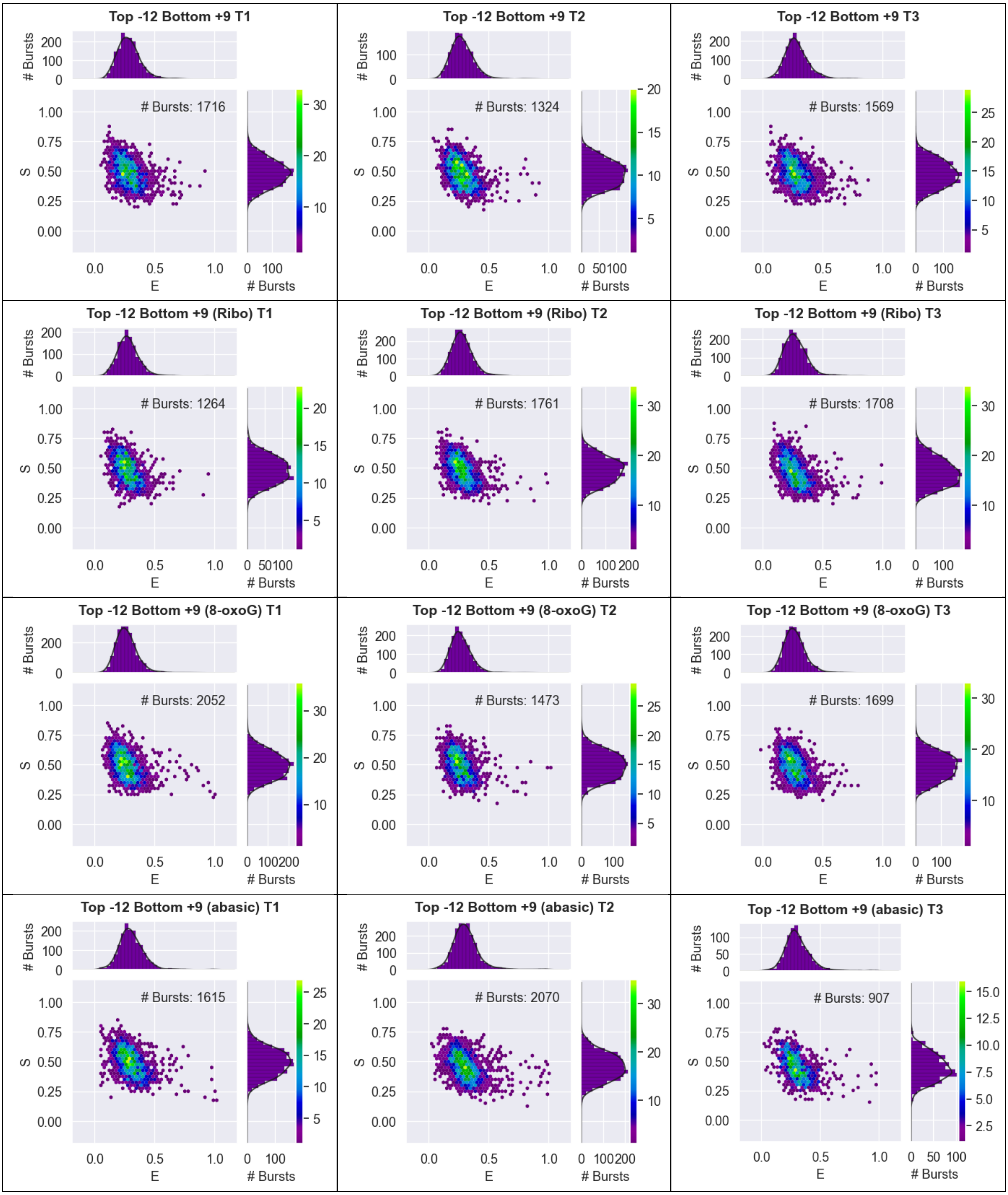
Top -12 Bottom +9 T1
### Bursts: 1716
Top -12 Bottom +9 T2
### Bursts: 1324
Top -12 Bottom +9 T3
### Bursts: 1569
Top -12 Bottom +9 (Ribo) T1
### Bursts: 1264
Top -12 Bottom +9 (Ribo) T2
### Bursts: 1761
Top -12 Bottom +9 (Ribo) T3
### Bursts: 1708
Top -12 Bottom +9 (8-oxoG) T1
### Bursts: 2052
Top -12 Bottom +9 (8-oxoG) T2
### Bursts: 1473
Top -12 Bottom +9 (8-oxoG) T3
### Bursts: 1699
Top -12 Bottom +9 (abasic) T1
### Bursts: 1615
Top -12 Bottom +9 (abasic) T2
### Bursts: 2070
Top -12 Bottom +9 (abasic) T3
### Bursts: 907
S
E
### Bursts

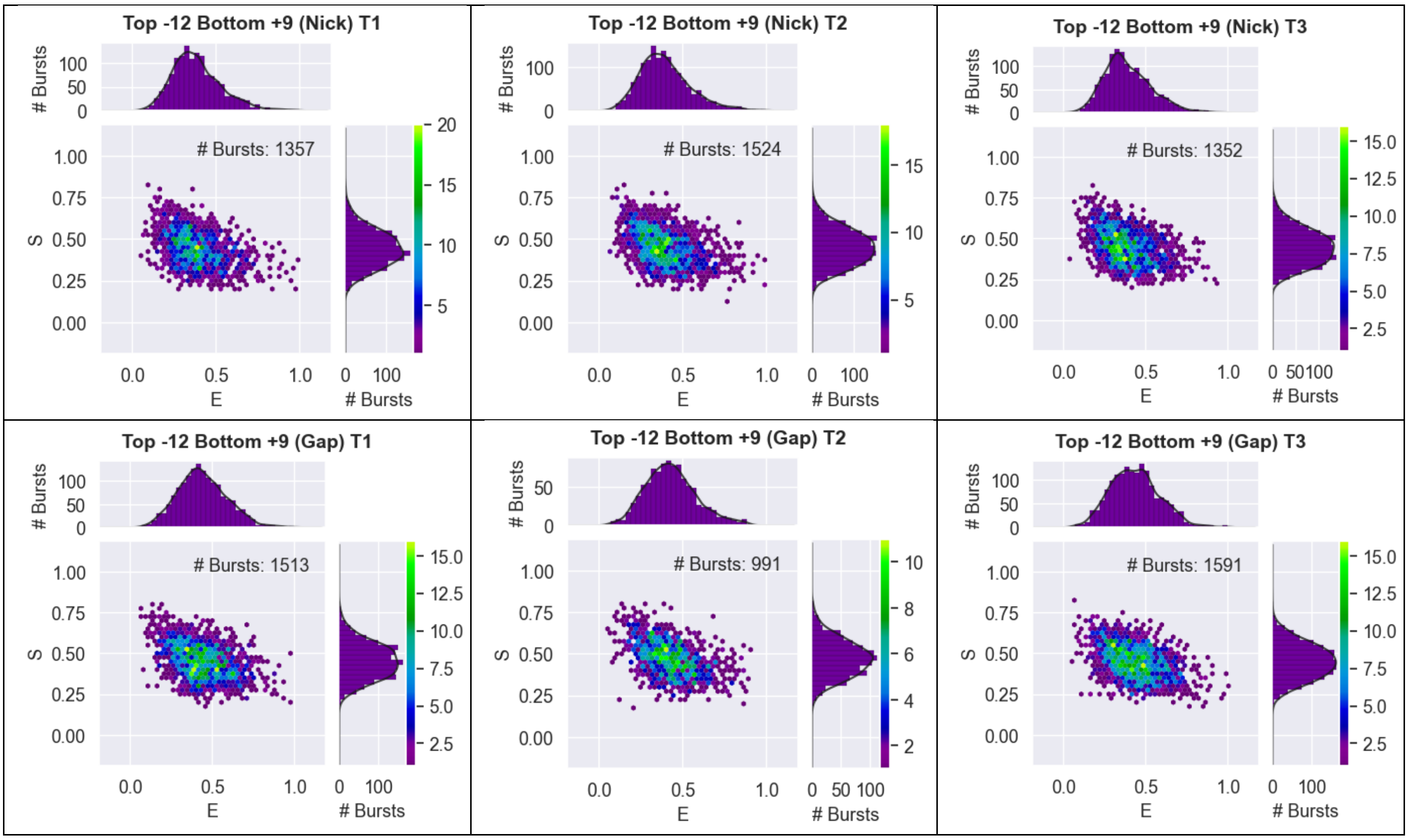

Supplementary Figure 5: 2D ES histograms from DCBS analysis showing FRET efficiency (E) and stoichiometry (S) for each independent repeat of the seven duplex constructs for the dye-pair Top -12 Bottom +9.

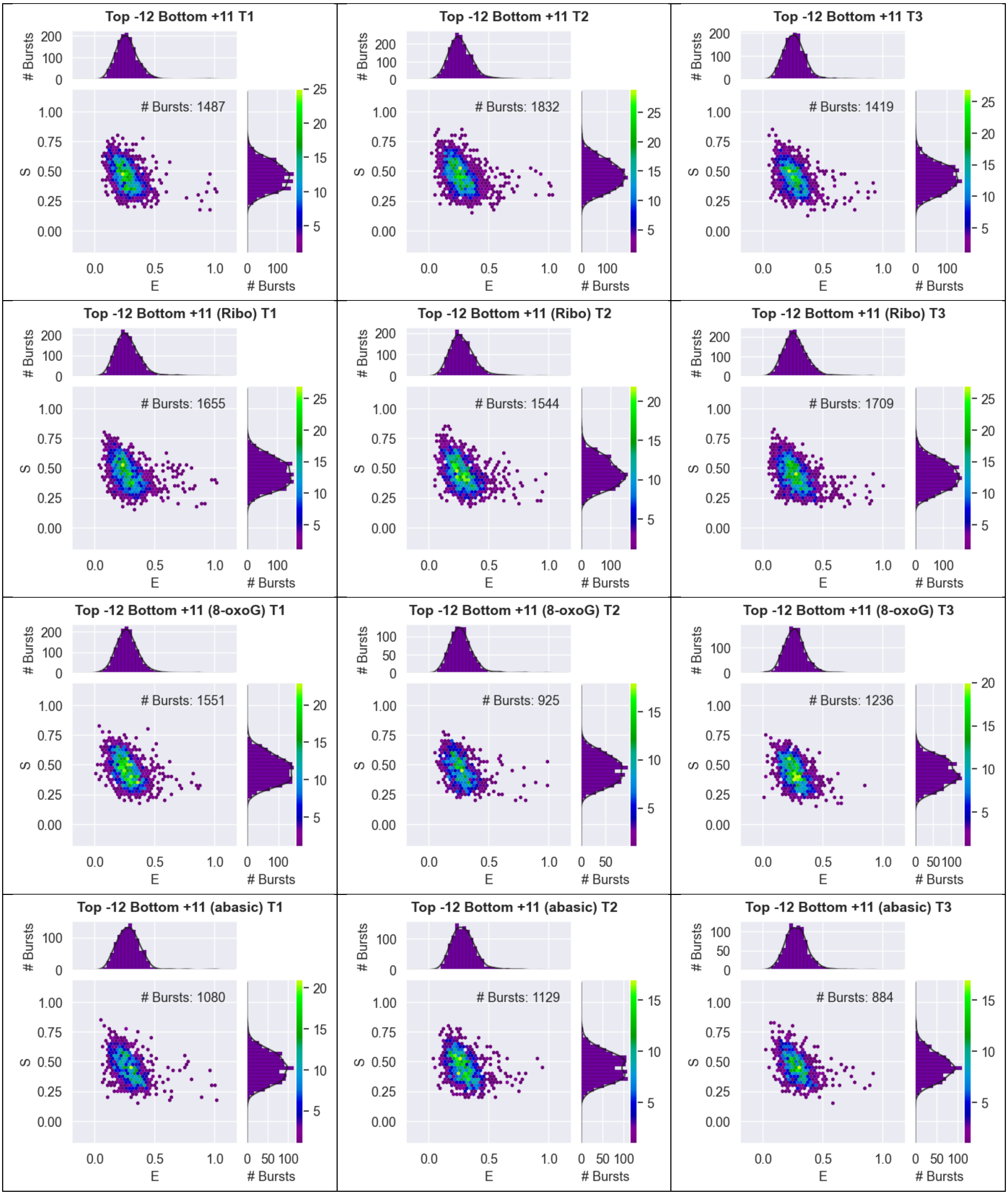
Top -12 Bottom +11 T1
### Bursts: 1487
Top -12 Bottom +11 T2
### Bursts: 1832
Top -12 Bottom +11 T3
### Bursts: 1419
Top -12 Bottom +11 (Ribo) T1
### Bursts: 1655
Top -12 Bottom +11 (Ribo) T2
### Bursts: 1544
Top -12 Bottom +11 (Ribo) T3
### Bursts: 1709
Top -12 Bottom +11 (8-oxoG) T1
### Bursts: 1551
Top -12 Bottom +11 (8-oxoG) T2
### Bursts: 925
Top -12 Bottom +11 (8-oxoG) T3
### Bursts: 1236
Top -12 Bottom +11 (abasic) T1
### Bursts: 1080
Top -12 Bottom +11 (abasic) T2
### Bursts: 1129
Top -12 Bottom +11 (abasic) T3
### Bursts: 884
### Bursts
S
E

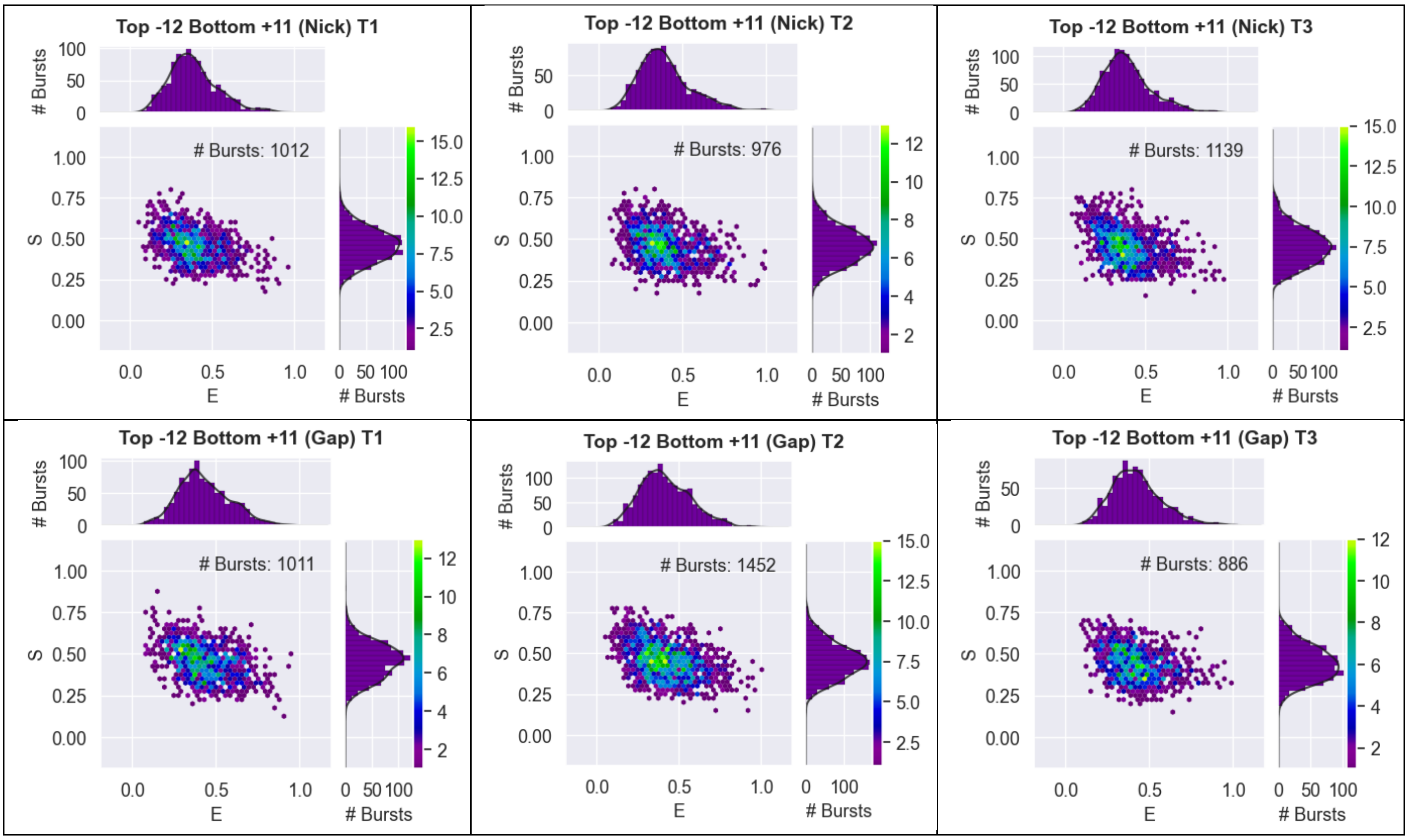

Supplementary Figure 6: 2D ES histograms from DCBS analysis showing FRET efficiency (E) and stoichiometry (S) for each independent repeat of the seven duplex constructs for the dye-pair Top -12 Bottom +11.

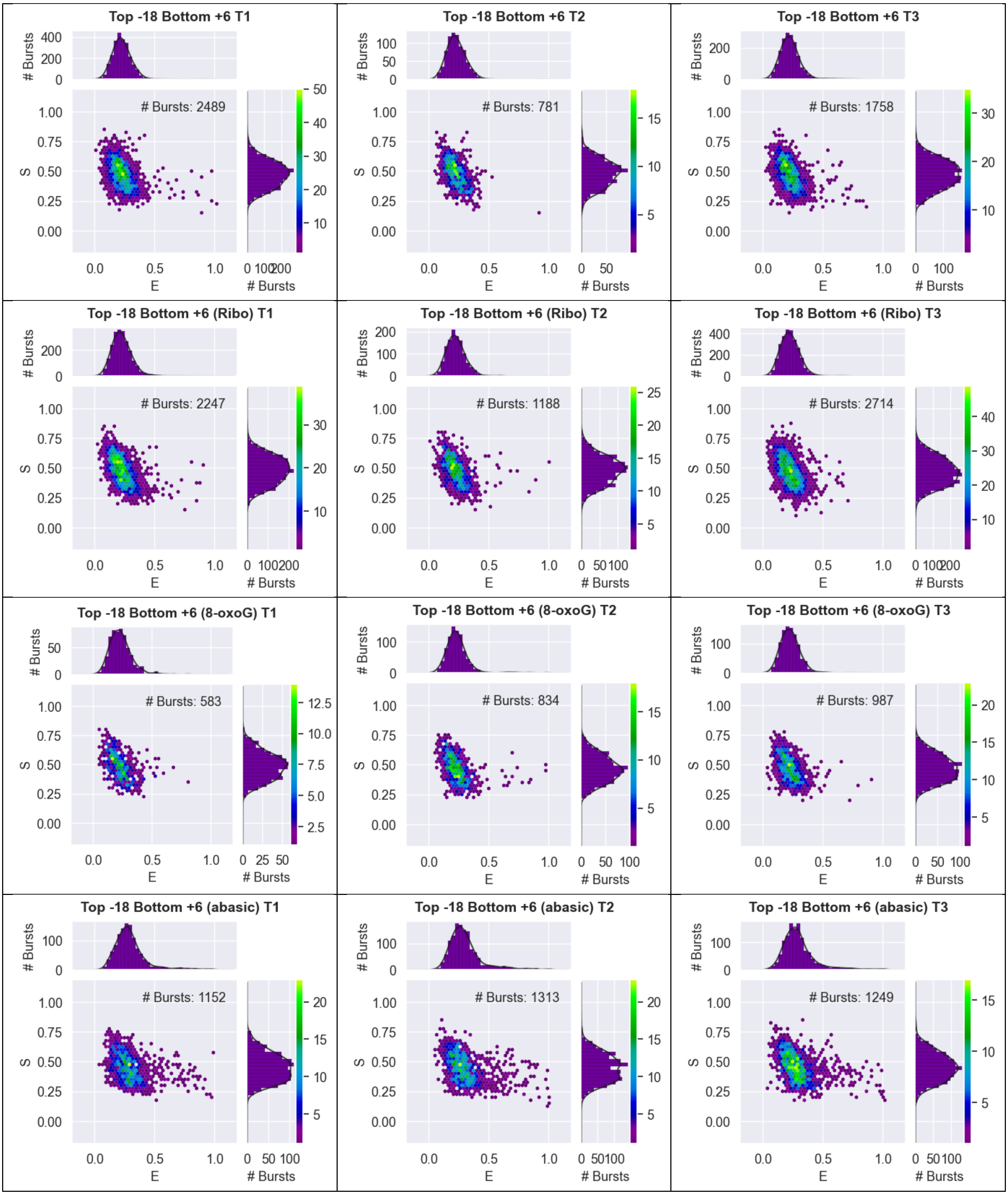

Top -18 Bottom +6 T1
### Bursts: 2489
Top -18 Bottom +6 T2
### Bursts: 781
Top -18 Bottom +6 T3
### Bursts: 1758
Top -18 Bottom +6 (Ribo) T1
### Bursts: 2247
Top -18 Bottom +6 (Ribo) T2
### Bursts: 1188
Top -18 Bottom +6 (Ribo) T3
### Bursts: 2714
Top -18 Bottom +6 (8-oxoG) T1
### Bursts: 583
Top -18 Bottom +6 (8-oxoG) T2
### Bursts: 834
Top -18 Bottom +6 (8-oxoG) T3
### Bursts: 987
Top -18 Bottom +6 (abasic) T1
### Bursts: 1152
Top -18 Bottom +6 (abasic) T2
### Bursts: 1313
Top -18 Bottom +6 (abasic) T3
### Bursts: 1249
E
S
### Bursts

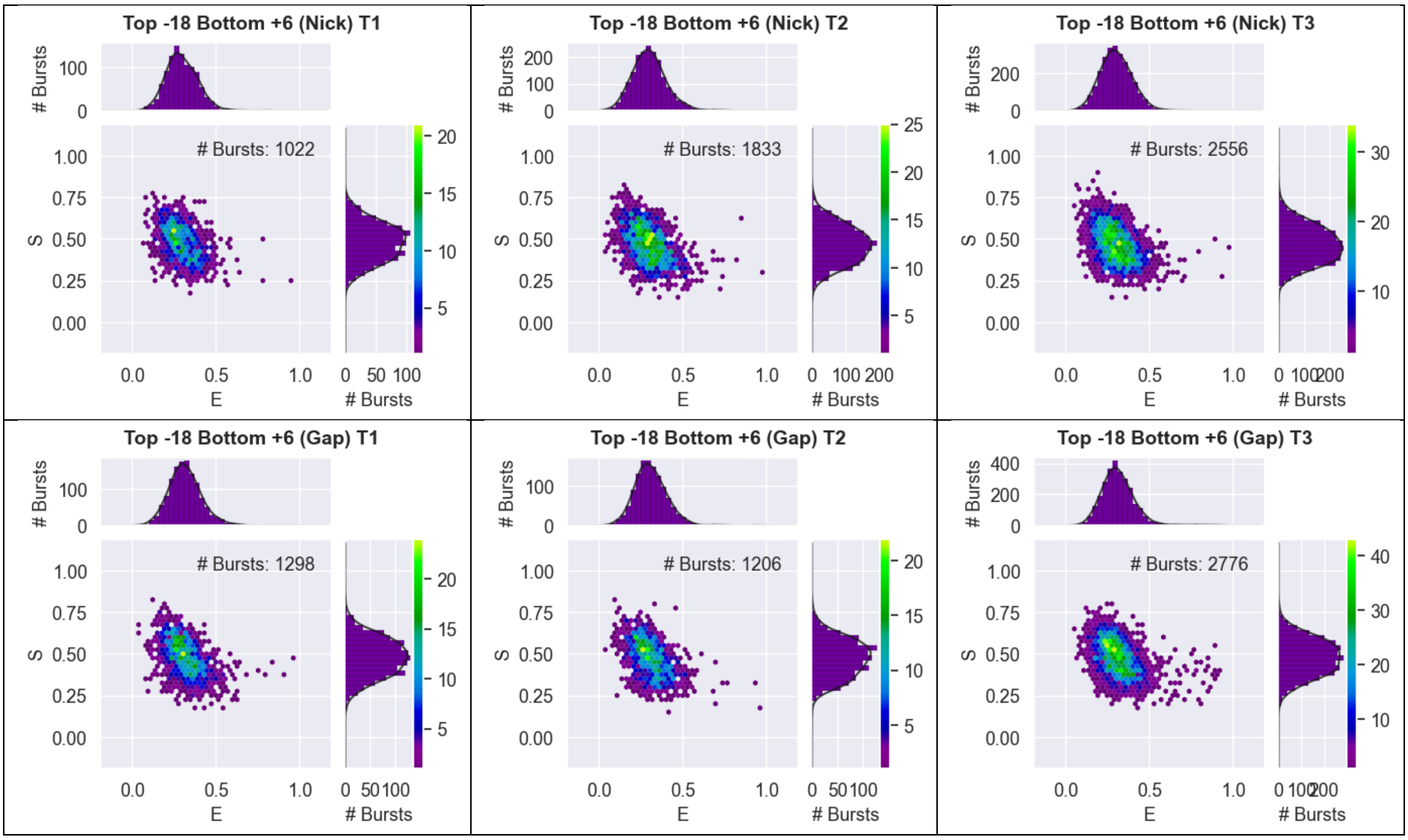

Supplementary Figure 7: 2D ES histograms from DCBS analysis showing FRET efficiency (E) and stoichiometry (S) for each independent repeat of the seven duplex constructs for the dye-pair Top -18 Bottom +6.

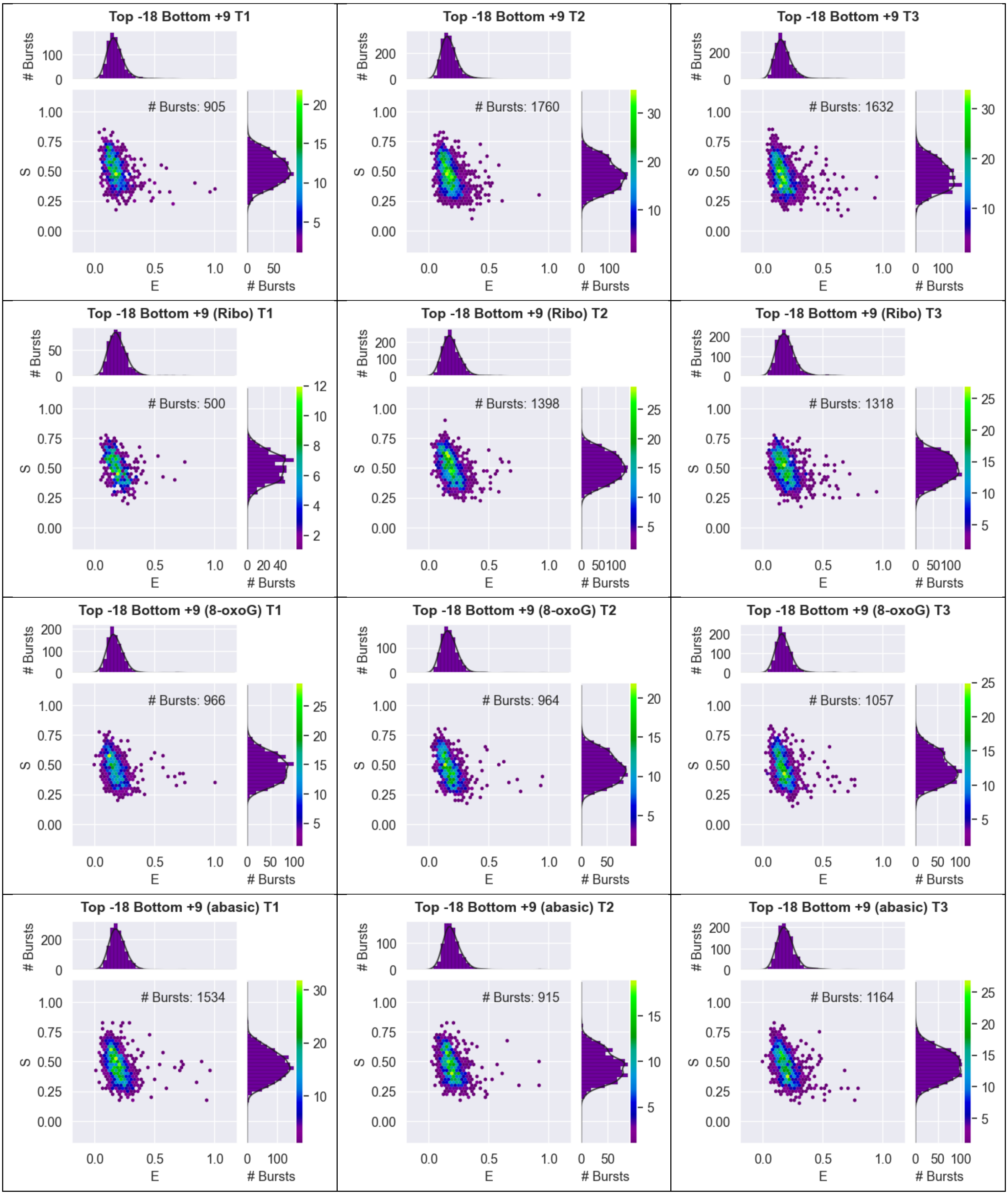
Top -18 Bottom +9 T1
### Bursts: 905
Top -18 Bottom +9 T2
### Bursts: 1760
Top -18 Bottom +9 T3
### Bursts: 1632
Top -18 Bottom +9 (Ribo) T1
### Bursts: 500
Top -18 Bottom +9 (Ribo) T2
### Bursts: 1398
Top -18 Bottom +9 (Ribo) T3
### Bursts: 1318
Top -18 Bottom +9 (8-oxoG) T1
### Bursts: 966
Top -18 Bottom +9 (8-oxoG) T2
### Bursts: 964
Top -18 Bottom +9 (8-oxoG) T3
### Bursts: 1057
Top -18 Bottom +9 (abasic) T1
### Bursts: 1534
Top -18 Bottom +9 (abasic) T2
### Bursts: 915
Top -18 Bottom +9 (abasic) T3
### Bursts: 1164
S
E
### Bursts

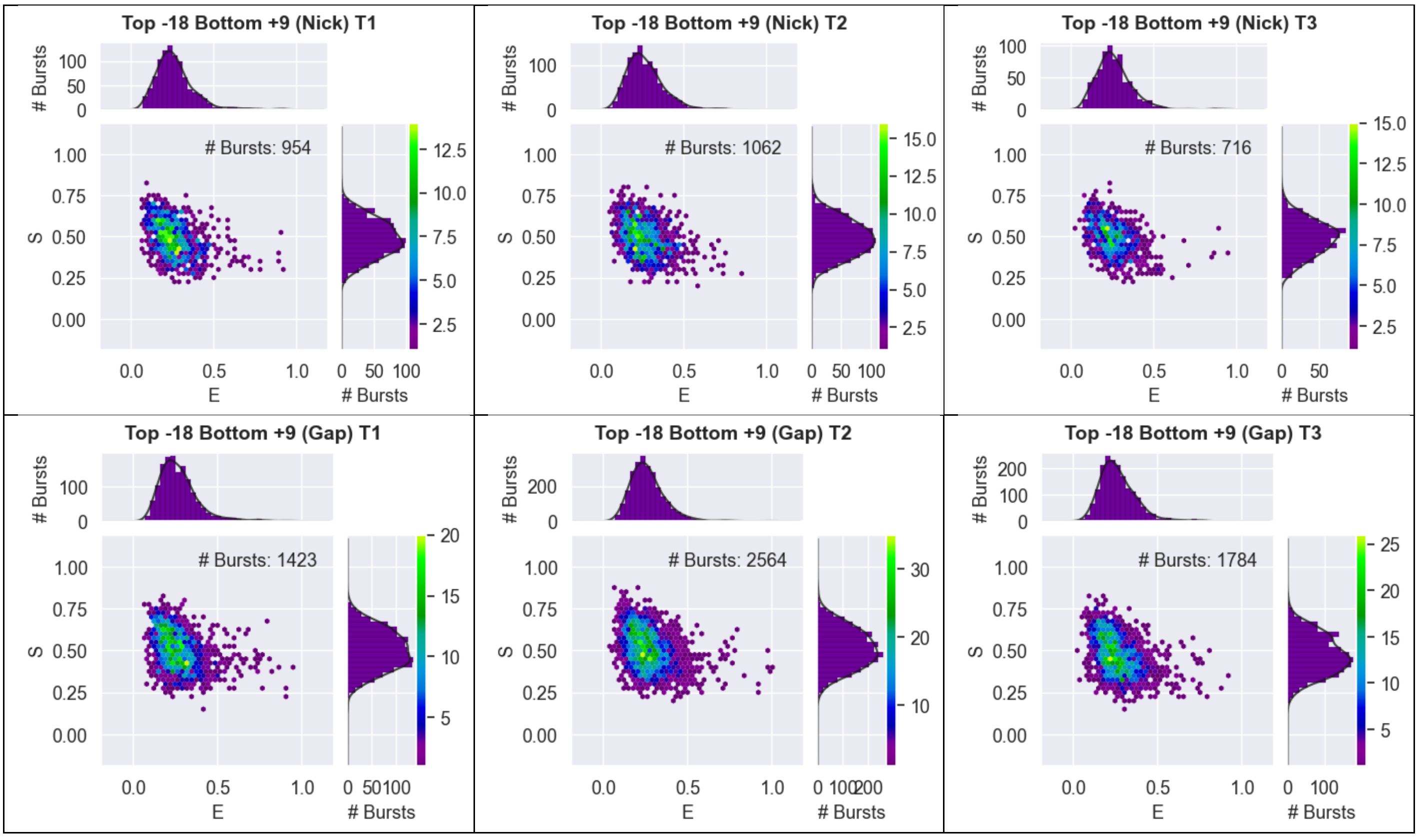

Supplementary Figure 8: 2D ES histograms from DCBS analysis showing FRET efficiency (E) and stoichiometry (S) for each independent repeat of the seven duplex constructs for the dye-pair Top -18 Bottom +9.

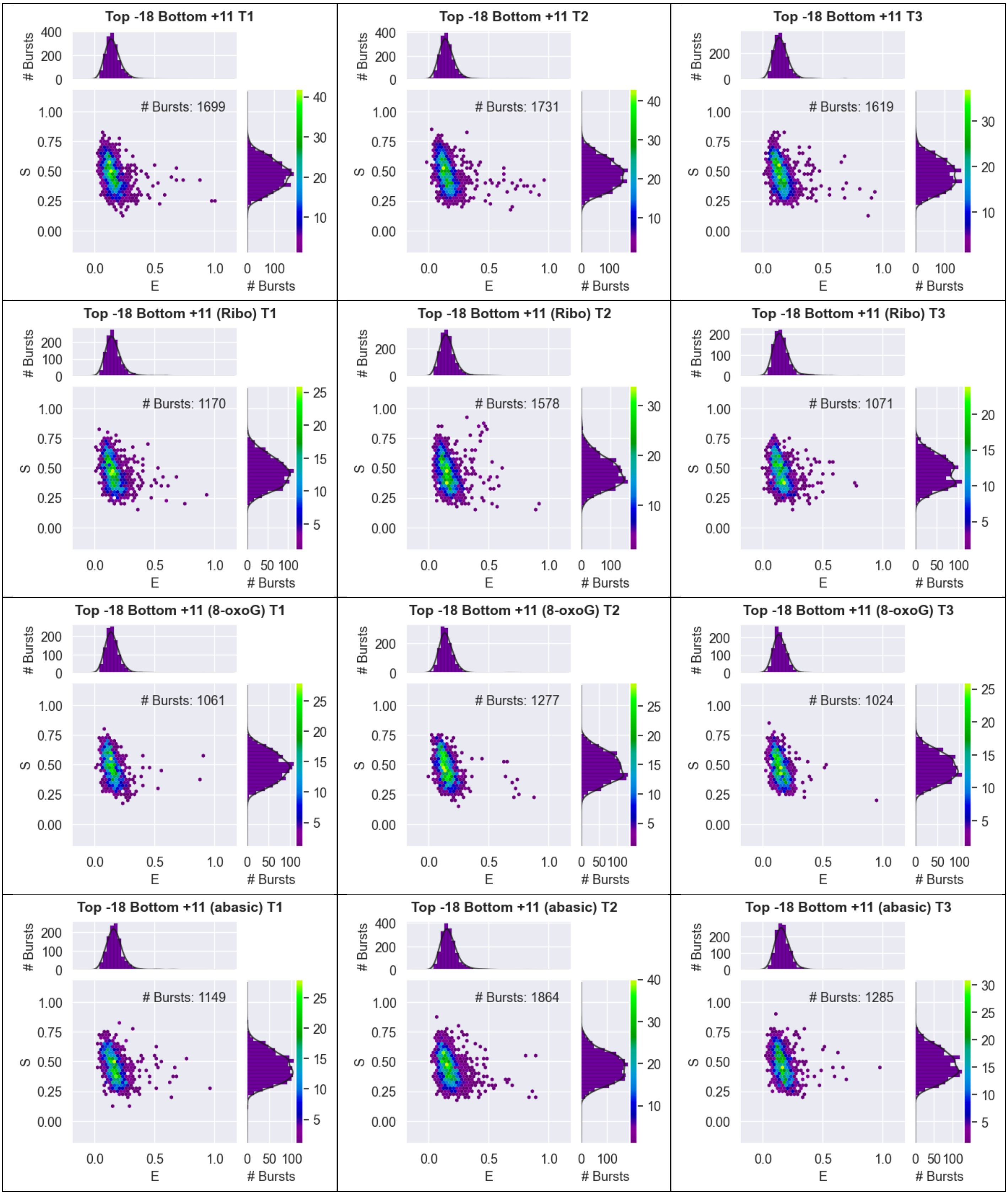
Top -18 Bottom +11 T1
### Bursts: 1699
Top -18 Bottom +11 T2
### Bursts: 1731
Top -18 Bottom +11 T3
### Bursts: 1619
Top -18 Bottom +11 (Ribo) T1
### Bursts: 1170
Top -18 Bottom +11 (Ribo) T2
### Bursts: 1578
Top -18 Bottom +11 (Ribo) T3
### Bursts: 1071
Top -18 Bottom +11 (8-oxoG) T1
### Bursts: 1061
Top -18 Bottom +11 (8-oxoG) T2
### Bursts: 1277
Top -18 Bottom +11 (8-oxoG) T3
### Bursts: 1024
Top -18 Bottom +11 (abasic) T1
### Bursts: 1149
Top -18 Bottom +11 (abasic) T2
### Bursts: 1864
Top -18 Bottom +11 (abasic) T3
### Bursts: 1285
E
S
### Bursts

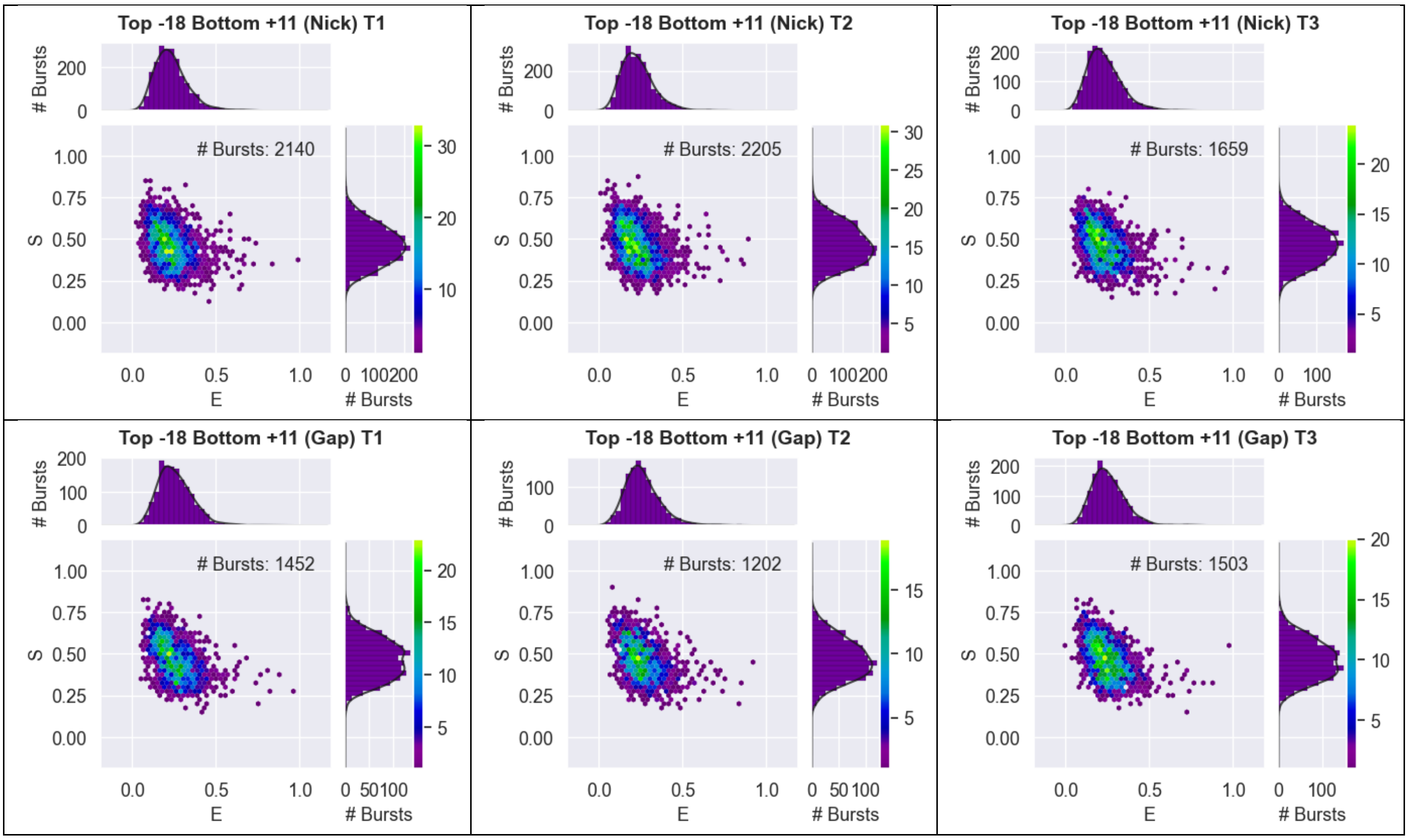

Supplementary Figure 9: 2D ES histograms from DCBS analysis showing FRET efficiency (E) and stoichiometry (S) for each independent repeat of the seven duplex constructs for the dye-pair Top -18 Bottom +11.

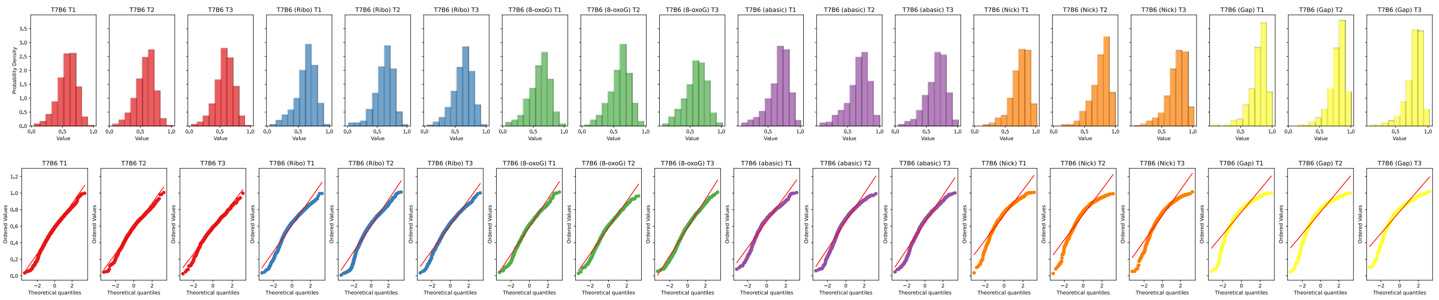

| Construct | Kolmogorov–Smirnov test statistic | p-value* |
|---|---|---|
| T7B6 T1 | 0.5678 | 0 |
| T7B6 T2 | 0.5681 | 0 |
| T7B6 T3 | 0.565 | 8.634e-224 |
| T7B6 (Ribo) T1 | 0.5684 | 0 |
| T7B6 (Ribo) T2 | 0.5765 | 0 |
| T7B6 (Ribo) T3 | 0.5808 | 0 |
| T7B6 (8-oxoG) T1 | 0.5585 | 0 |
| T7B6 (8-oxoG) T2 | 0.5635 | 0 |
| T7B6 (8-oxoG) T3 | 0.5559 | 0 |
| T7B6 (abasic) T1 | 0.5785 | 9.861e-255 |
| T7B6 (abasic) T2 | 0.5757 | 0 |
| T7B6 (abasic) T3 | 0.5818 | 0 |
| T7B6 (Nick) T1 | 0.6177 | 0 |
| T7B6 (Nick) T2 | 0.6207 | 0 |
| T7B6 (Nick) T3 | 0.612 | 0 |
| T7B6 (Gap) T1 | 0.6442 | 0 |
| T7B6 (Gap) T2 | 0.6417 | 0 |
| T7B6 (Gap) T3 | 0.646 | 0 |

*p-values given as 0 mean $< 10^{-307}$

Kruskal-Wallis test statistic: 5056
p-value: 0

Supplementary Figure 10: Quantile-Quantile (QQ) plots, and the Kolmogorov-Smirnov test for each independent repeat of the seven duplex constructs for the dye-pair Top -7 Bottom +6. Kruskal-Wallis test result to confirm that within all the groups there were samples originating from different distributions.

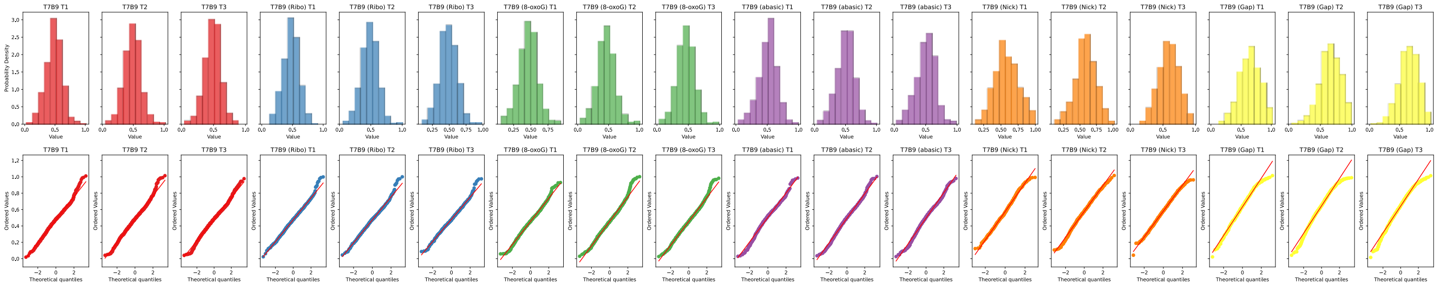

| Construct | Kolmogorov–Smirnov test statistic | p-value |
|---|---|---|
| T7B9 T1 | 0.5499 | 0 |
| T7B9 T2 | 0.5543 | 0 |
| T7B9 T3 | 0.5593 | 0 |
| T7B9 (Ribo) T1 | 0.559 | 0 |
| T7B9 (Ribo) T2 | 0.5568 | 1.485e-290 |
| T7B9 (Ribo) T3 | 0.562 | 8.072e-300 |
| T7B9 (8-oxoG) T1 | 0.5403 | 0 |
| T7B9 (8-oxoG) T2 | 0.5381 | 0 |
| T7B9 (8-oxoG) T3 | 0.5429 | 0 |
| T7B9 (abasic) T1 | 0.545 | 0 |
| T7B9 (abasic) T2 | 0.5445 | 0 |
| T7B9 (abasic) T3 | 0.5391 | 0 |
| T7B9 (Nick) T1 | 0.5753 | 1.679e-170 |
| T7B9 (Nick) T2 | 0.5719 | 5.625e-314 |
| T7B9 (Nick) T3 | 0.583 | 5.777e-254 |
| T7B9 (Gap) T1 | 0.5846 | 1.229e-302 |
| T7B9 (Gap) T2 | 0.5901 | 0 |
| T7B9 (Gap) T3 | 0.5938 | 0 |

*p-values given as 0 mean < $10^{-307}$

Kruskal-Wallis test statistic: 4014
p-value: 0

Supplementary Figure 11: Quantile-Quantile (QQ) plots, and the Kolmogorov-Smirnov test for each independent repeat of the seven duplex constructs for the dye-pair Top -7 Bottom +9. Kruskal-Wallis test result to confirm that within all the groups there were samples originating from different distributions.

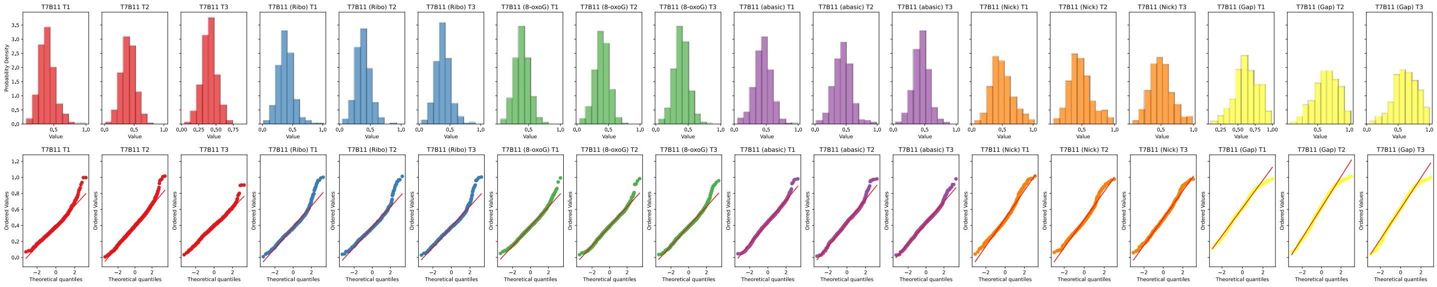

| Construct | Kolmogorov–Smirnov test statistic | p-value |
|---|---|---|
| T7B11 T1 | 0.5464 | 8.838e-264 |
| T7B11 T2 | 0.5403 | 0 |
| T7B11 T3 | 0.5437 | 0 |
| T7B11 (Ribo) T1 | 0.5312 | 0 |
| T7B11 (Ribo) T2 | 0.5374 | 0 |
| T7B11 (Ribo) T3 | 0.5373 | 2.941e-304 |
| T7B11 (8-oxoG) T1 | 0.5407 | 0 |
| T7B11 (8-oxoG) T2 | 0.533 | 0 |
| T7B11 (8-oxoG) T3 | 0.5374 | 0 |
| T7B11 (abasic) T1 | 0.5375 | 0 |
| T7B11 (abasic) T2 | 0.5432 | 4.532e-293 |
| T7B11 (abasic) T3 | 0.5464 | 0 |
| T7B11 (Nick) T1 | 0.5549 | 3.289e-210 |
| T7B11 (Nick) T2 | 0.5434 | 6.817e-139 |
| T7B11 (Nick) T3 | 0.5515 | 9.672e-198 |
| T7B11 (Gap) T1 | 0.5816 | 5.039e-128 |
| T7B11 (Gap) T2 | 0.5691 | 5.166e-217 |
| T7B11 (Gap) T3 | 0.5648 | 1.119e-132 |

*p-values given as 0 mean $< 10^{-307}$

Kruskal-Wallis test statistic: 2730
p-value: 0

Supplementary Figure 12: Quantile-Quantile (QQ) plots, and the Kolmogorov-Smirnov test for each independent repeat of the seven duplex constructs for the dye-pair Top -7 Bottom +11. Kruskal-Wallis test result to confirm that within all the groups there were samples originating from different distributions.

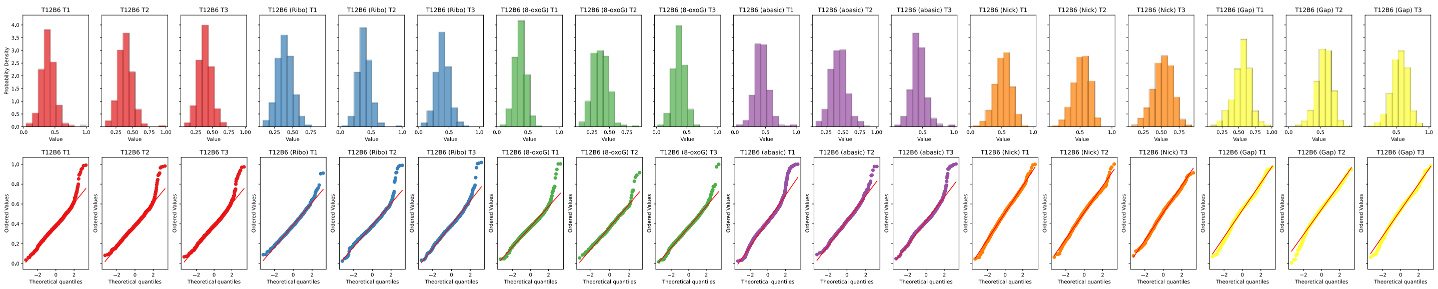

| Construct | Kolmogorov–Smirnov test statistic | p-value |
|---|---|---|
| T12B6 T1 | 0.5451 | 0 |
| T12B6 T2 | 0.5499 | 0 |
| T12B6 T3 | 0.5475 | 0 |
| T12B6 (Ribo) T1 | 0.5507 | 6.9e-261 |
| T12B6 (Ribo) T2 | 0.5539 | 4.976e-285 |
| T12B6 (Ribo) T3 | 0.5422 | 1.135e-257 |
| T12B6 (8-oxoG) T1 | 0.5479 | 0 |
| T12B6 (8-oxoG) T2 | 0.5419 | 2.703e-176 |
| T12B6 (8-oxoG) T3 | 0.5457 | 0 |
| T12B6 (abasic) T1 | 0.5524 | 0 |
| T12B6 (abasic) T2 | 0.5533 | 6.657e-268 |
| T12B6 (abasic) T3 | 0.5534 | 0 |
| T12B6 (Nick) T1 | 0.5676 | 0 |
| T12B6 (Nick) T2 | 0.5706 | 0 |
| T12B6 (Nick) T3 | 0.5638 | 0 |
| T12B6 (Gap) T1 | 0.5685 | 0 |
| T12B6 (Gap) T2 | 0.5805 | 0 |
| T12B6 (Gap) T3 | 0.5754 | 0 |

*p-values given as 0 mean $< 10^{-307}$

Kruskal-Wallis test statistic: 6372
p-value: 0

Supplementary Figure 13: Quantile-Quantile (QQ) plots, and the Kolmogorov-Smirnov test for each independent repeat of the seven duplex constructs for the dye-pair Top -12 Bottom +6. Kruskal-Wallis test result to confirm that within all the groups there were samples originating from different distributions.

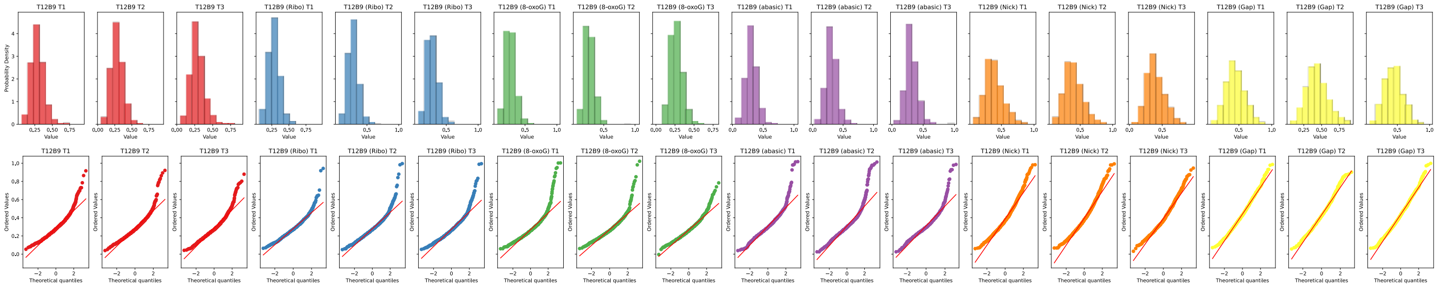

| Construct | Kolmogorov–Smirnov test statistic | p-value |
|---|---|---|
| T12B9 T1 | 0.5363 | 0 |
| T12B9 T2 | 0.5331 | 0 |
| T12B9 T3 | 0.526 | 0 |
| T12B9 (Ribo) T1 | 0.5357 | 0 |
| T12B9 (Ribo) T2 | 0.5333 | 0 |
| T12B9 (Ribo) T3 | 0.5311 | 0 |
| T12B9 (8-oxoG) T1 | 0.5332 | 0 |
| T12B9 (8-oxoG) T2 | 0.5354 | 0 |
| T12B9 (8-oxoG) T3 | 0.5324 | 0 |
| T12B9 (abasic) T1 | 0.5294 | 0 |
| T12B9 (abasic) T2 | 0.5273 | 0 |
| T12B9 (abasic) T3 | 0.532 | 3.577e-240 |
| T12B9 (Nick) T1 | 0.5401 | 0 |
| T12B9 (Nick) T2 | 0.539 | 0 |
| T12B9 (Nick) T3 | 0.5433 | 0 |
| T12B9 (Gap) T1 | 0.5485 | 0 |
| T12B9 (Gap) T2 | 0.5441 | 1.771e-275 |
| T12B9 (Gap) T3 | 0.5484 | 0 |

*p-values given as 0 mean < $10^{-307}$

Kruskal-Wallis test statistic: 5735
p-value: 0

Supplementary Figure 14: Quantile-Quantile (QQ) plots, and the Kolmogorov-Smirnov test for each independent repeat of the seven duplex constructs for the dye-pair Top -12 Bottom +9. Kruskal-Wallis test result to confirm that within all the groups there were samples originating from different distributions.

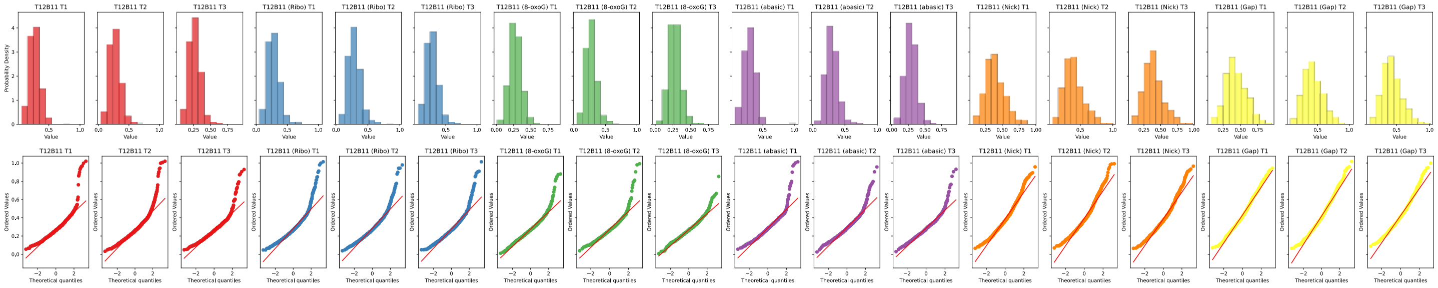

| Construct | Kolmogorov–Smirnov test statistic | p-value |
|---|---|---|
| T12B11 T1 | 0.5334 | 0 |
| T12B11 T2 | 0.5236 | 0 |
| T12B11 T3 | 0.5299 | 0 |
| T12B11 (Ribo) T1 | 0.5279 | 0 |
| T12B11 (Ribo) T2 | 0.5287 | 0 |
| T12B11 (Ribo) T3 | 0.5245 | 0 |
| T12B11 (8-oxoG) T1 | 0.5223 | 0 |
| T12B11 (8-oxoG) T2 | 0.5305 | 1.795e-243 |
| T12B11 (8-oxoG) T3 | 0.5252 | 4.073e-318 |
| T12B11 (abasic) T1 | 0.5331 | 3.819e-287 |
| T12B11 (abasic) T2 | 0.5348 | 2.661e-302 |
| T12B11 (abasic) T3 | 0.5283 | 1.33e-230 |
| T12B11 (Nick) T1 | 0.5395 | 3.559e-276 |
| T12B11 (Nick) T2 | 0.541 | 6.226e-268 |
| T12B11 (Nick) T3 | 0.5377 | 1.708e-308 |
| T12B11 (Gap) T1 | 0.538 | 3.246e-274 |
| T12B11 (Gap) T2 | 0.5393 | 0 |
| T12B11 (Gap) T3 | 0.5495 | 1.006e-251 |

*p-values given as 0 mean < $10^{-307}$

Kruskal-Wallis test statistic: 4260
p-value: 0

Supplementary Figure 15: Quantile-Quantile (QQ) plots, and the Kolmogorov-Smirnov test for each independent repeat of the seven duplex constructs for the dye-pair Top -12 Bottom +11. Kruskal-Wallis test result to confirm that within all the groups there were samples originating from different distributions.

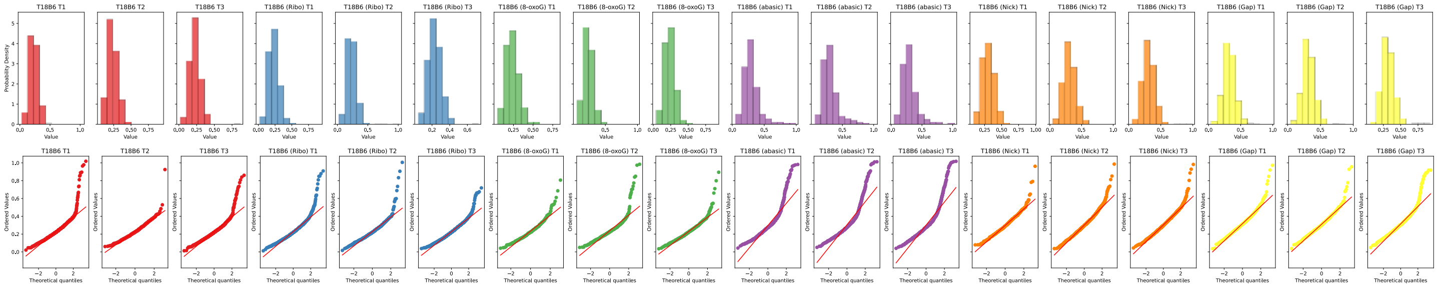

| Construct | Kolmogorov–Smirnov test statistic | p-value |
|---|---|---|
| T18B6 T1 | 0.5225 | 0 |
| T18B6 T2 | 0.5251 | 3.681e-201 |
| T18B6 T3 | 0.5204 | 0 |
| T18B6 (Ribo) T1 | 0.5216 | 0 |
| T18B6 (Ribo) T2 | 0.5213 | 6.975e-301 |
| T18B6 (Ribo) T3 | 0.5215 | 0 |
| T18B6 (8-oxoG) T1 | 0.5241 | 1.219e-149 |
| T18B6 (8-oxoG) T2 | 0.5243 | 4.467e-214 |
| T18B6 (8-oxoG) T3 | 0.5235 | 2.094e-252 |
| T18B6 (abasic) T1 | 0.5318 | 9.961e-305 |
| T18B6 (abasic) T2 | 0.5333 | 0 |
| T18B6 (abasic) T3 | 0.5267 | 0 |
| T18B6 (Nick) T1 | 0.5365 | 1.637e-275 |
| T18B6 (Nick) T2 | 0.5301 | 0 |
| T18B6 (Nick) T3 | 0.5332 | 0 |
| T18B6 (Gap) T1 | 0.5384 | 0 |
| T18B6 (Gap) T2 | 0.5364 | 0 |
| T18B6 (Gap) T3 | 0.5417 | 0 |

*p-values given as 0 mean < $10^{-307}$

Kruskal-Wallis test statistic: 4748
p-value: 0

Supplementary Figure 16: Quantile-Quantile (QQ) plots, and the Kolmogorov-Smirnov test for each independent repeat of the seven duplex constructs for the dye-pair Top -18 Bottom +6. Kruskal-Wallis test result to confirm that within all the groups there were samples originating from different distributions.

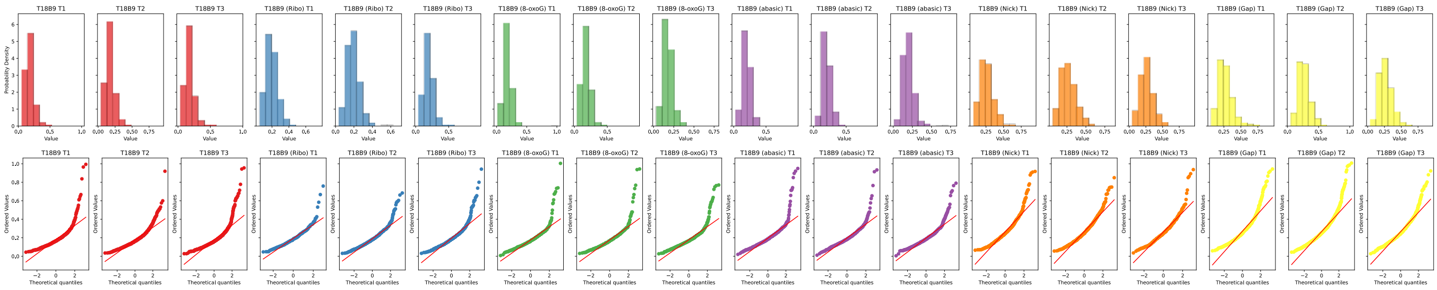

| Construct | Kolmogorov–Smirnov test statistic | p-value |
|---|---|---|
| T18B9 T1 | 0.5186 | 7.738e-227 |
| T18B9 T2 | 0.5154 | 0 |
| T18B9 T3 | 0.5176 | 0 |
| T18B9 (Ribo) T1 | 0.5187 | 1.314e-125 |
| T18B9 (Ribo) T2 | 0.5151 | 0 |
| T18B9 (Ribo) T3 | 0.5154 | 0 |
| T18B9 (8-oxoG) T1 | 0.5142 | 1.191e-237 |
| T18B9 (8-oxoG) T2 | 0.5135 | 1.621e-236 |
| T18B9 (8-oxoG) T3 | 0.5151 | 5.028e-261 |
| T18B9 (abasic) T1 | 0.5206 | 0 |
| T18B9 (abasic) T2 | 0.515 | 6.448e-226 |
| T18B9 (abasic) T3 | 0.5214 | 5.698e-295 |
| T18B9 (Nick) T1 | 0.5268 | 2.65e-247 |
| T18B9 (Nick) T2 | 0.525 | 2.943e-273 |
| T18B9 (Nick) T3 | 0.5248 | 3.405e-184 |
| T18B9 (Gap) T1 | 0.5314 | 0 |
| T18B9 (Gap) T2 | 0.5272 | 0 |
| T18B9 (Gap) T3 | 0.5277 | 0 |

*p-values given as 0 mean $< 10^{-307}$

Kruskal-Wallis test statistic: 4240
p-value: 0

Supplementary Figure 17: Quantile-Quantile (QQ) plots, and the Kolmogorov-Smirnov test for each independent repeat of the seven duplex constructs for the dye-pair Top -18 Bottom +9. Kruskal-Wallis test result to confirm that within all the groups there were samples originating from different distributions.

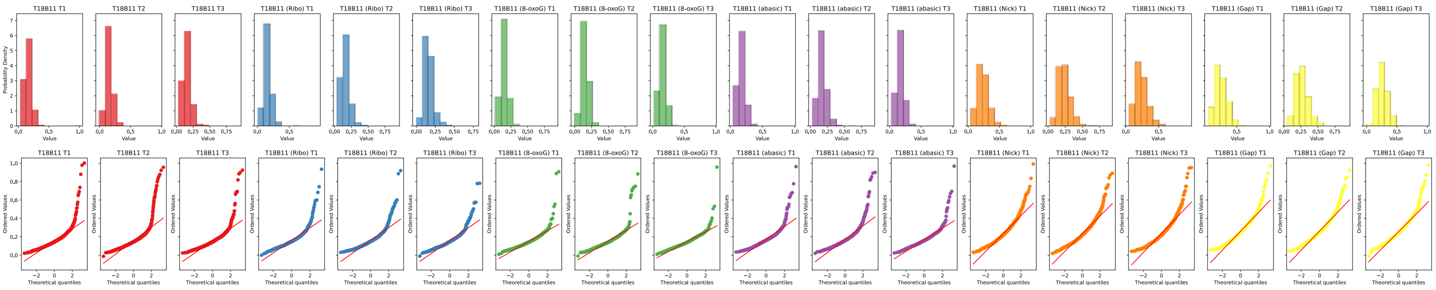

| Construct | Kolmogorov–Smirnov test statistic | p-value |
|---|---|---|
| T18B11 T1 | 0.5125 | 0 |
| T18B11 T2 | 0.5107 | 0 |
| T18B11 T3 | 0.5123 | 0 |
| T18B11 (Ribo) T1 | 0.5116 | 1.182e-284 |
| T18B11 (Ribo) T2 | 0.5145 | 0 |
| T18B11 (Ribo) T3 | 0.5095 | 2.648e-258 |
| T18B11 (8-oxoG) T1 | 0.5136 | 2.506e-260 |
| T18B11 (8-oxoG) T2 | 0.5104 | 4.507e-309 |
| T18B11 (8-oxoG) T3 | 0.5098 | 2.688e-247 |
| T18B11 (abasic) T1 | 0.5149 | 1.74e-283 |
| T18B11 (abasic) T2 | 0.5152 | 0 |
| T18B11 (abasic) T3 | 0.5127 | 5.311e-314 |
| T18B11 (Nick) T1 | 0.5206 | 0 |
| T18B11 (Nick) T2 | 0.5188 | 0 |
| T18B11 (Nick) T3 | 0.5206 | 0 |
| T18B11 (Gap) T1 | 0.5241 | 0 |
| T18B11 (Gap) T2 | 0.5228 | 2.205e-306 |
| T18B11 (Gap) T3 | 0.5236 | 0 |

*p-values given as 0 mean < $10^{-307}$

Kruskal-Wallis test statistic: 6230
p-value: 0

Supplementary Figure 18: Quantile-Quantile (QQ) plots, and the Kolmogorov-Smirnov test for each independent repeat of the seven duplex constructs for the dye-pair Top -18 Bottom +11. Kruskal-Wallis test result to confirm that within all the groups there were samples originating from different distributions.

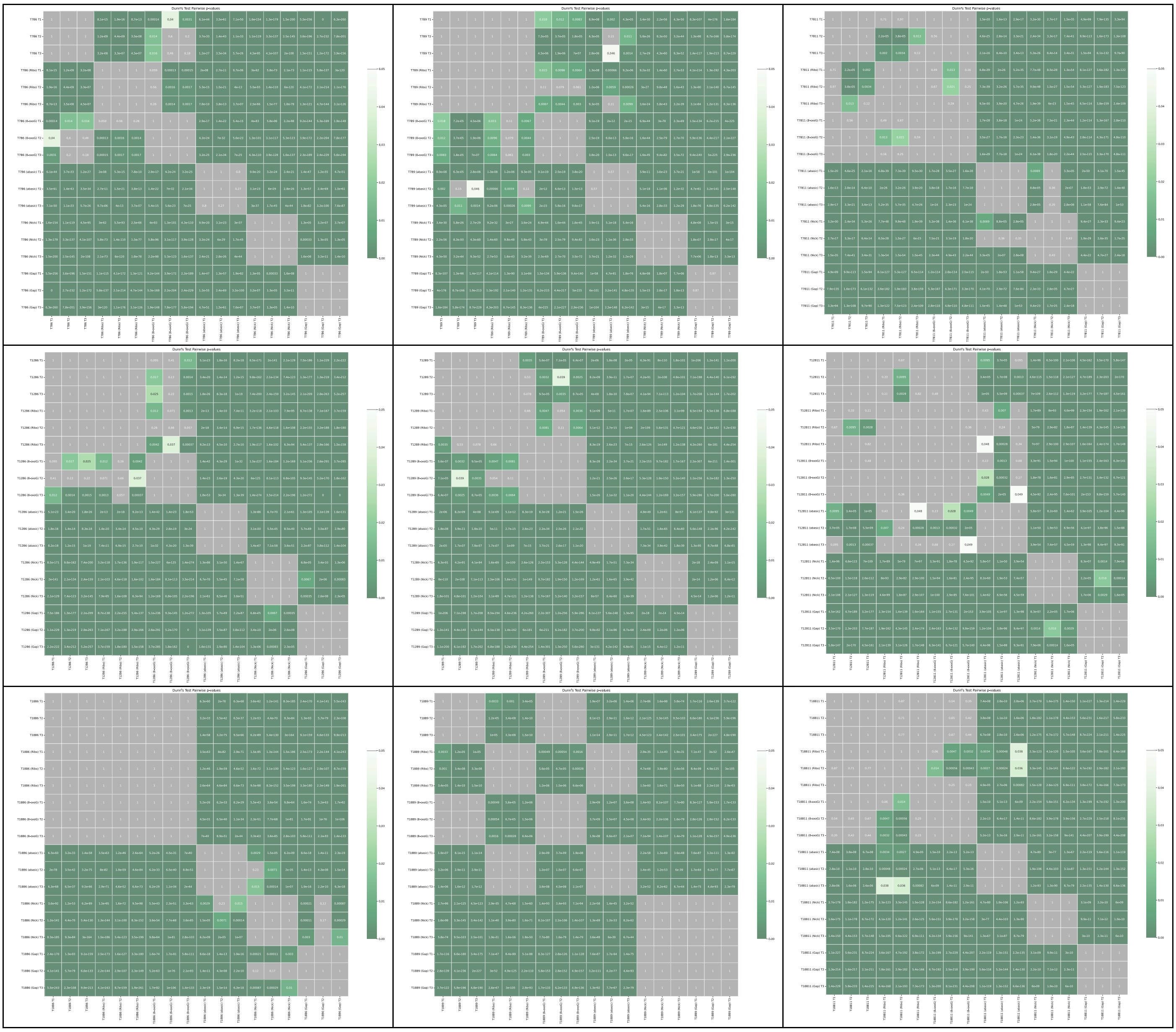

Supplementary Figure 19: Dunn’s Tests for each of the repeats for each of the labelling positions for each of the different types of damage.

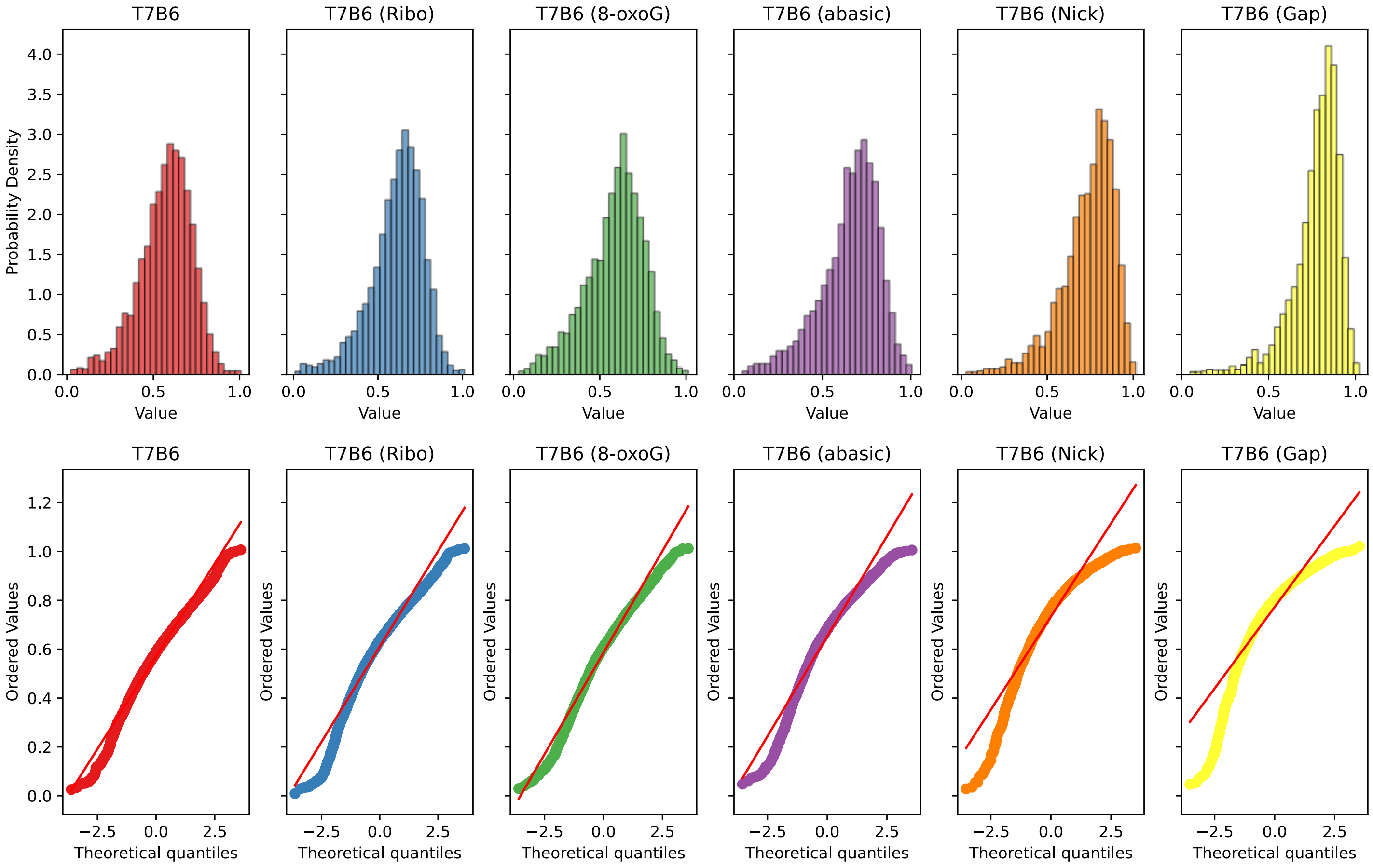

| Construct | Kolmogorov–Smirnov test statistic | p-value* |
|---|---|---|
| T7B6 | 0.5663 | 0 |
| T7B6 (Ribo) | 0.5734 | 0 |
| T7B6 (8-oxoG) | 0.5565 | 0 |
| T7B6 (abasic) | 0.5771 | 0 |
| T7B6 (Nick) | 0.6143 | 0 |
| T7B6 (Gap) | 0.6424 | 0 |

*p-values given as 0 mean < $10^{-307}$

Kruskal-Wallis test statistic: 5041
p-value: 0

Supplementary Figure 20: Quantile-Quantile (QQ) plots, and the Kolmogorov-Smirnov test for each combined repeats of the seven duplex constructs for the dye-pair Top -7 Bottom +6. Kruskal-Wallis test result to confirm that within all the groups there were samples originating from different distributions.

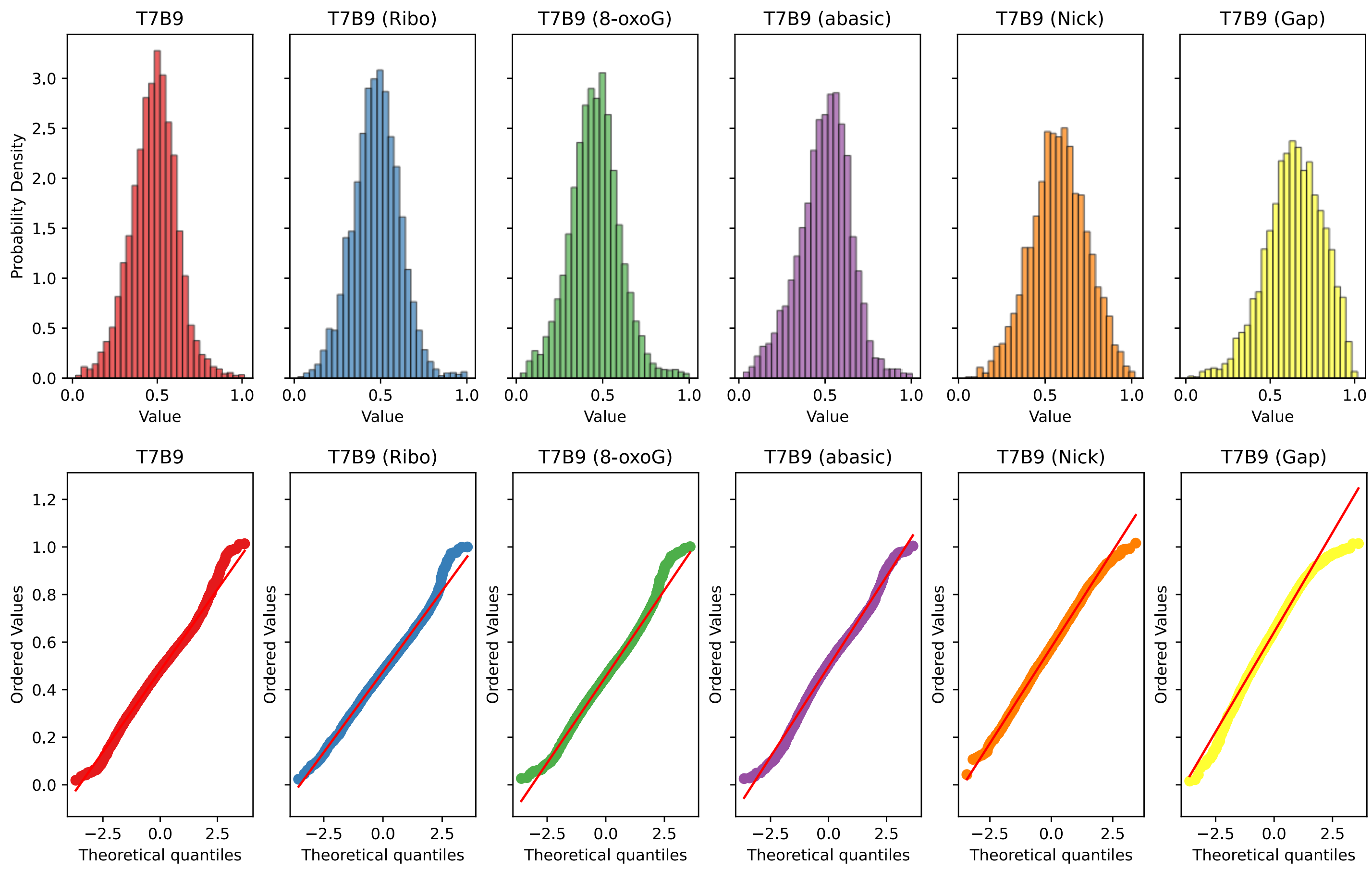

| Construct | Kolmogorov–Smirnov test statistic | p-value* |
|---|---|---|
| T7B9 | 0.5551 | 0 |
| T7B9 (Ribo) | 0.5578 | 0 |
| T7B9 (8-oxoG) | 0.5394 | 0 |
| T7B9 (abasic) | 0.5414 | 0 |
| T7B9 (Nick) | 0.5746 | 0 |
| T7B9 (Gap) | 0.5892 | 0 |

*p-values given as 0 mean $< 10^{-307}$

Kruskal-Wallis test statistic: 3998
p-value: 0

Supplementary Figure 21: Quantile-Quantile (QQ) plots, and the Kolmogorov-Smirnov test for each combined repeats of the seven duplex constructs for the dye-pair Top -7 Bottom +9. Kruskal-Wallis test result to confirm that within all the groups there were samples originating from different distributions.

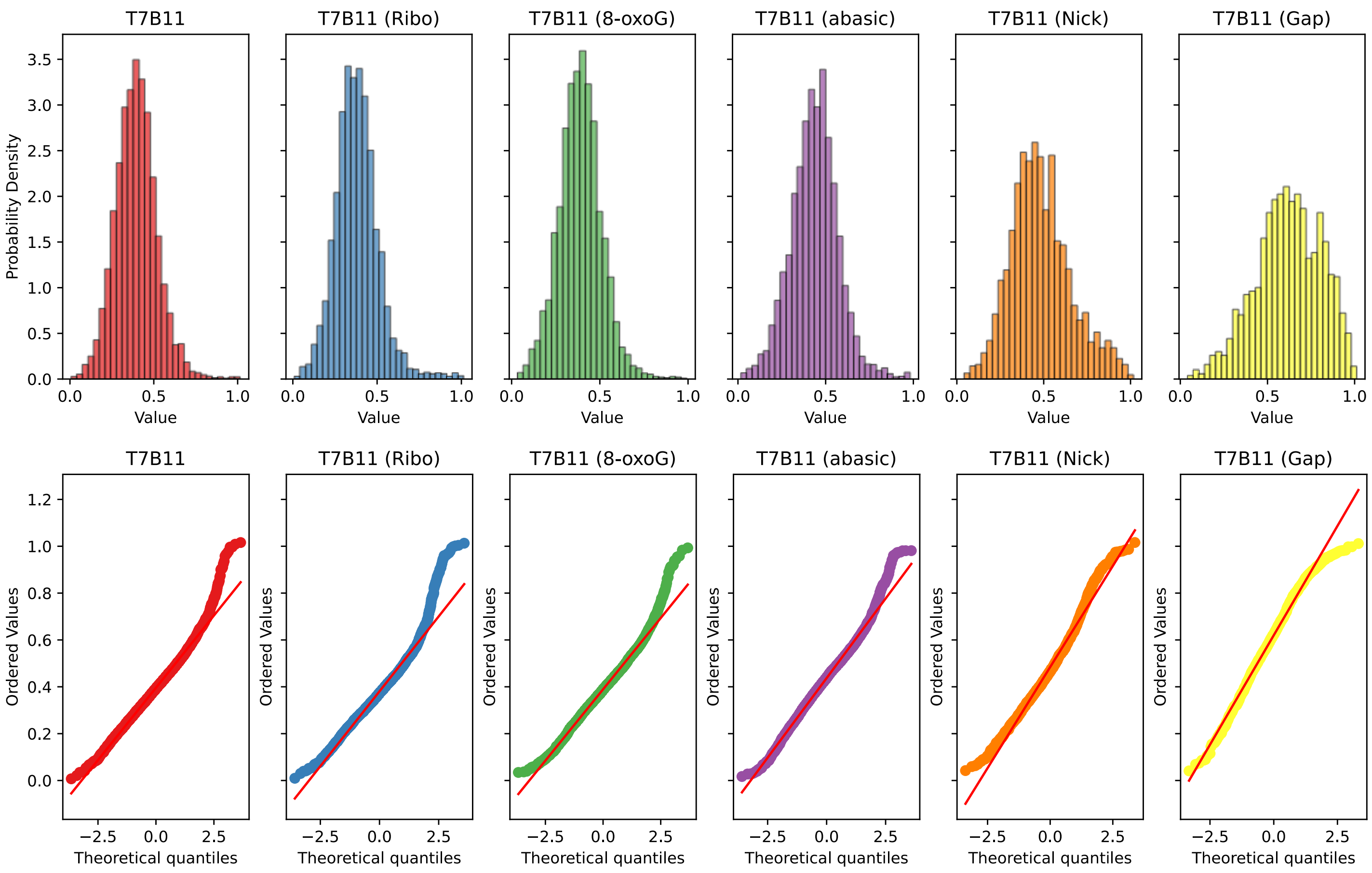

| Construct | Kolmogorov–Smirnov test statistic | p-value* |
|---|---|---|
| T7B11 | 0.5408 | 0 |
| T7B11 (Ribo) | 0.5342 | 0 |
| T7B11 (8-oxoG) | 0.5362 | 0 |
| T7B11 (abasic) | 0.5421 | 0 |
| T7B11 (Nick) | 0.5483 | 0 |
| T7B11 (Gap) | 0.5684 | 0 |

*p-values given as 0 mean < $10^{-307}$

Kruskal-Wallis test statistic: 2714
p-value: 0

Supplementary Figure 22: Quantile-Quantile (QQ) plots, and the Kolmogorov-Smirnov test for each combined repeats of the seven duplex constructs for the dye-pair Top -7 Bottom +11. Kruskal-Wallis test result to confirm that within all the groups there were samples originating from different distributions.

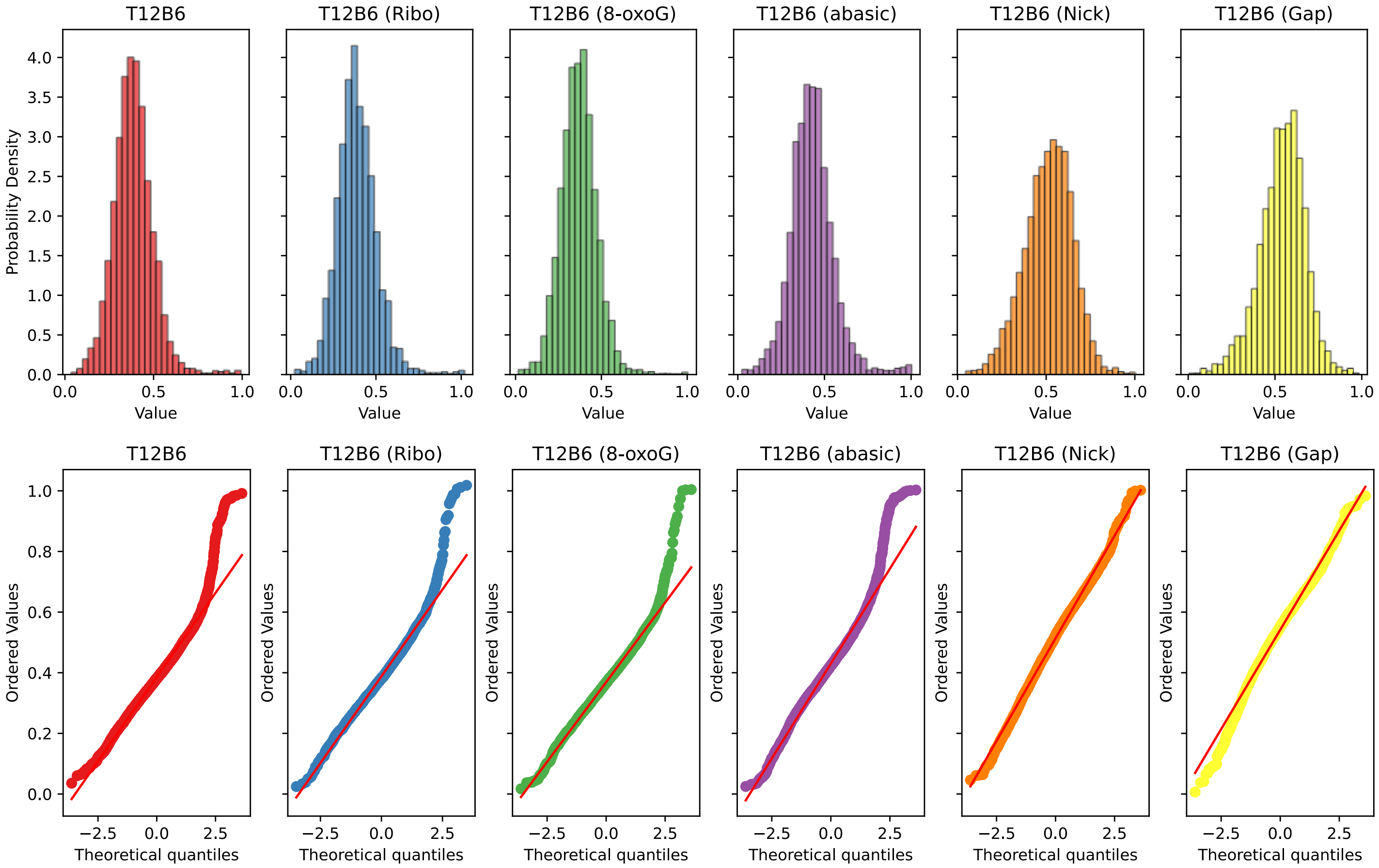

| Construct | Kolmogorov–Smirnov test statistic | p-value* |
|---|---|---|
| T12B6 | 0.546 | 0 |
| T12B6 (Ribo) | 0.5473 | 0 |
| T12B6 (8-oxoG) | 0.5452 | 0 |
| T12B6 (abasic) | 0.5514 | 0 |
| T12B6 (Nick) | 0.5664 | 0 |
| T12B6 (Gap) | 0.5741 | 0 |

*p-values given as 0 mean < $10^{-307}$

Kruskal-Wallis test statistic: 6372
p-value: 0

Supplementary Figure 23: Quantile-Quantile (QQ) plots, and the Kolmogorov-Smirnov test for each combined repeats of the seven duplex constructs for the dye-pair Top -12 Bottom +6. Kruskal-Wallis test result to confirm that within all the groups there were samples originating from different distributions.

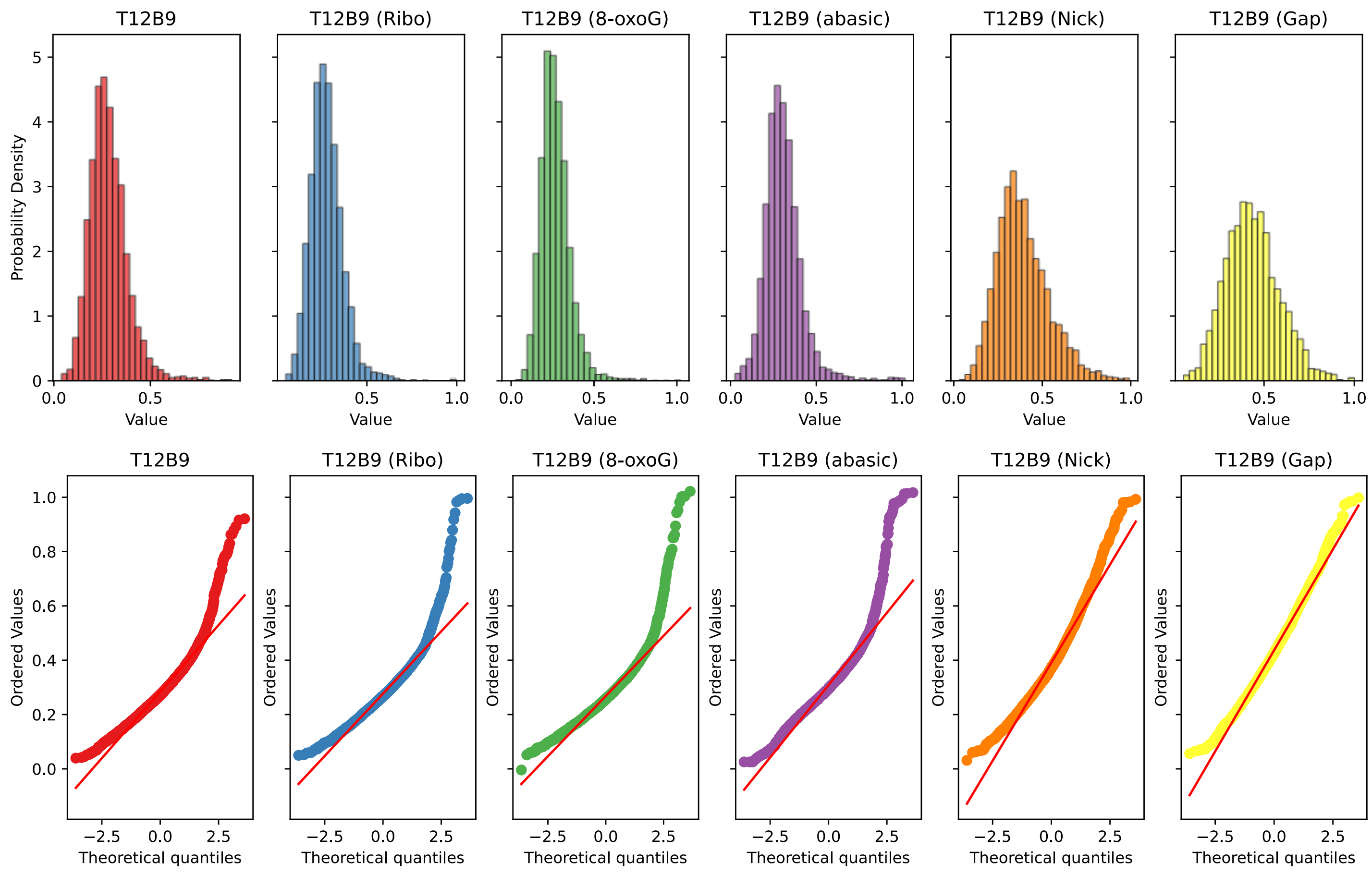

| Construct | Kolmogorov–Smirnov test statistic | p-value* |
|---|---|---|
| T12B9 | 0.5308 | 0 |
| T12B9 (Ribo) | 0.5324 | 0 |
| T12B9 (8-oxoG) | 0.5328 | 0 |
| T12B9 (abasic) | 0.5281 | 0 |
| T12B9 (Nick) | 0.5399 | 0 |
| T12B9 (Gap) | 0.5468 | 0 |

*p-values given as 0 mean $< 10^{-307}$

Kruskal-Wallis test statistic: 5735
p-value: 0

Supplementary Figure 24: Quantile-Quantile (QQ) plots, and the Kolmogorov-Smirnov test for each combined repeats of the seven duplex constructs for the dye-pair Top -12 Bottom +9. Kruskal-Wallis test result to confirm that within all the groups there were samples originating from different distributions.

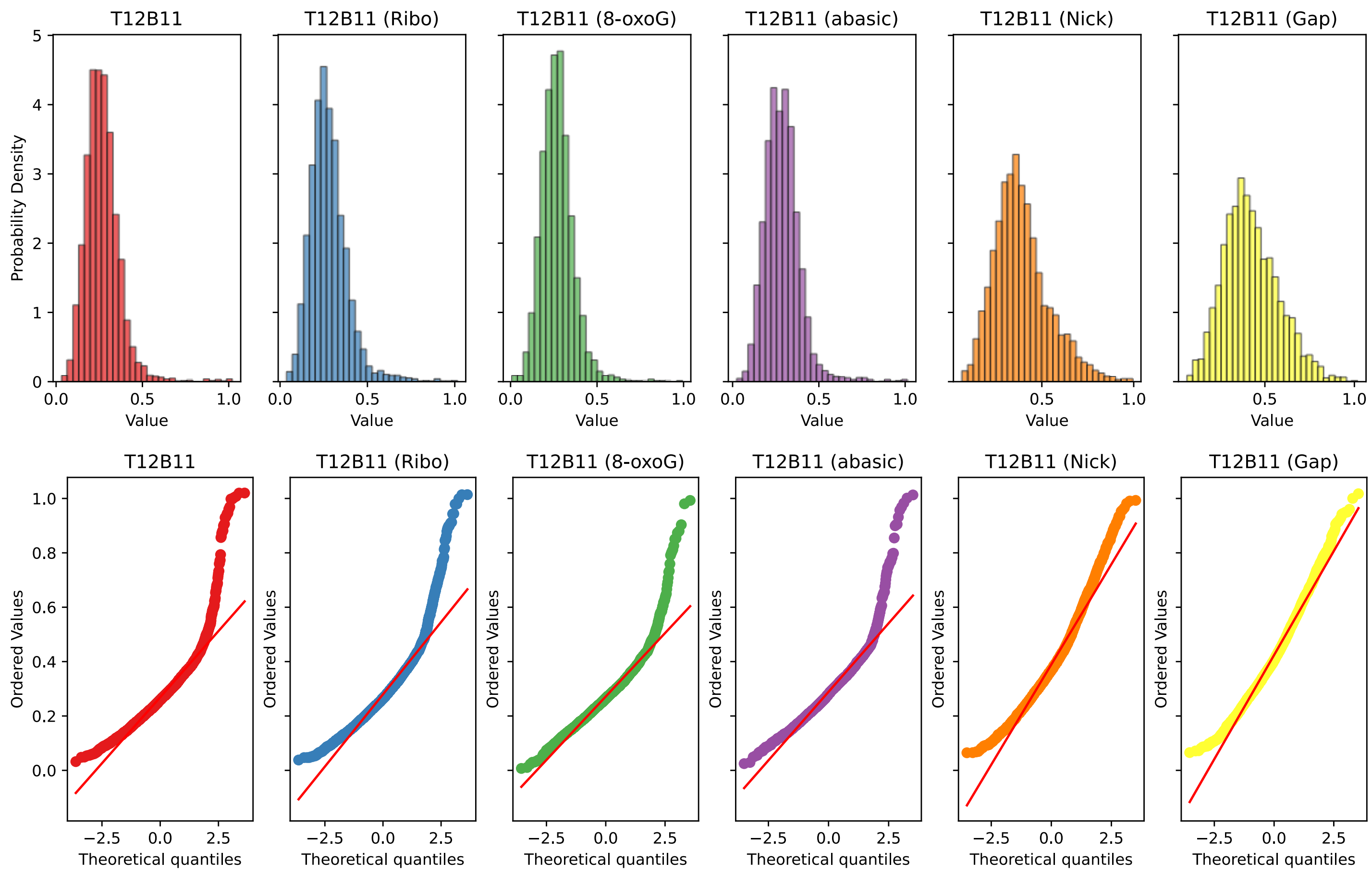

| Construct | Kolmogorov–Smirnov test statistic | p-value* |
|---|---|---|
| T12B11 | 0.5277 | 0 |
| T12B11 (Ribo) | 0.5268 | 0 |
| T12B11 (8-oxoG) | 0.5243 | 0 |
| T12B11 (abasic) | 0.5314 | 0 |
| T12B11 (Nick) | 0.5388 | 0 |
| T12B11 (Gap) | 0.5408 | 0 |

*p-values given as 0 mean < $10^{-307}$

Kruskal-Wallis test statistic: 4260
p-value: 0

Supplementary Figure 25: Quantile-Quantile (QQ) plots, and the Kolmogorov-Smirnov test for each combined repeats of the seven duplex constructs for the dye-pair Top -12 Bottom +11. Kruskal-Wallis test result to confirm that within all the groups there were samples originating from different distributions.

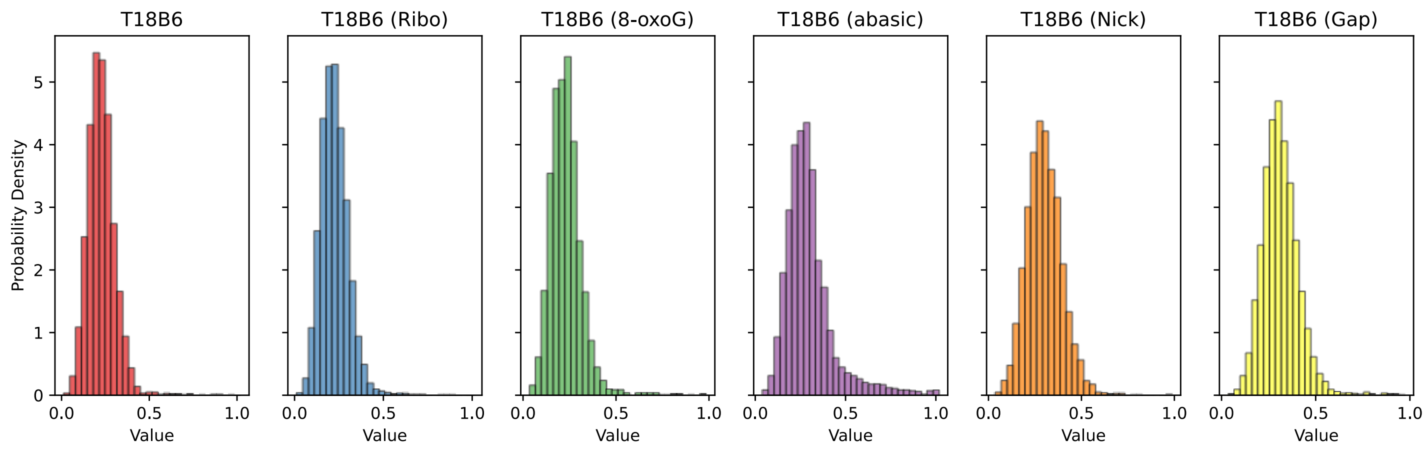

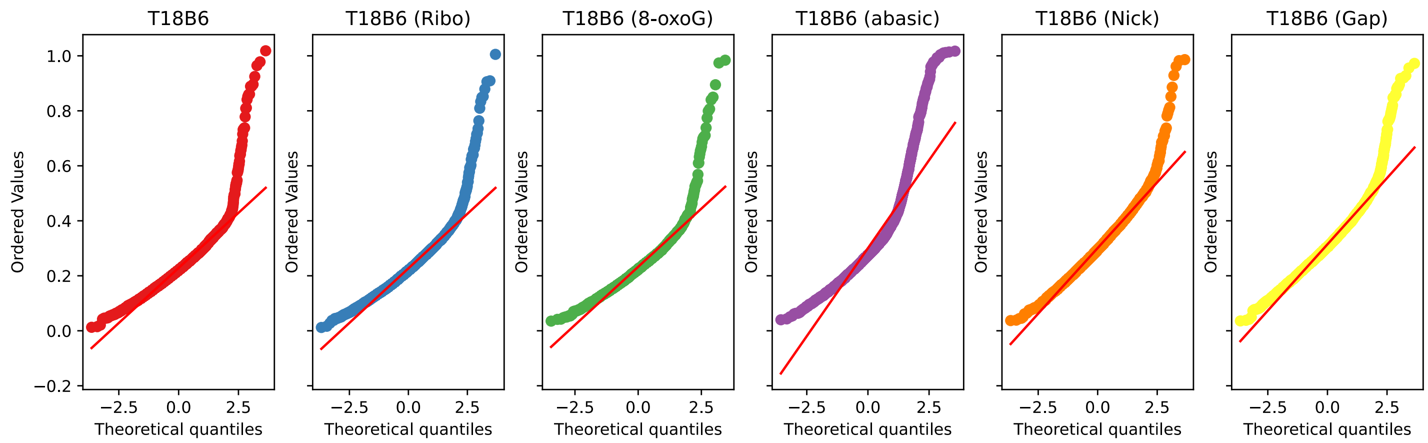

| Construct | Kolmogorov–Smirnov test statistic | p-value* |
|---|---|---|
| T18B6 | 0.5216 | 0 |
| T18B6 (Ribo) | 0.5211 | 0 |
| T18B6 (8-oxoG) | 0.5229 | 0 |
| T18B6 (abasic) | 0.5301 | 0 |
| T18B6 (Nick) | 0.532 | 0 |
| T18B6 (Gap) | 0.5391 | 0 |

*p-values given as 0 mean < $10^{-307}$

Kruskal-Wallis test statistic: 4738
p-value: 0

Supplementary Figure 26: Quantile-Quantile (QQ) plots, and the Kolmogorov-Smirnov test for each combined repeats of the seven duplex constructs for the dye-pair Top -18 Bottom +6. Kruskal-Wallis test result to confirm that within all the groups there were samples originating from different distributions.

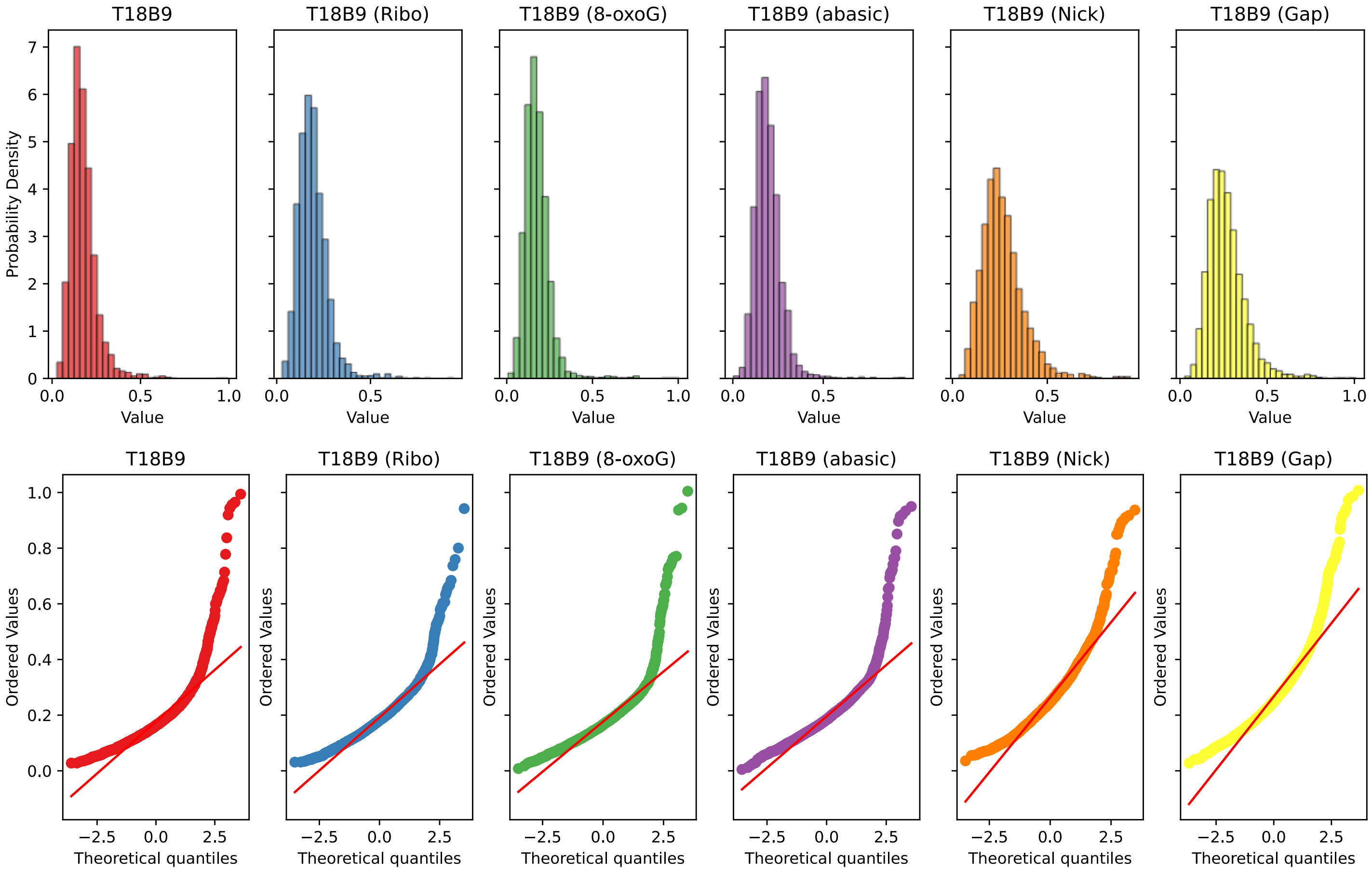

| Construct | Kolmogorov–Smirnov test statistic | p-value* |
|---|---|---|
| T18B9 | 0.5165 | 0 |
| T18B9 (Ribo) | 0.5153 | 0 |
| T18B9 (8-oxoG) | 0.5136 | 0 |
| T18B9 (abasic) | 0.5182 | 0 |
| T18B9 (Nick) | 0.5246 | 0 |
| T18B9 (Gap) | 0.5279 | 0 |

*p-values given as 0 mean < $10^{-307}$

Kruskal-Wallis test statistic: 4236
p-value: 0

Supplementary Figure 27: Quantile-Quantile (QQ) plots, and the Kolmogorov-Smirnov test for each combined repeats of the seven duplex constructs for the dye-pair Top -18 Bottom +9. Kruskal-Wallis test result to confirm that within all the groups there were samples originating from different distributions.

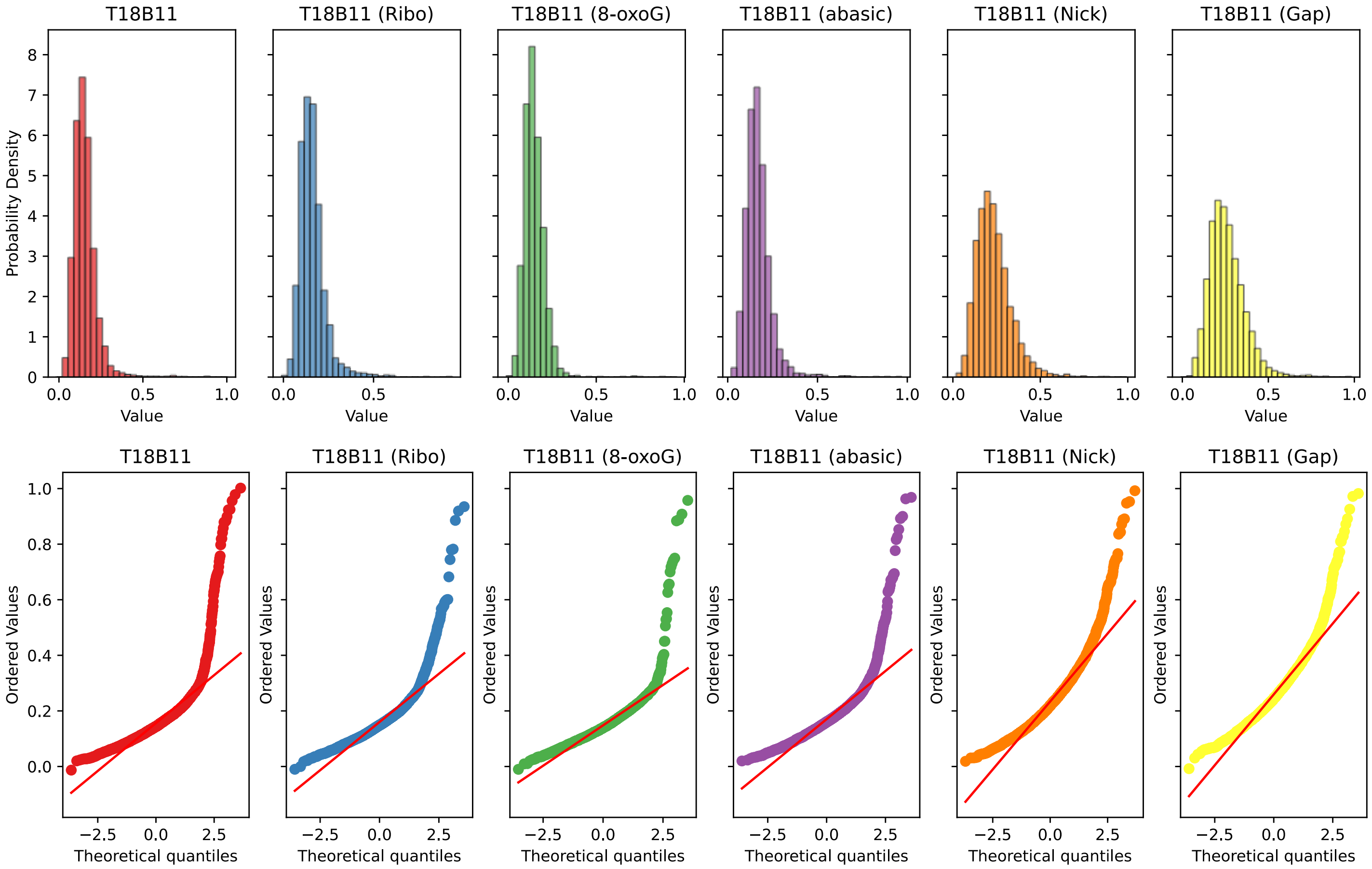

| Construct | Kolmogorov–Smirnov test statistic | p-value* |
|---|---|---|
| T18B11 | 0.511 | 0 |
| T18B11 (Ribo) | 0.5113 | 0 |
| T18B11 (8-oxoG) | 0.5104 | 0 |
| T18B11 (abasic) | 0.5142 | 0 |
| T18B11 (Nick) | 0.5188 | 0 |
| T18B11 (Gap) | 0.5231 | 0 |

*p-values given as 0 mean $< 10^{-307}$

Kruskal-Wallis test statistic: 6225
p-value: 0

Supplementary Figure 28: Quantile-Quantile (QQ) plots, and the Kolmogorov-Smirnov test for each combined repeats of the seven duplex constructs for the dye-pair Top -18 Bottom +11. Kruskal-Wallis test result to confirm that within all the groups there were samples originating from different distributions.

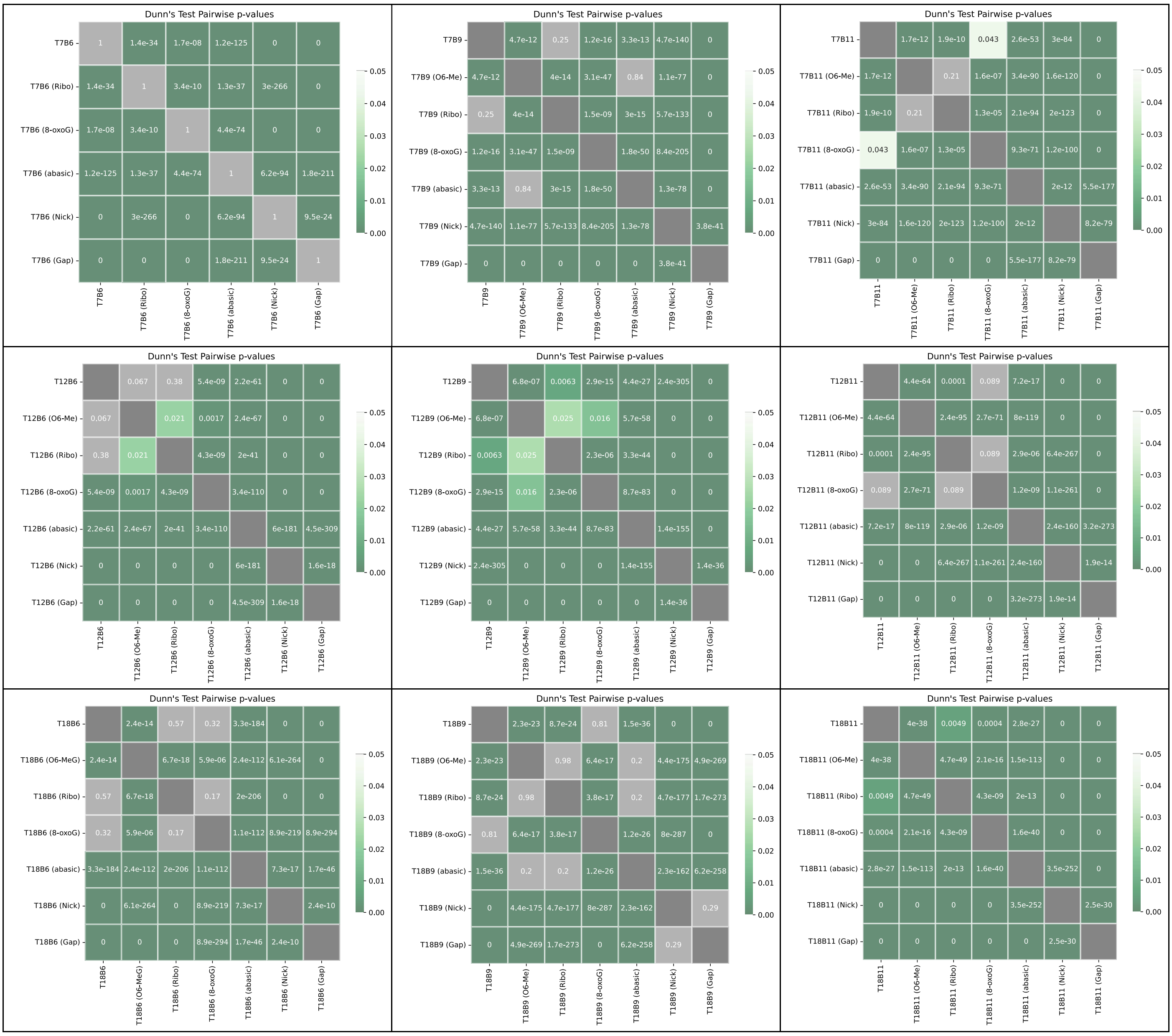

Supplementary Figure 29: Dunn's Tests results for the combined repeats for each of the labelling positions for each of the different types of damage.

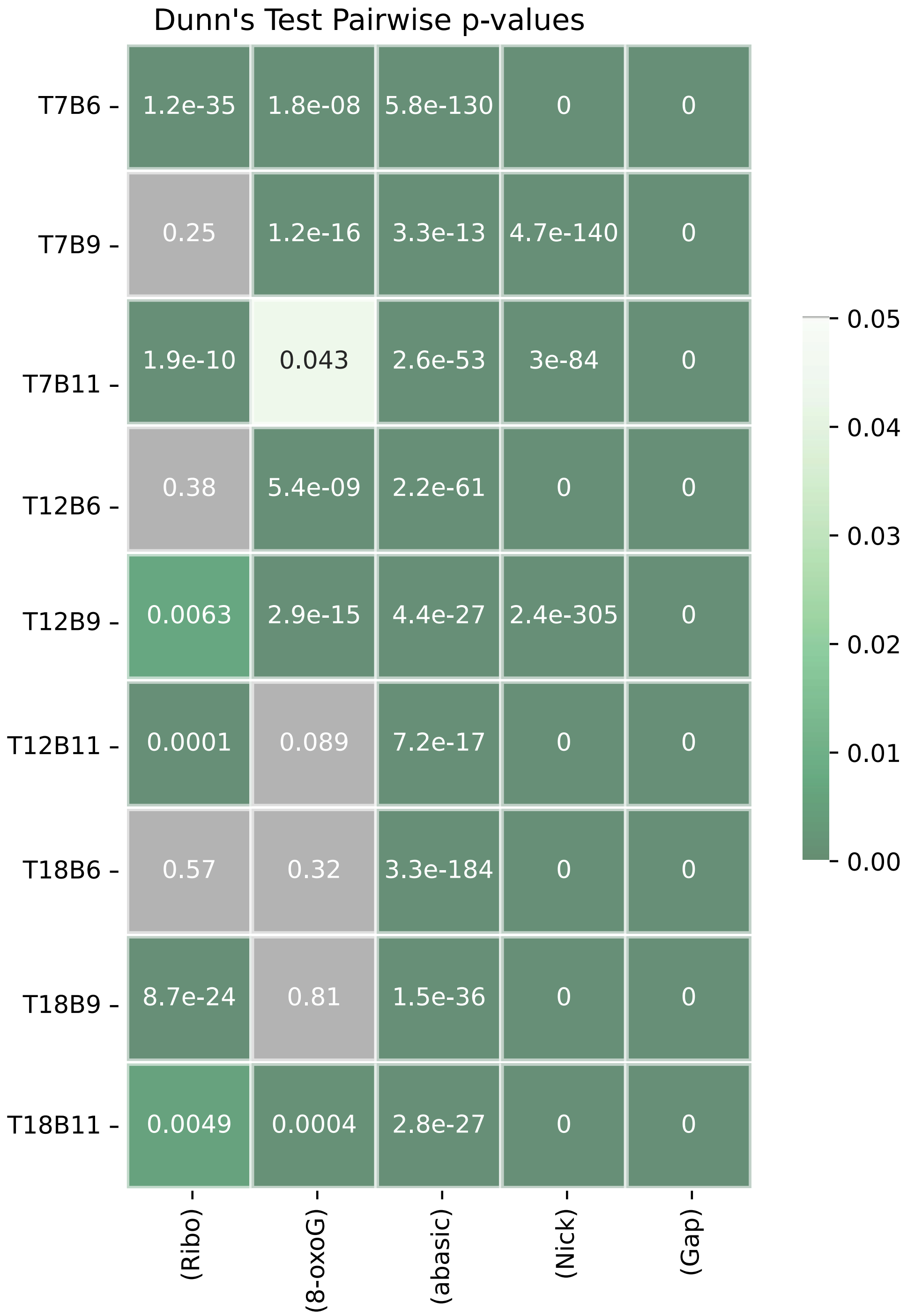

Supplementary Figure 30: All Dunn of each labelling positions with each of the different types of damage.

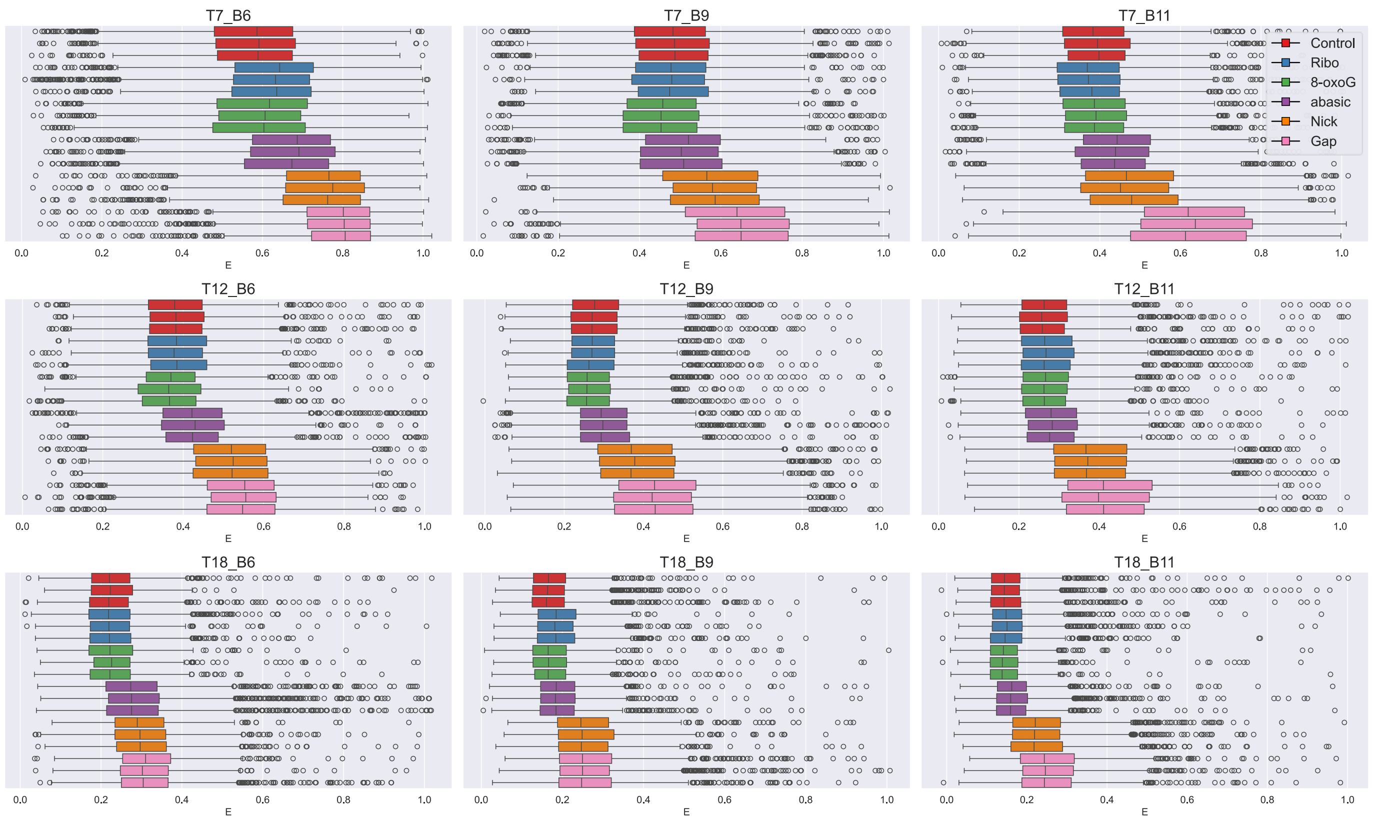

Supplementary Figure 31: Boxplots showing FRET efficiency (E) for each independent repeat of the seven duplex constructs for all dye-pair combinations.

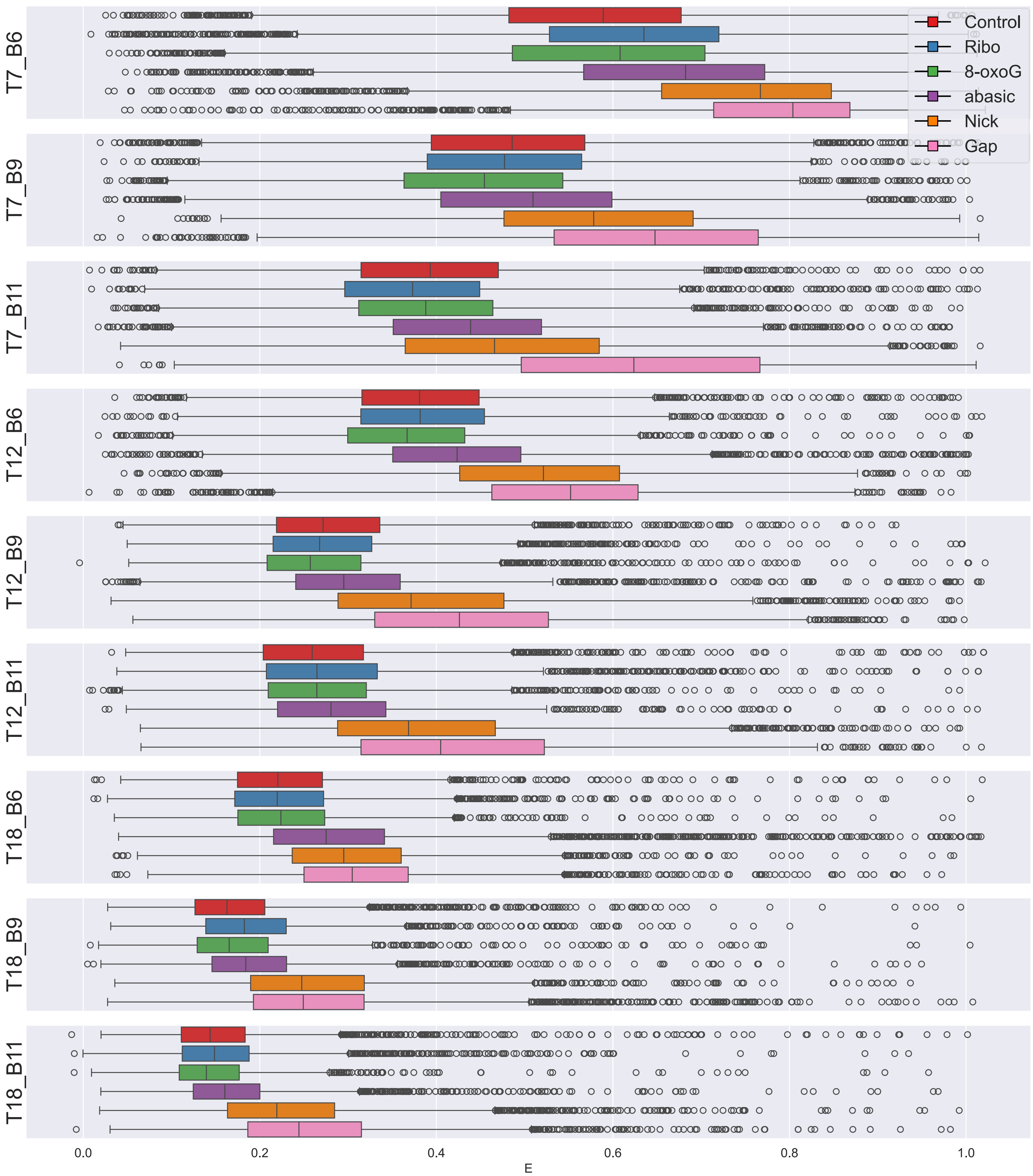

Supplementary Figure 32: Boxplots showing FRET efficiency (E) for the triplicate dataset for the seven duplex constructs for all dye-pair combination.

Example Quantum Yield calculation:

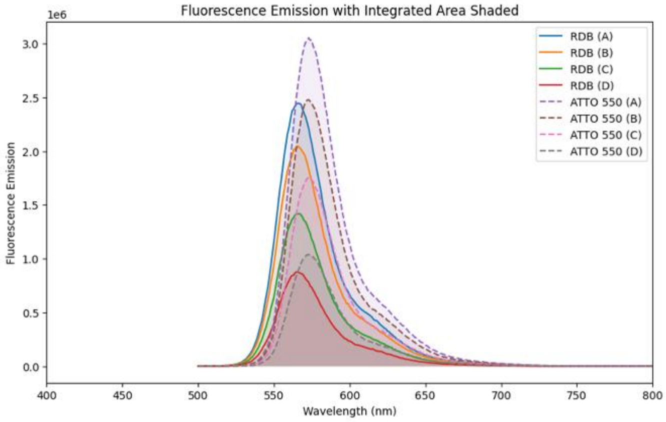

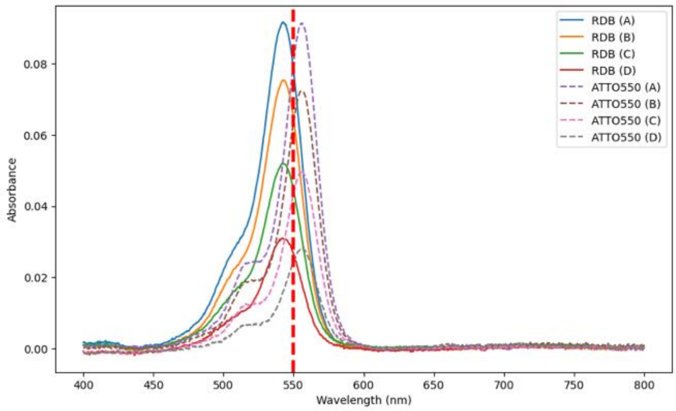

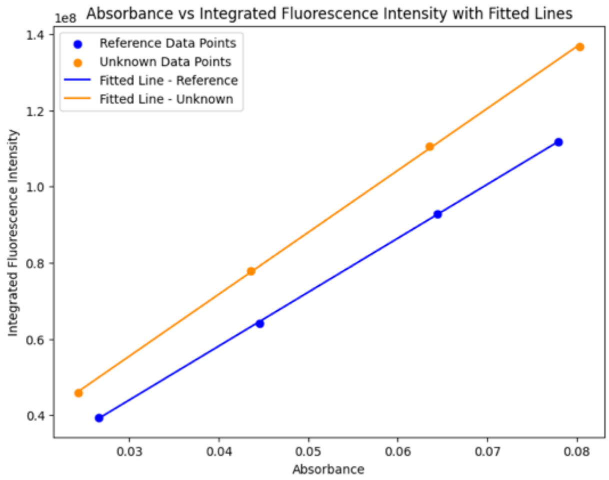

Reference data: y = 1480035464.96x + 6148414.91
R-squared: 0.9963
Unknown data: y = 1428523583.16x + 4156403.06
R-squared: 0.9993

Qr = 0.70
nr = 1.33
ns = 1.33
Qs = Qr*(ms/mr)*(ns/nr)$^2$
Unknown Sample's Quantum Yield: 0.8041

Supplementary Figure 33: Example quantum yield calculation with using RDB as reference dye to work out quantum yield of ATTO 550.

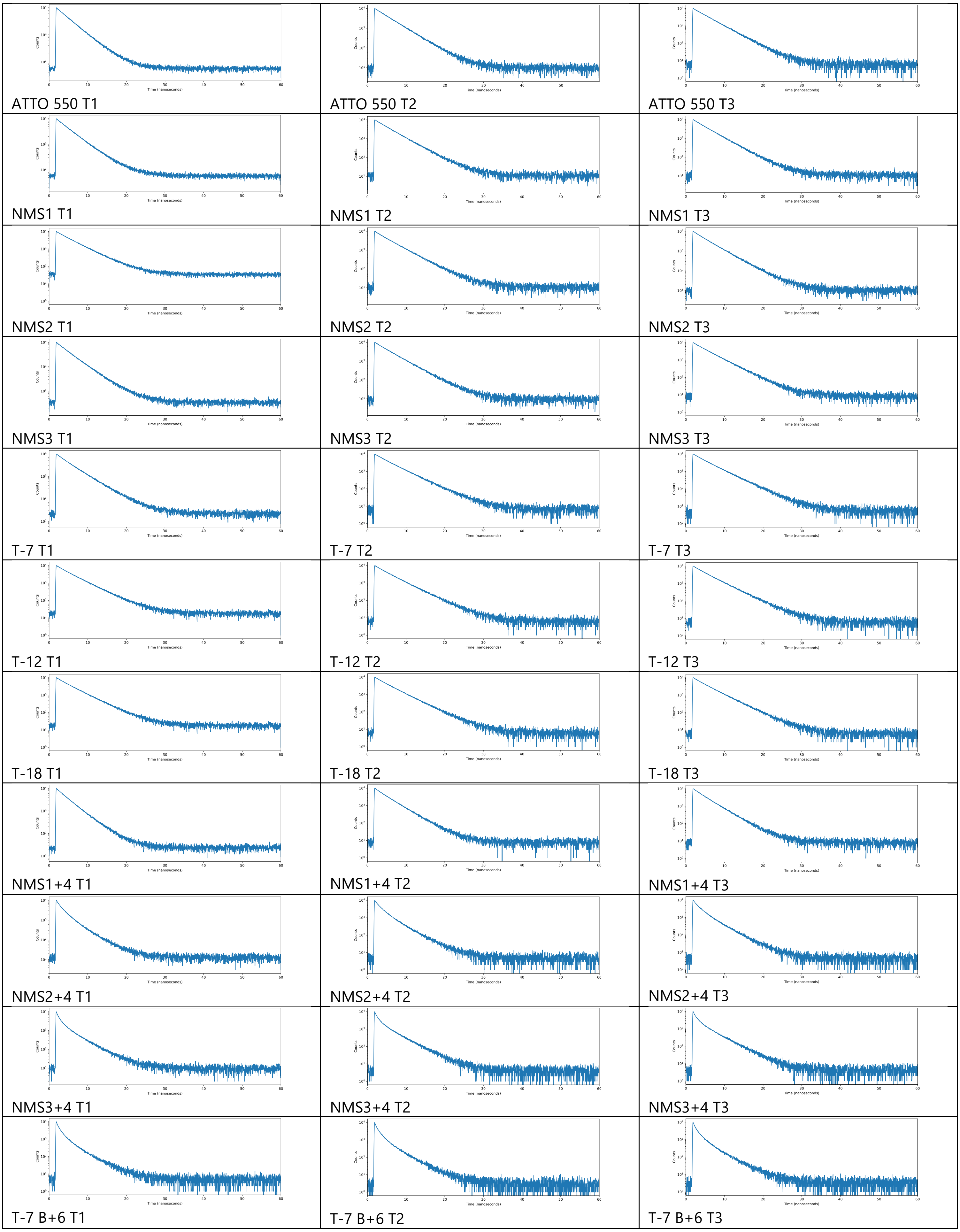
Counts
Time (nanoseconds)
ATTO 550 T1
ATTO 550 T2
ATTO 550 T3
NMS1 T1
NMS1 T2
NMS1 T3
NMS2 T1
NMS2 T2
NMS2 T3
NMS3 T1
NMS3 T2
NMS3 T3
T-7 T1
T-7 T2
T-7 T3
T-12 T1
T-12 T2
T-12 T3
T-18 T1
T-18 T2
T-18 T3
NMS1+4 T1
NMS1+4 T2
NMS1+4 T3
NMS2+4 T1
NMS2+4 T2
NMS2+4 T3
NMS3+4 T1
NMS3+4 T2
NMS3+4 T3
T-7 B+6 T1
T-7 B+6 T2
T-7 B+6 T3

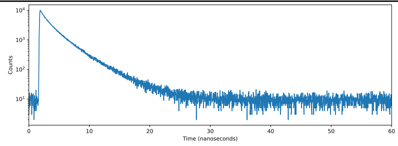

T-7 B+9 T1

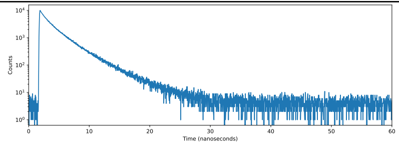

T-7 B+9 T2

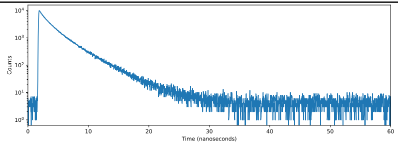

T-7 B+9 T3

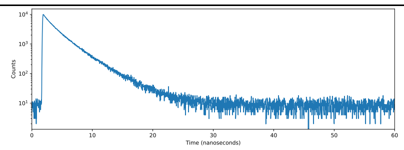

T-7 B+11 T1

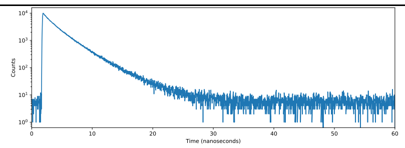

T-7 B+11 T2

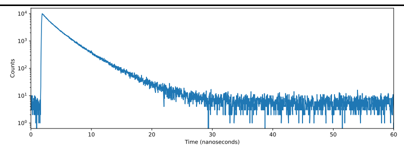

T-7 B+11 T3

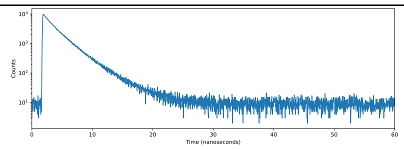

T-12 B+6 T1

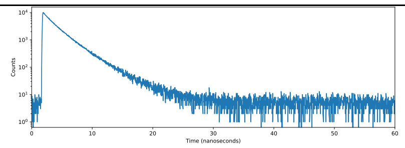

T-12 B+6 T2

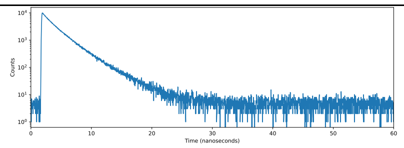

T-12 B+6 T3

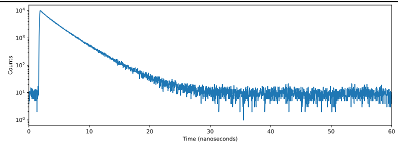

T-12 B+9 T1

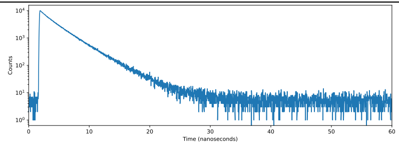

T-12 B+9 T2

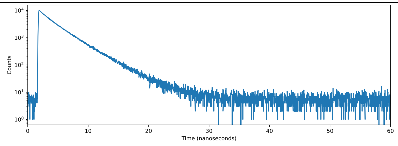

T-12 B+9 T3

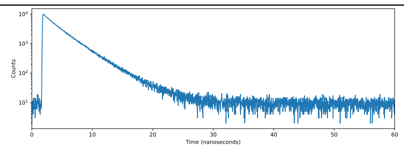

T-12 B+11 T1

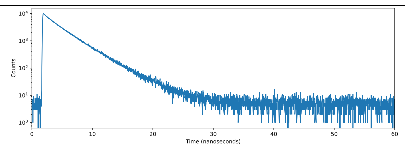

T-12 B+11 T2

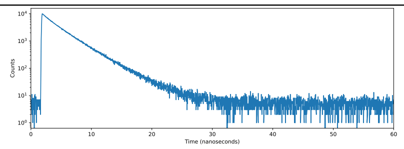

T-12 B+11 T3

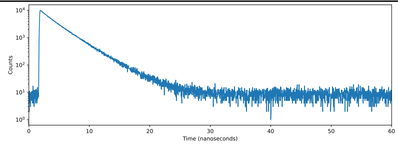

T-18 B+6 T1

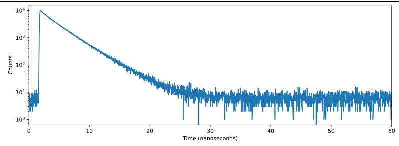

T-18 B+6 T2

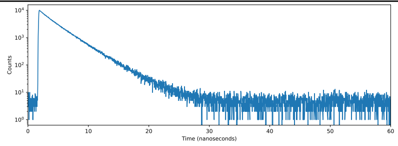

T-18 B+6 T3

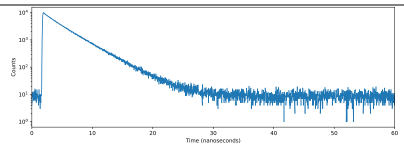

T-18 B+9 T1

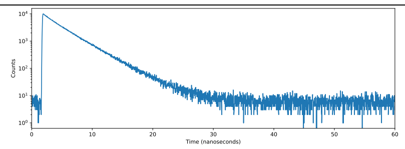

T-18 B+9 T2

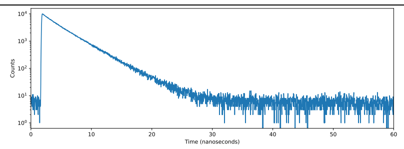

T-18 B+9 T3

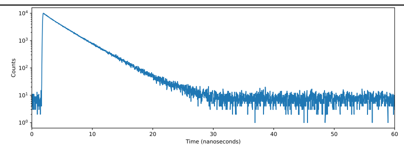

T-18 B+11 T1

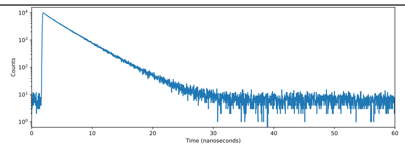

T-18 B+11 T2

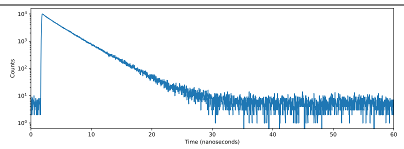

T-18 B+11 T3

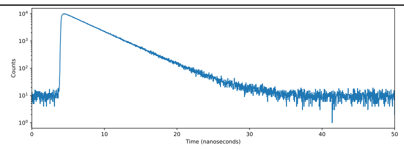

ATTO 647N T1

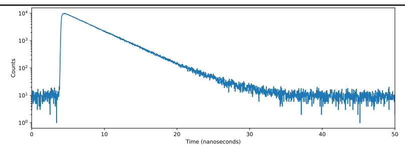

ATTO 647N T2

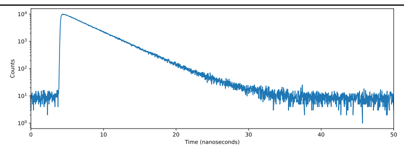

ATTO 647N T1

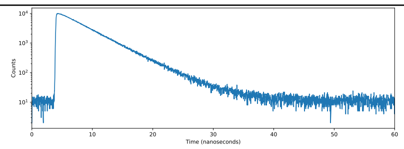

NMS4 T1

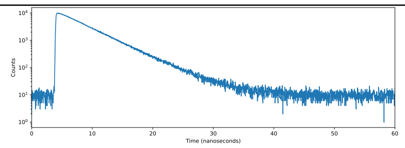

NMS4 T2

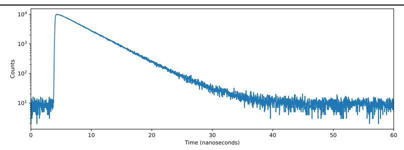

NMS4 T3

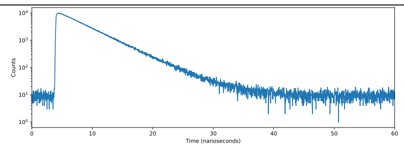

B+6 T1

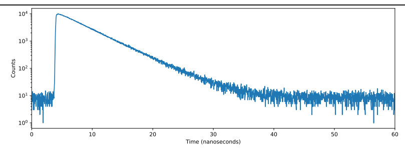

B+6 T2

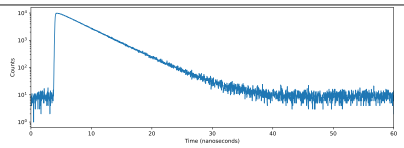

B+6 T3

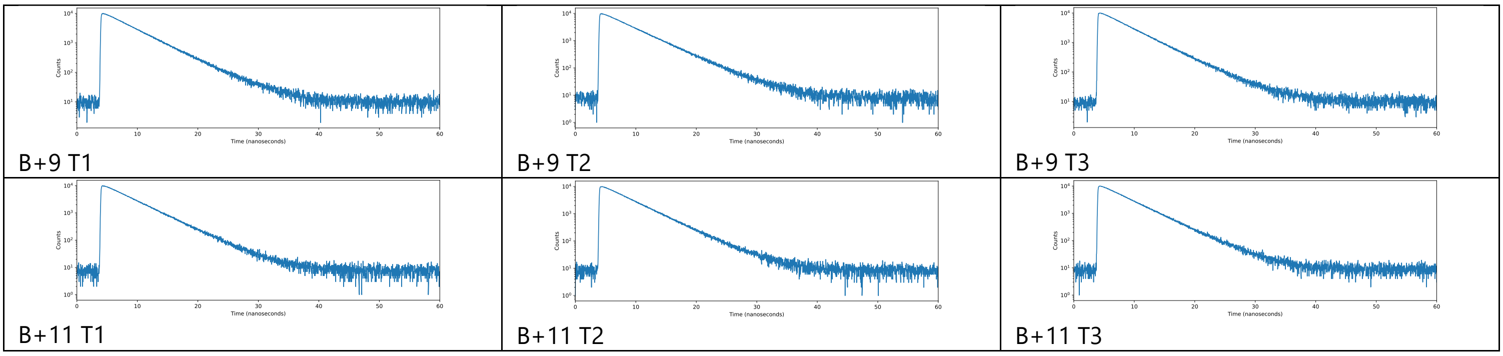

Supplementary Figure 34: Lifetime measured with 3 repeats for free dyes and each single labelled duplex DNA.

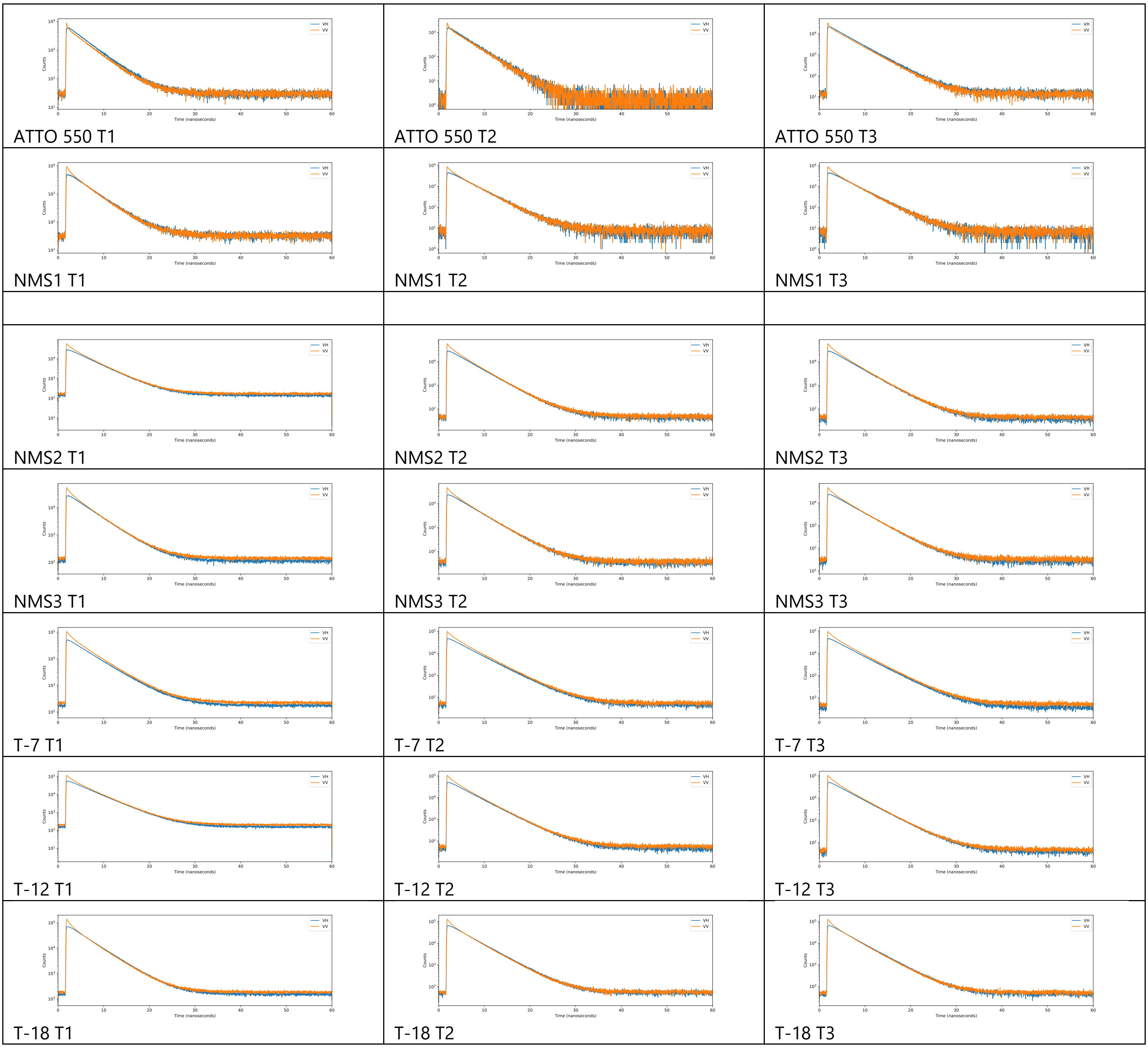

Supplementary Figure 35: VH and VV measurements for each of the free dyes and single labelled DNA duplex.

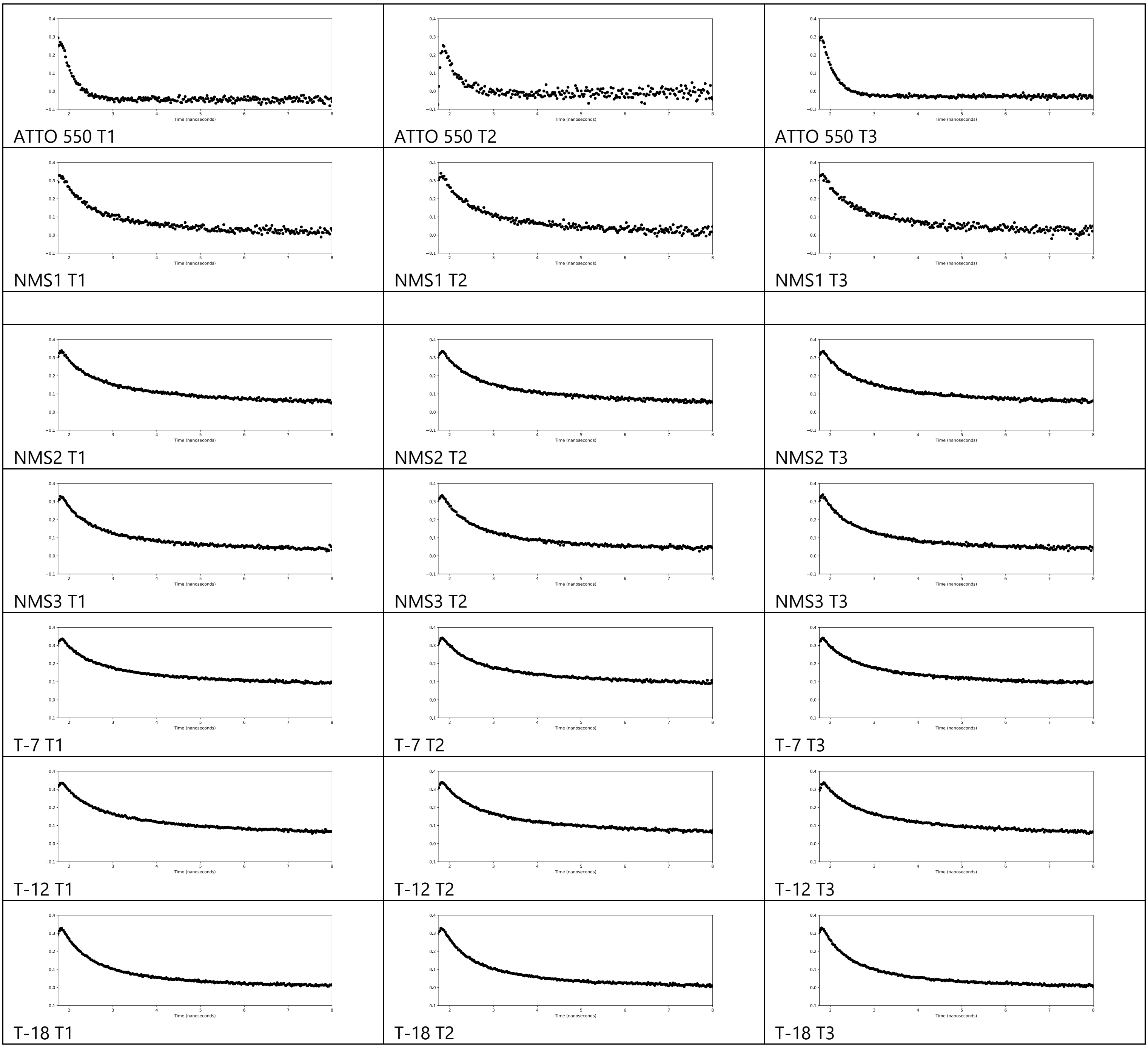

Supplementary Figure 36: Anisotropy decay for each of the free dyes and single labelled DNA duplex.

O6-MeG vs Control

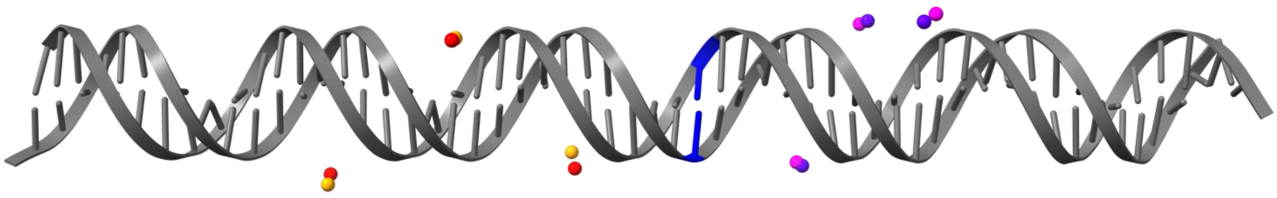

Ribo vs Control

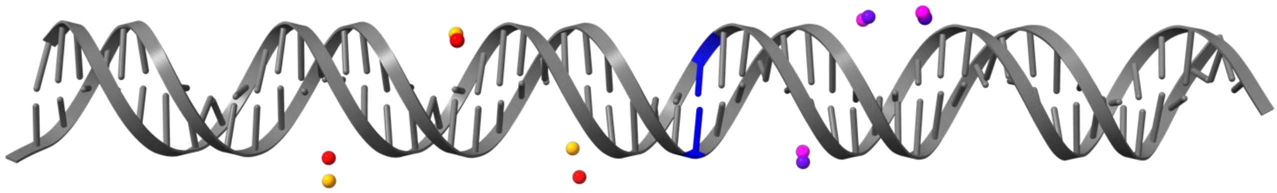

8-oxoG vs Control

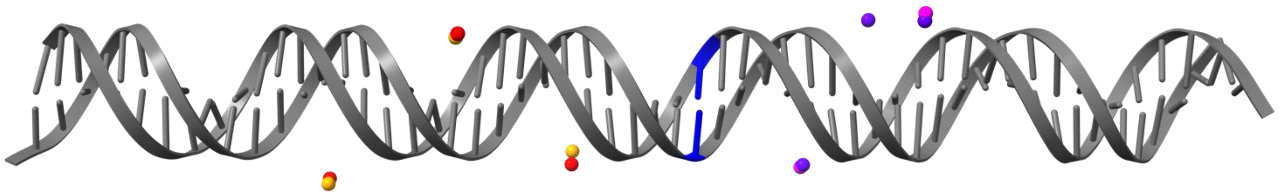

Abasic vs Control

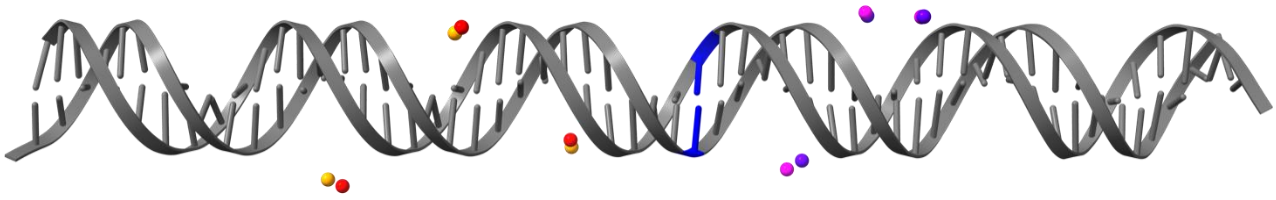

Nick vs Control

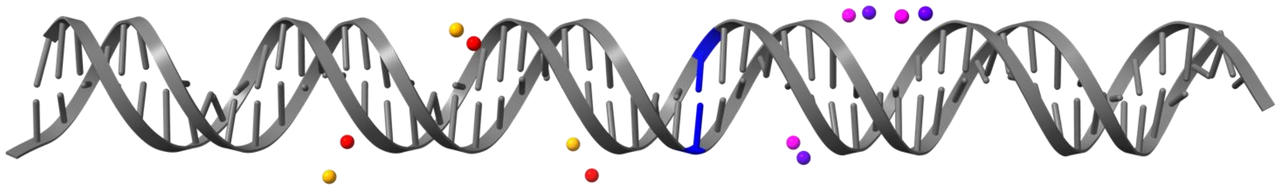

Gap vs Control

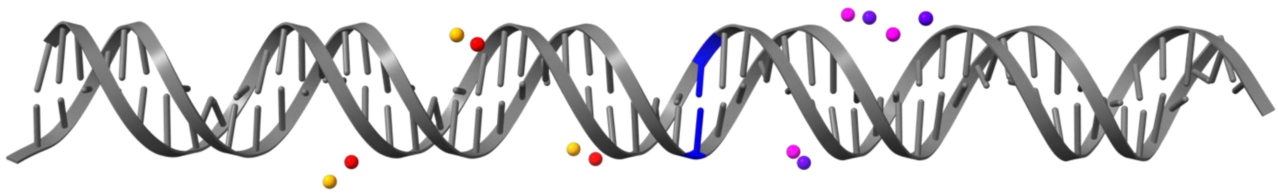

Supplementary Figure 37: Schematic showing the different type of damage position (blue) and how this has affected the mean position of the AV cloud with control position of donor ATTO 550 in pale orange and moved position in dark orange/red and for control position of acceptor ATTO 647N in dark purple and moved position in pink.

| Construct | Bend (°) Left | Bend (°) Right | Total Angle (°) Left | Total Angle (°) Right | Twist (°) Left | Twist (°) Right | RMSD Left (Å) | RMSD Right (Å) | RMSD Total (Å) |
|---|---|---|---|---|---|---|---|---|---|
| Ribo | 12.17 | 3.68 | 12.45 | 3.91 | –2.64 | –1.32 | 3.419 | 1.540 | 2.652 |
| 8-oxoG | 9.90 | 3.10 | 10.18 | 3.22 | –2.38 | –0.86 | 3.071 | 1.007 | 2.285 |
| Abasic | 2.04 | 1.11 | 5.04 | 3.14 | –4.61 | –2.94 | 2.431 | 1.608 | 2.061 |
| Nick | 10.88 | 2.61 | 12.74 | 3.22 | +6.62 | –1.88 | 4.730 | 2.972 | 3.950 |
| Gap | 5.49 | 5.74 | 5.88 | 5.77 | –2.12 | –0.51 | 3.813 | 3.791 | 3.802 |

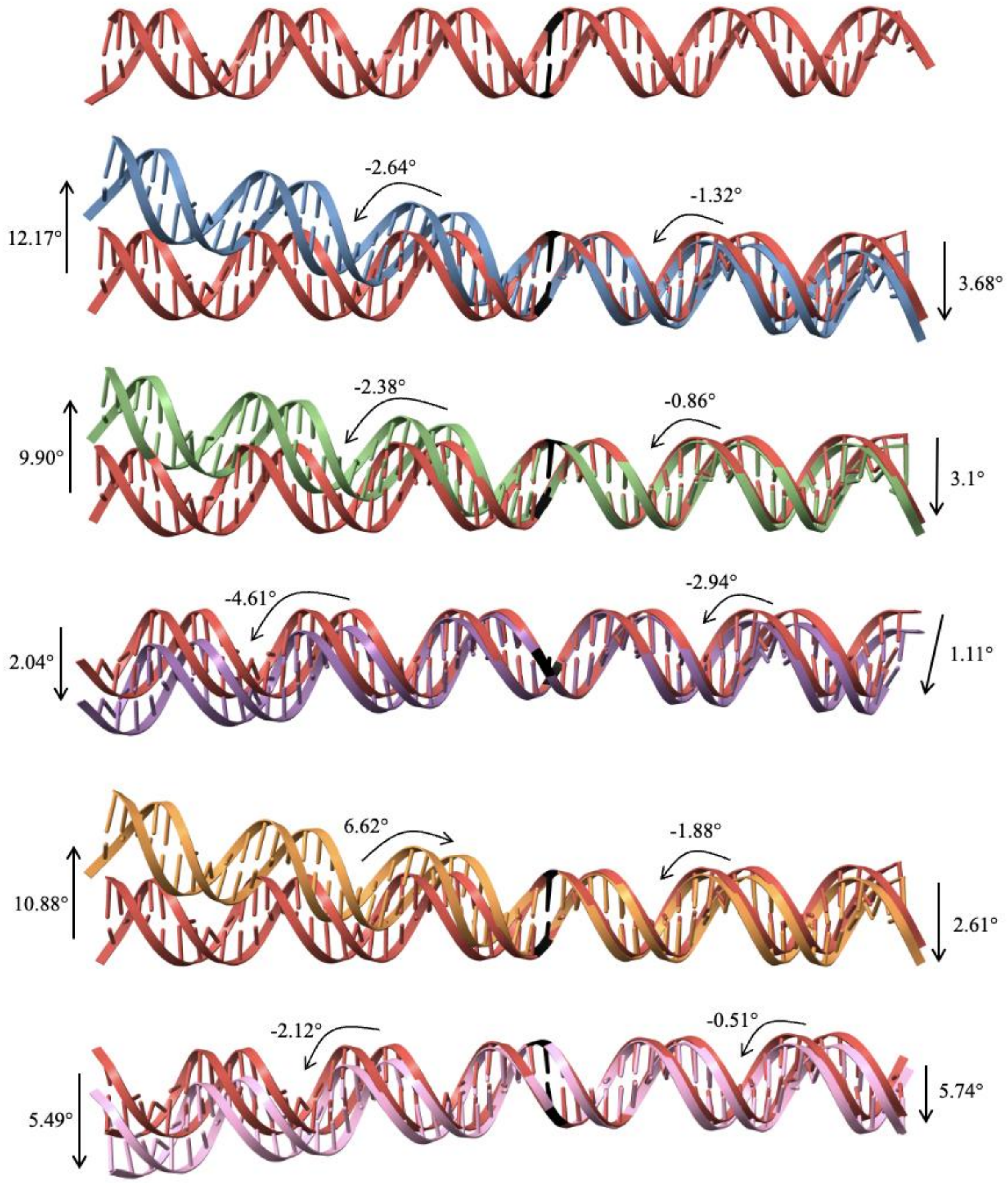

Supplementary Figure 38: Bending and twisting angles were determined from duplex models of chemically and structurally modified constructs relative to the control (unmodified) duplex shown in red. Positive twist values indicate over-twisting and negative values indicate under-twisting relative to the control. Left and right refer to the respective helical halves flanking the lesion site. RMSD values represent the deviation of each half from the control structure following alignment of the undamaged region.

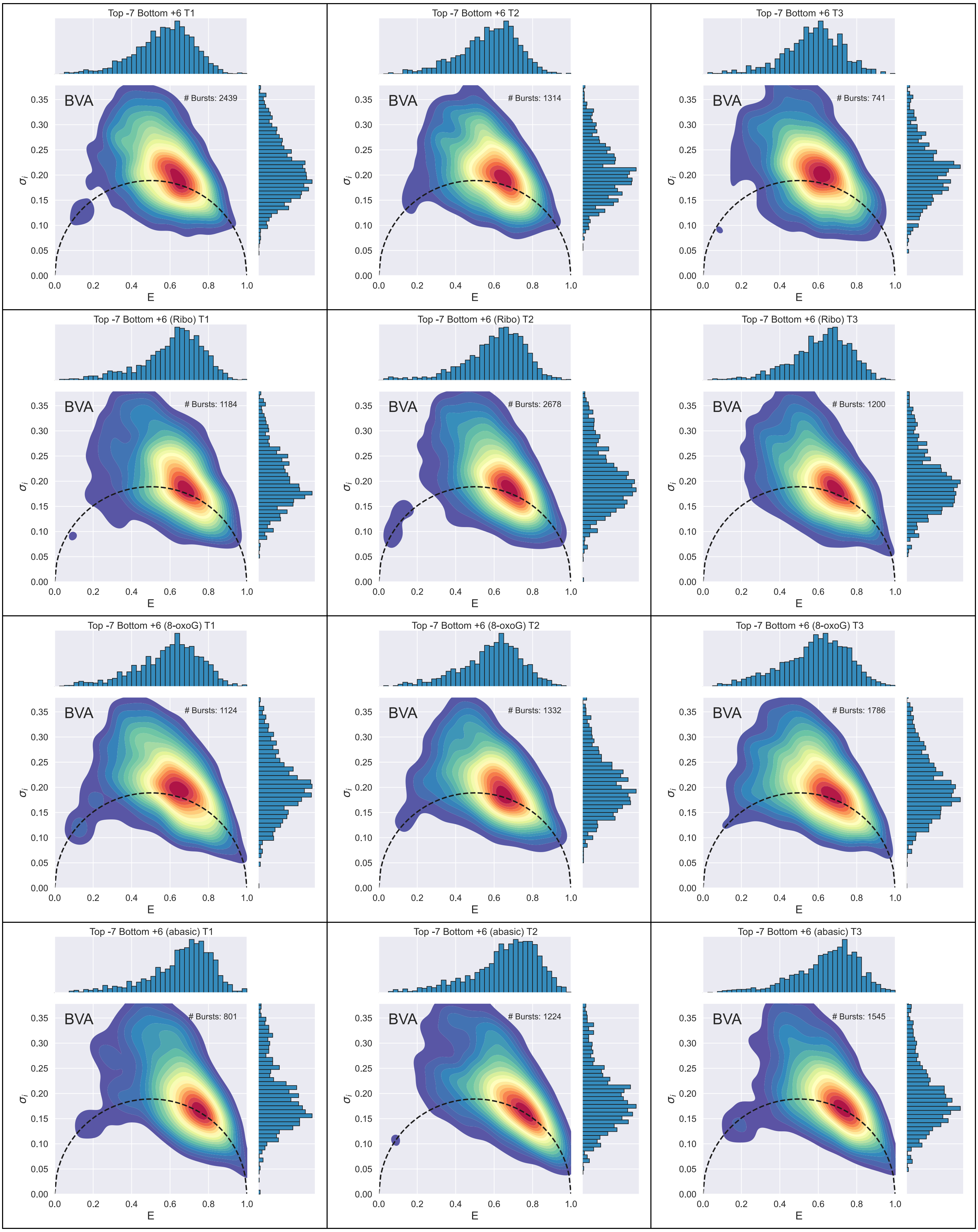
Top -7 Bottom +6 T1
BVA
### Bursts: 2439
Top -7 Bottom +6 T2
BVA
### Bursts: 1314
Top -7 Bottom +6 T3
BVA
### Bursts: 741
Top -7 Bottom +6 (Ribo) T1
BVA
### Bursts: 1184
Top -7 Bottom +6 (Ribo) T2
BVA
### Bursts: 2678
Top -7 Bottom +6 (Ribo) T3
BVA
### Bursts: 1200
Top -7 Bottom +6 (8-oxoG) T1
BVA
### Bursts: 1124
Top -7 Bottom +6 (8-oxoG) T2
BVA
### Bursts: 1332
Top -7 Bottom +6 (8-oxoG) T3
BVA
### Bursts: 1786
Top -7 Bottom +6 (abasic) T1
BVA
### Bursts: 801
Top -7 Bottom +6 (abasic) T2
BVA
### Bursts: 1224
Top -7 Bottom +6 (abasic) T3
BVA
### Bursts: 1545
E
σi
0.00 0.05 0.10 0.15 0.20 0.25 0.30 0.35
0.0 0.2 0.4 0.6 0.8 1.0

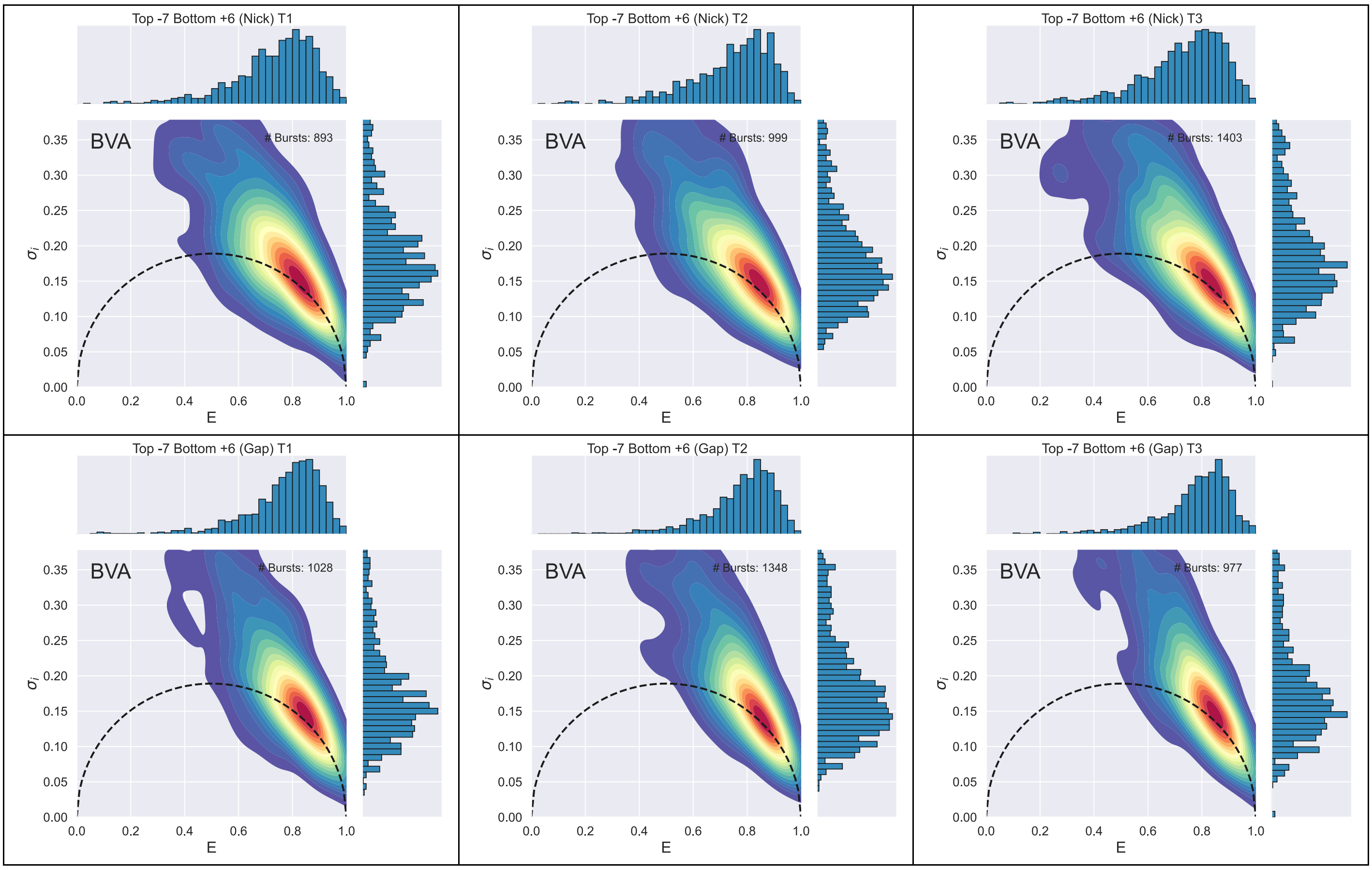

Supplementary Figure 39: BVA for each independent repeat of the seven duplex constructs for the dye-pair Top -7 Bottom +6.

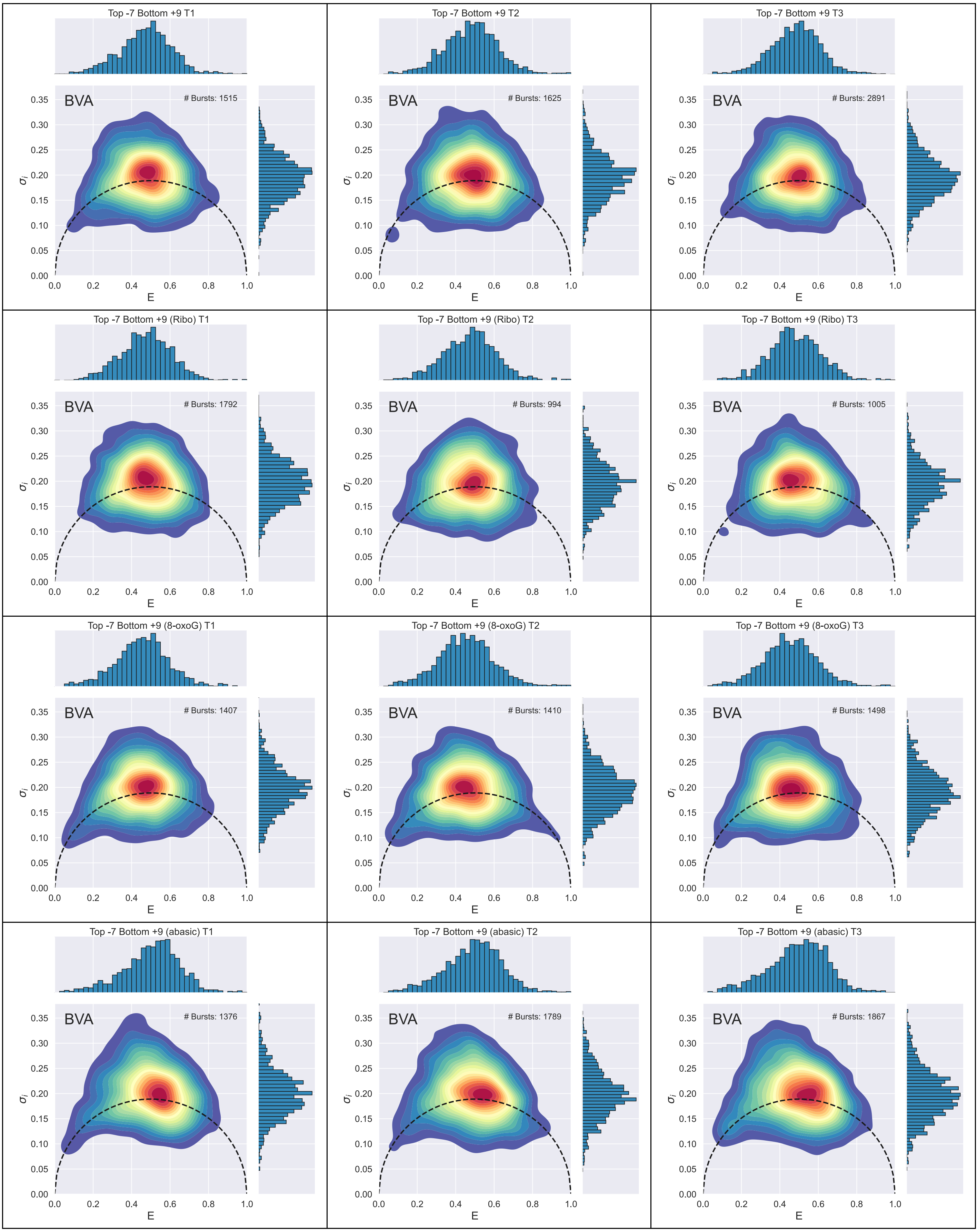
Top -7 Bottom +9 T1
BVA
### Bursts: 1515
Top -7 Bottom +9 T2
BVA
### Bursts: 1625
Top -7 Bottom +9 T3
BVA
### Bursts: 2891
Top -7 Bottom +9 (Ribo) T1
BVA
### Bursts: 1792
Top -7 Bottom +9 (Ribo) T2
BVA
### Bursts: 994
Top -7 Bottom +9 (Ribo) T3
BVA
### Bursts: 1005
Top -7 Bottom +9 (8-oxoG) T1
BVA
### Bursts: 1407
Top -7 Bottom +9 (8-oxoG) T2
BVA
### Bursts: 1410
Top -7 Bottom +9 (8-oxoG) T3
BVA
### Bursts: 1498
Top -7 Bottom +9 (abasic) T1
BVA
### Bursts: 1376
Top -7 Bottom +9 (abasic) T2
BVA
### Bursts: 1789
Top -7 Bottom +9 (abasic) T3
BVA
### Bursts: 1867
σi
E

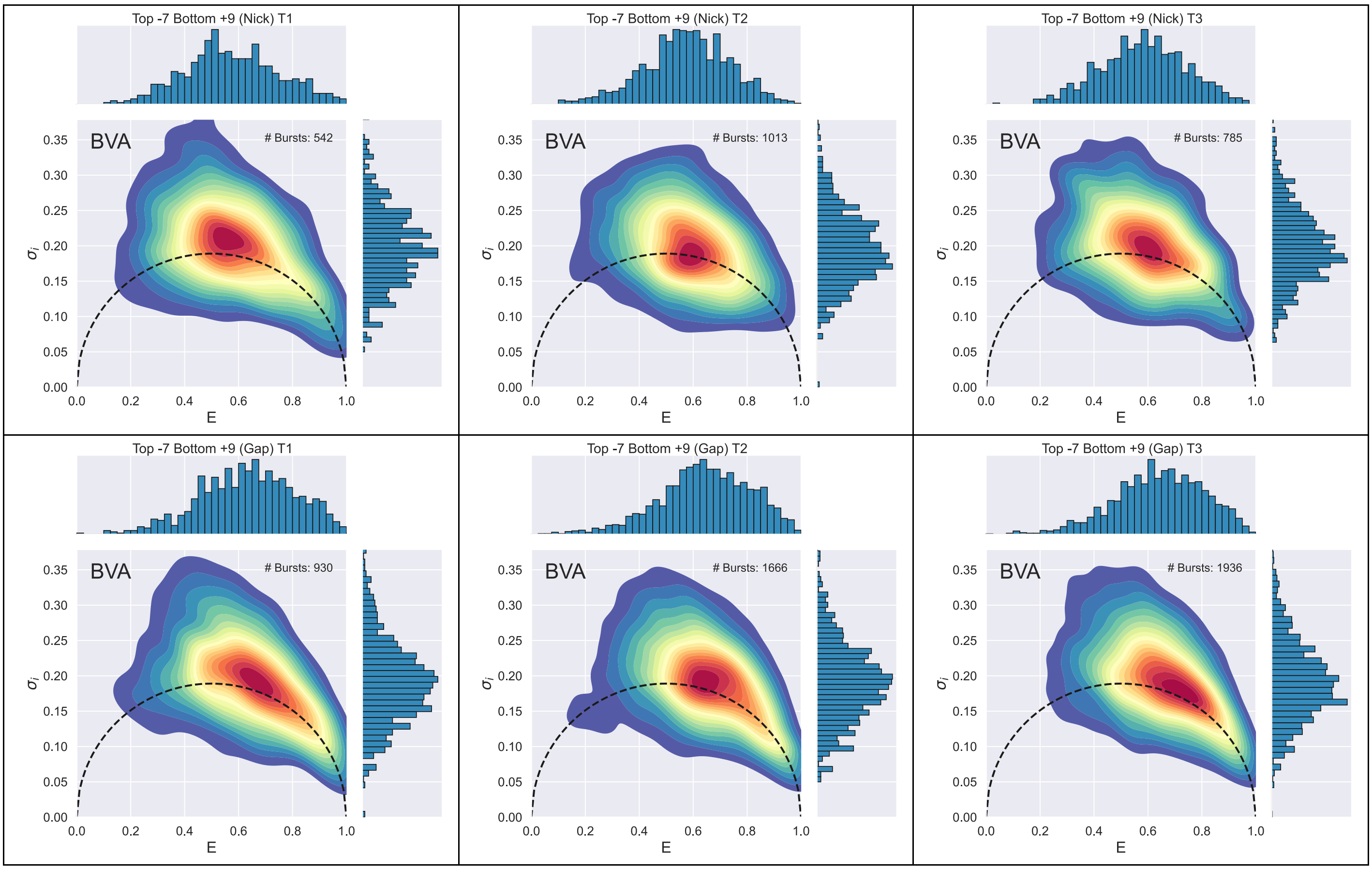

Supplementary Figure 40: BVA for each independent repeat of the seven duplex constructs for the dye-pair Top -7 Bottom +9.

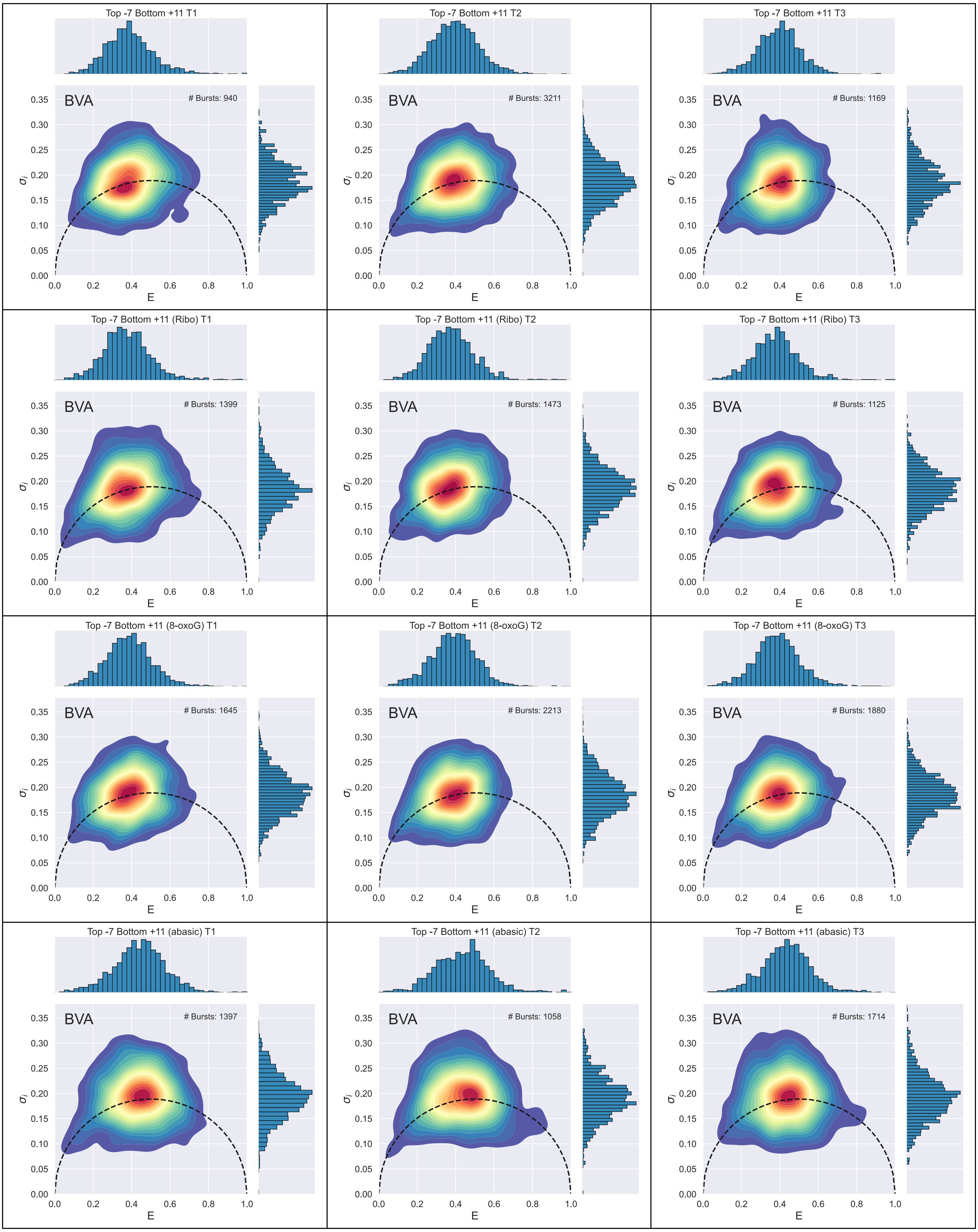
Top -7 Bottom +11 T1
BVA
### Bursts: 940
Top -7 Bottom +11 T2
BVA
### Bursts: 3211
Top -7 Bottom +11 T3
BVA
### Bursts: 1169
Top -7 Bottom +11 (Ribo) T1
BVA
### Bursts: 1399
Top -7 Bottom +11 (Ribo) T2
BVA
### Bursts: 1473
Top -7 Bottom +11 (Ribo) T3
BVA
### Bursts: 1125
Top -7 Bottom +11 (8-oxoG) T1
BVA
### Bursts: 1645
Top -7 Bottom +11 (8-oxoG) T2
BVA
### Bursts: 2213
Top -7 Bottom +11 (8-oxoG) T3
BVA
### Bursts: 1880
Top -7 Bottom +11 (abasic) T1
BVA
### Bursts: 1397
Top -7 Bottom +11 (abasic) T2
BVA
### Bursts: 1058
Top -7 Bottom +11 (abasic) T3
BVA
### Bursts: 1714
$\sigma_i$
E
0.00 0.05 0.10 0.15 0.20 0.25 0.30 0.35
0.0 0.2 0.4 0.6 0.8 1.0

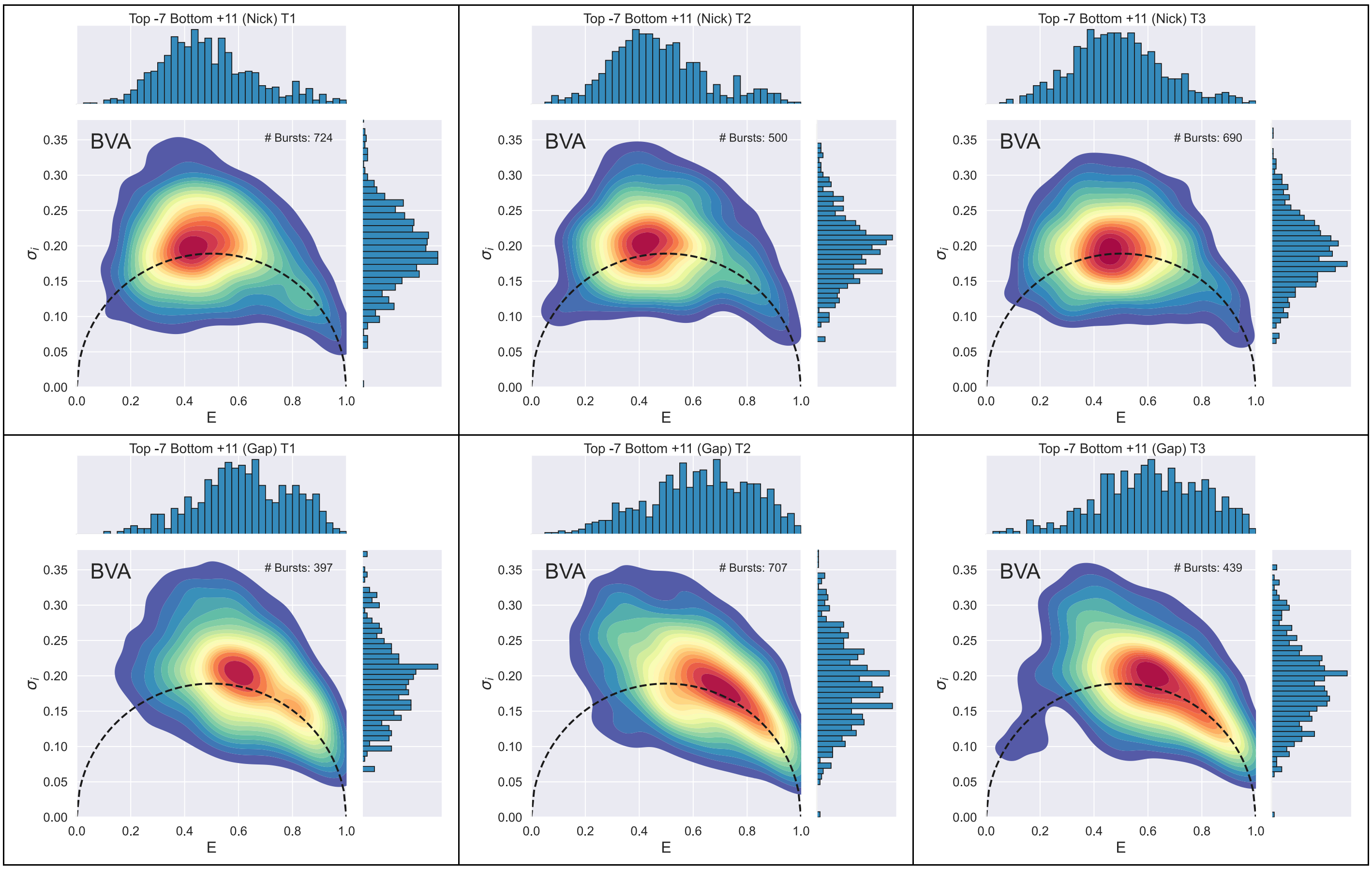

Supplementary Figure 41: BVA for each independent repeat of the seven duplex constructs for the dye-pair Top -7 Bottom +11.

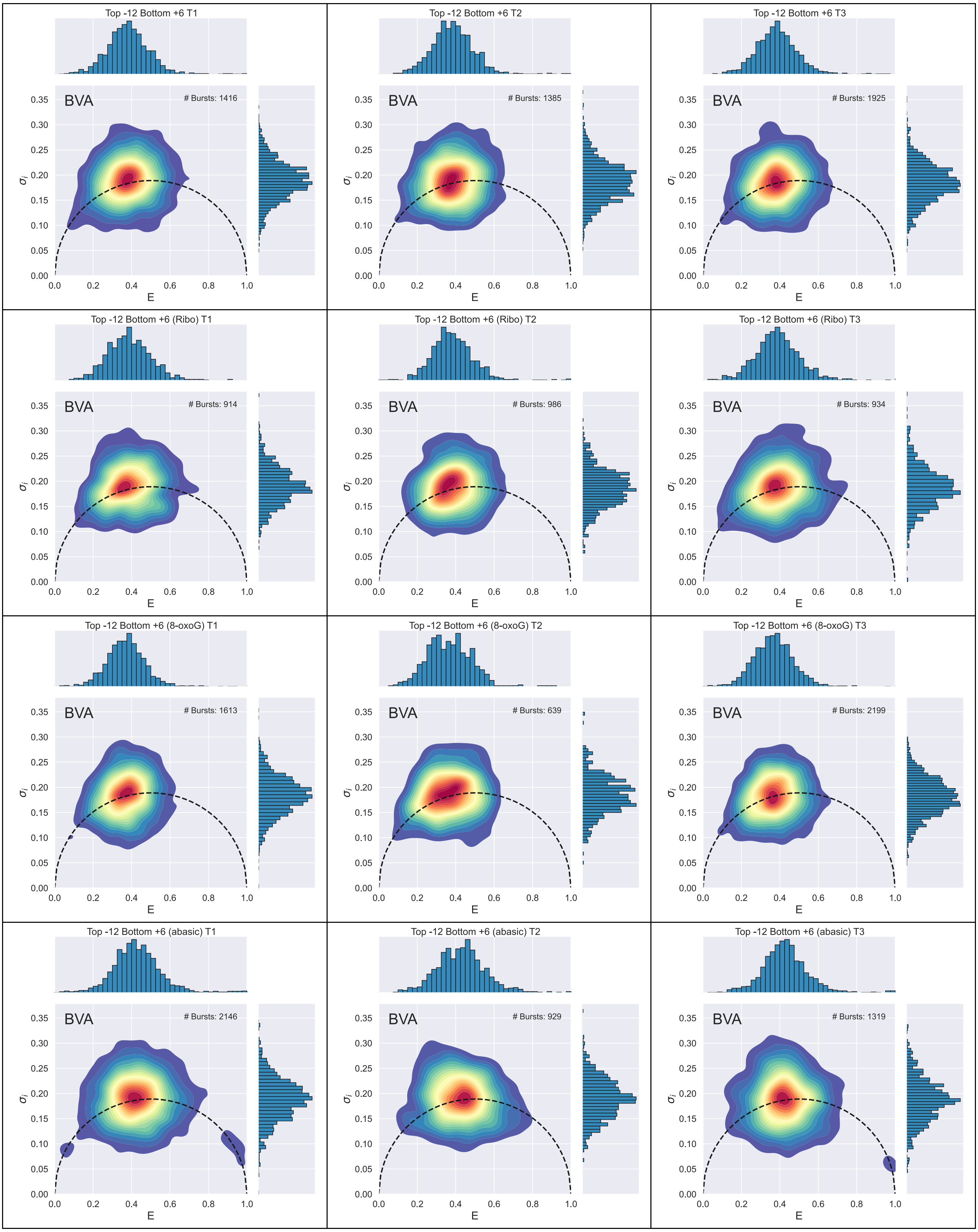
Top -12 Bottom +6 T1
BVA
### Bursts: 1416
Top -12 Bottom +6 T2
BVA
### Bursts: 1385
Top -12 Bottom +6 T3
BVA
### Bursts: 1925
Top -12 Bottom +6 (Ribo) T1
BVA
### Bursts: 914
Top -12 Bottom +6 (Ribo) T2
BVA
### Bursts: 986
Top -12 Bottom +6 (Ribo) T3
BVA
### Bursts: 934
Top -12 Bottom +6 (8-oxoG) T1
BVA
### Bursts: 1613
Top -12 Bottom +6 (8-oxoG) T2
BVA
### Bursts: 639
Top -12 Bottom +6 (8-oxoG) T3
BVA
### Bursts: 2199
Top -12 Bottom +6 (abasic) T1
BVA
### Bursts: 2146
Top -12 Bottom +6 (abasic) T2
BVA
### Bursts: 929
Top -12 Bottom +6 (abasic) T3
BVA
### Bursts: 1319
$\sigma_i$
E
0.00
0.05
0.10
0.15
0.20
0.25
0.30
0.35
0.0
0.2
0.4
0.6
0.8
1.0

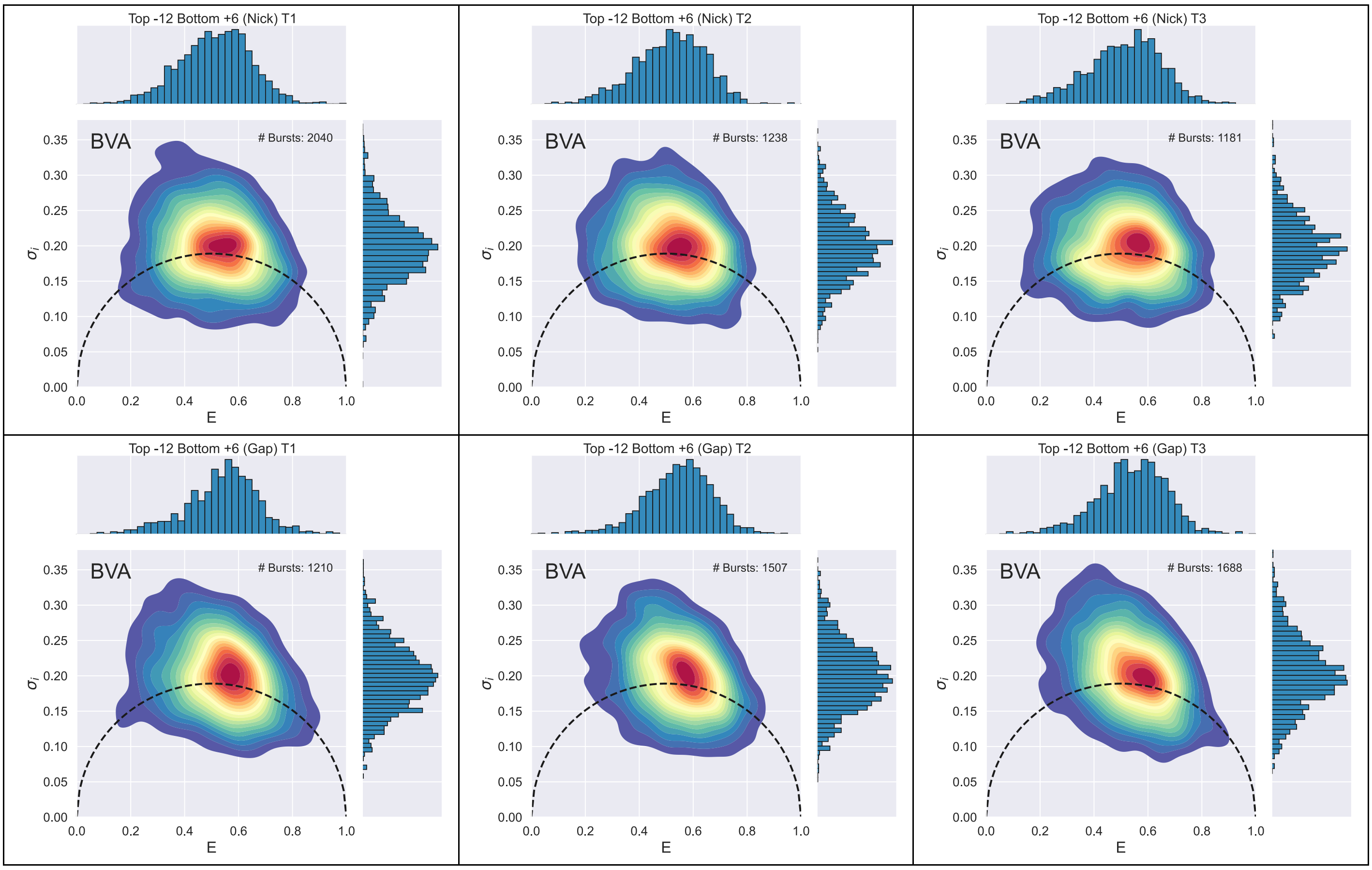

Supplementary Figure 42: BVA for each independent repeat of the seven duplex constructs for the dye-pair Top -12 Bottom +6.

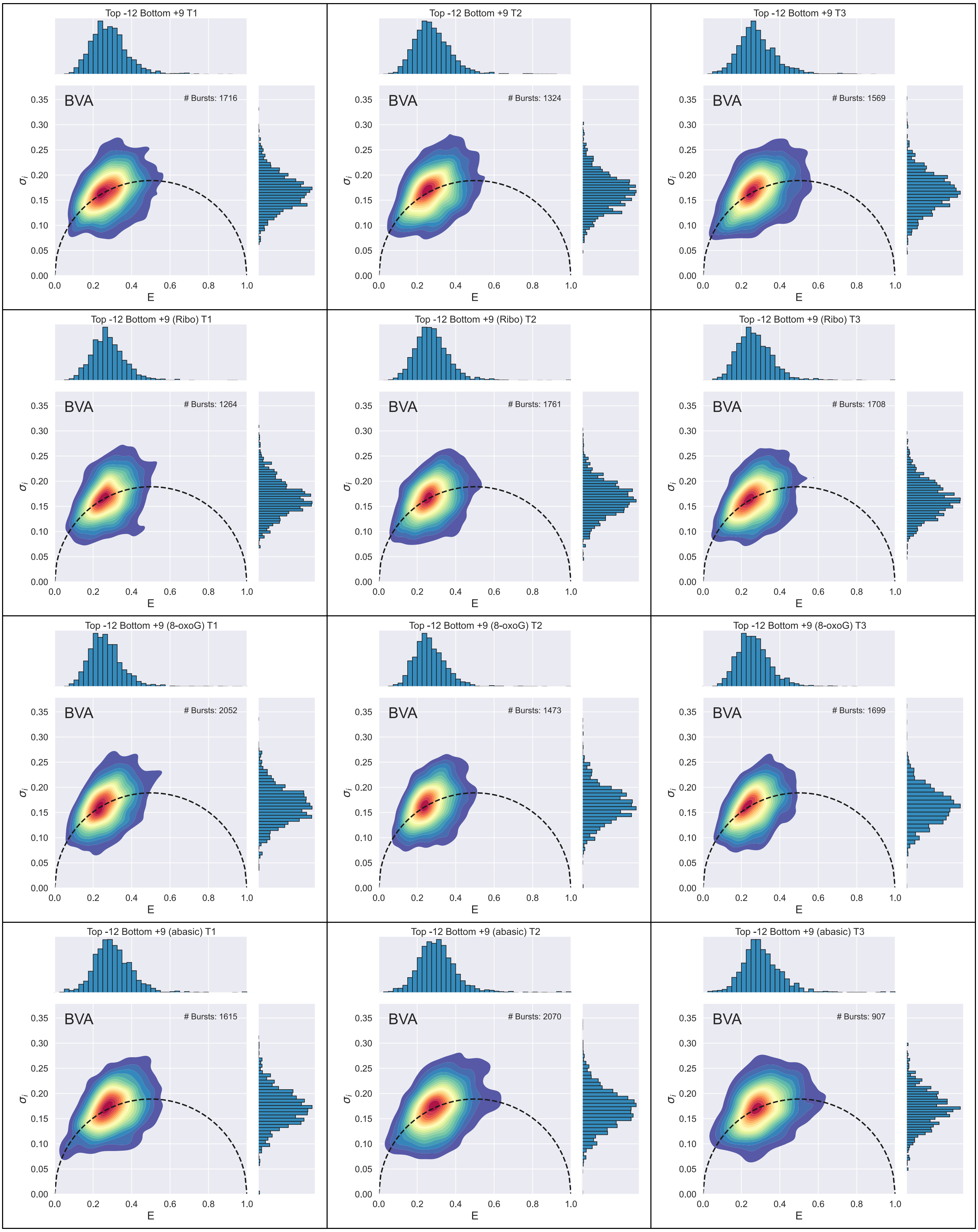

Top -12 Bottom +9 T1
BVA
### Bursts: 1716
Top -12 Bottom +9 T2
BVA
### Bursts: 1324
Top -12 Bottom +9 T3
BVA
### Bursts: 1569
Top -12 Bottom +9 (Ribo) T1
BVA
### Bursts: 1264
Top -12 Bottom +9 (Ribo) T2
BVA
### Bursts: 1761
Top -12 Bottom +9 (Ribo) T3
BVA
### Bursts: 1708
Top -12 Bottom +9 (8-oxoG) T1
BVA
### Bursts: 2052
Top -12 Bottom +9 (8-oxoG) T2
BVA
### Bursts: 1473
Top -12 Bottom +9 (8-oxoG) T3
BVA
### Bursts: 1699
Top -12 Bottom +9 (abasic) T1
BVA
### Bursts: 1615
Top -12 Bottom +9 (abasic) T2
BVA
### Bursts: 2070
Top -12 Bottom +9 (abasic) T3
BVA
### Bursts: 907
$\sigma_i$
0.00 0.05 0.10 0.15 0.20 0.25 0.30 0.35
E
0.0 0.2 0.4 0.6 0.8 1.0

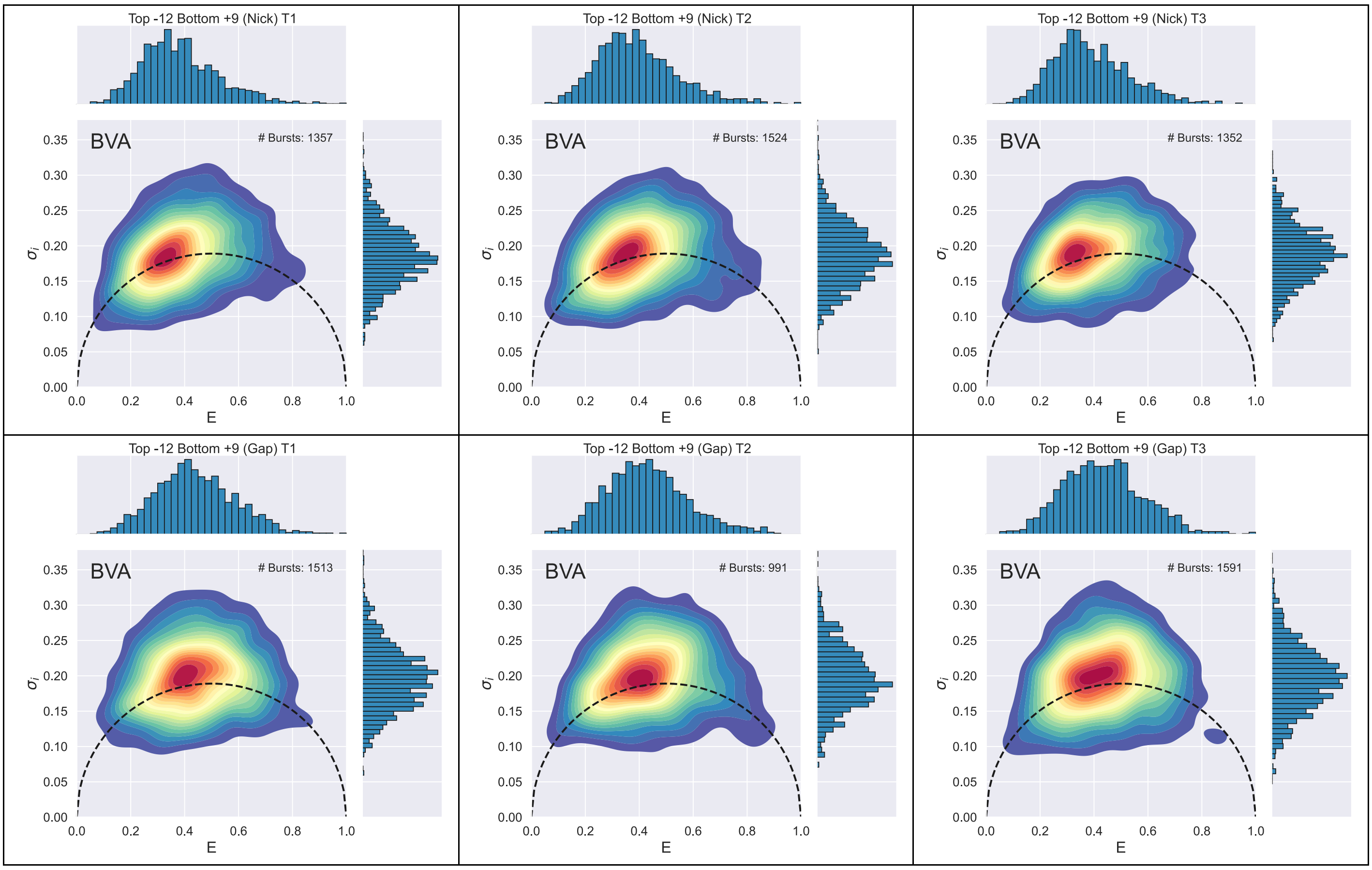

Supplementary Figure 43: BVA for each independent repeat of the seven duplex constructs for the dye-pair Top -12 Bottom +9.

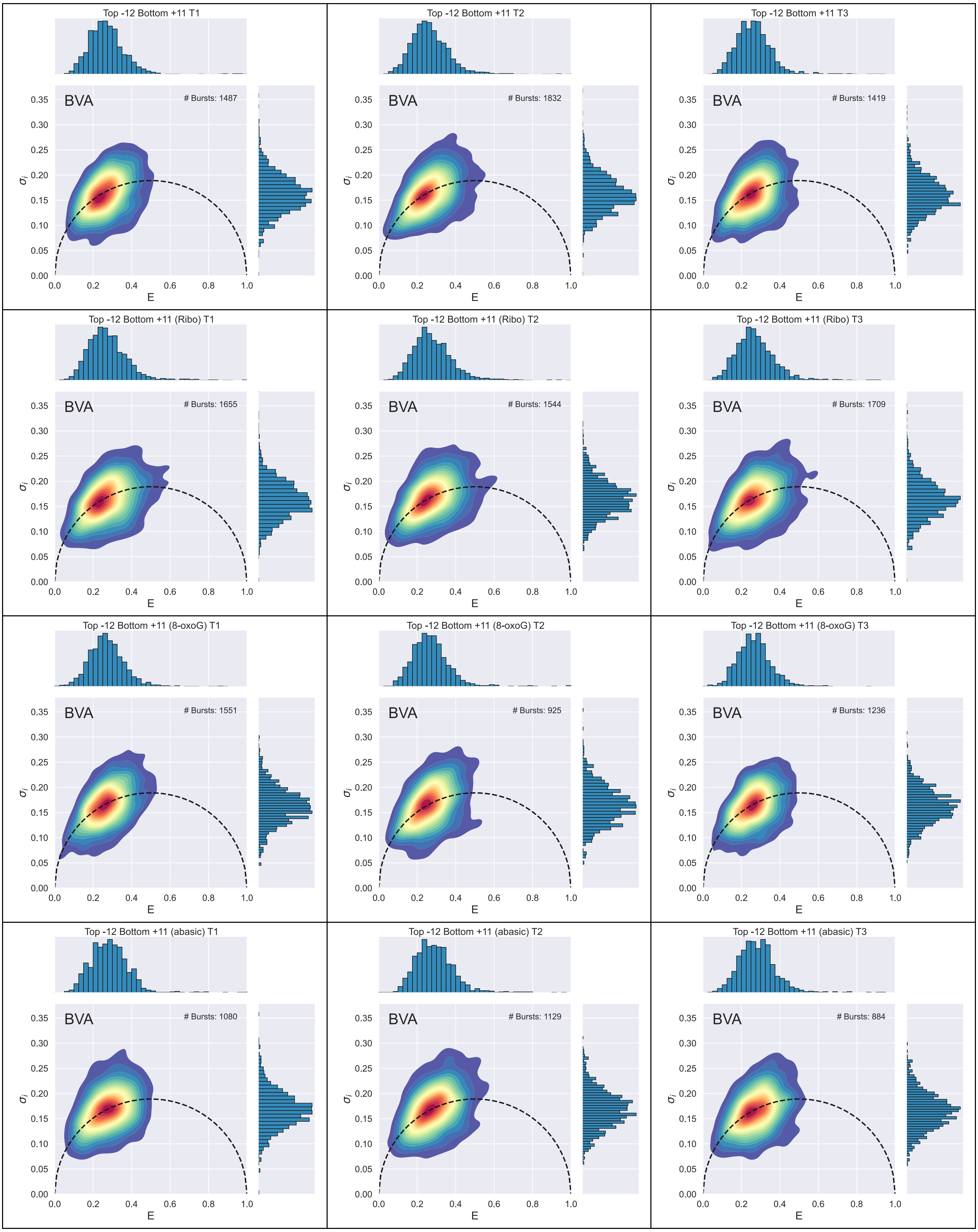
Top -12 Bottom +11 T1
BVA
### Bursts: 1487
Top -12 Bottom +11 T2
BVA
### Bursts: 1832
Top -12 Bottom +11 T3
BVA
### Bursts: 1419
Top -12 Bottom +11 (Ribo) T1
BVA
### Bursts: 1655
Top -12 Bottom +11 (Ribo) T2
BVA
### Bursts: 1544
Top -12 Bottom +11 (Ribo) T3
BVA
### Bursts: 1709
Top -12 Bottom +11 (8-oxoG) T1
BVA
### Bursts: 1551
Top -12 Bottom +11 (8-oxoG) T2
BVA
### Bursts: 925
Top -12 Bottom +11 (8-oxoG) T3
BVA
### Bursts: 1236
Top -12 Bottom +11 (abasic) T1
BVA
### Bursts: 1080
Top -12 Bottom +11 (abasic) T2
BVA
### Bursts: 1129
Top -12 Bottom +11 (abasic) T3
BVA
### Bursts: 884
E
$\sigma_i$

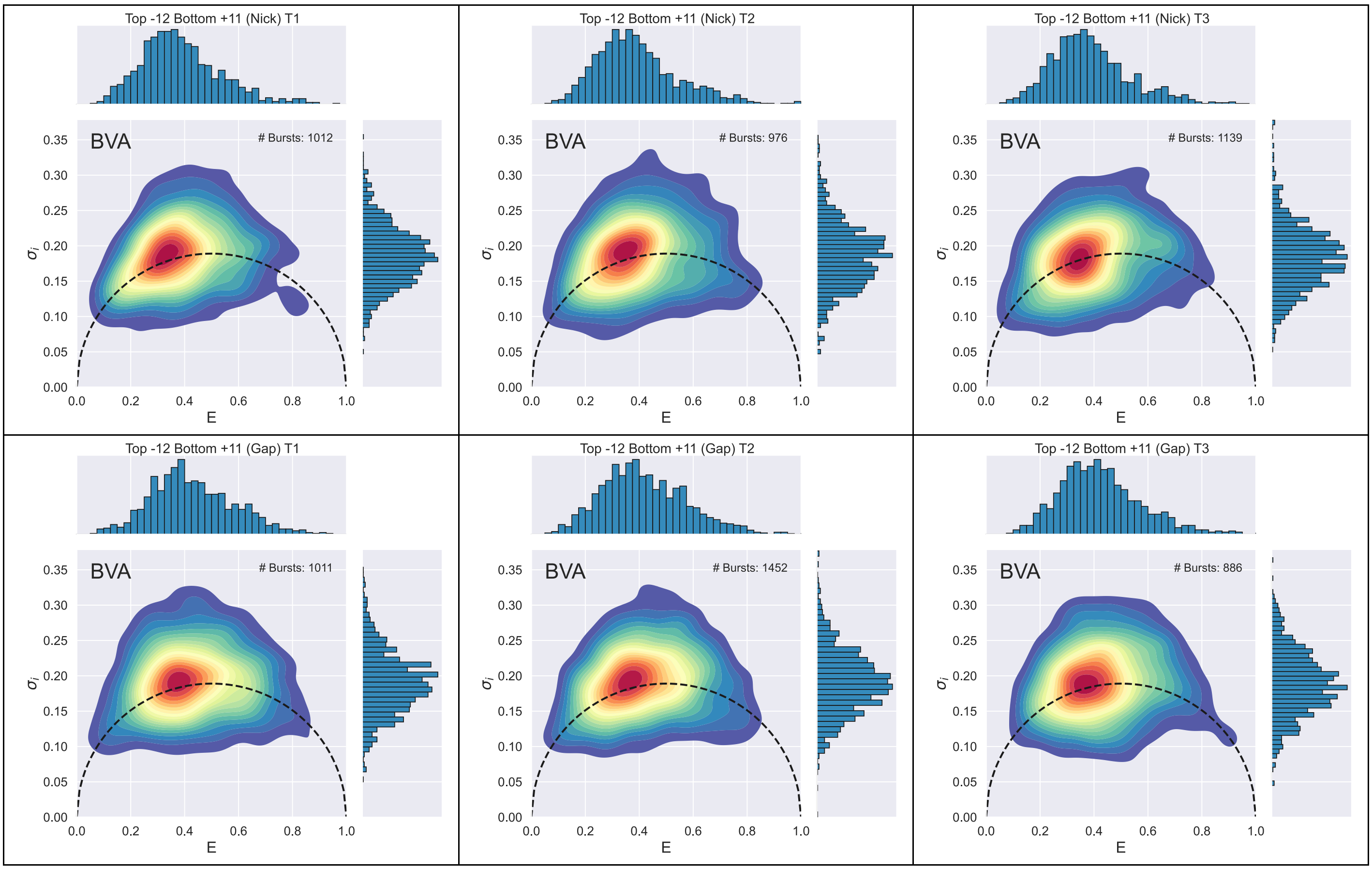

Supplementary Figure 44: BVA for each independent repeat of the seven duplex constructs for the dye-pair Top -12 Bottom +11.

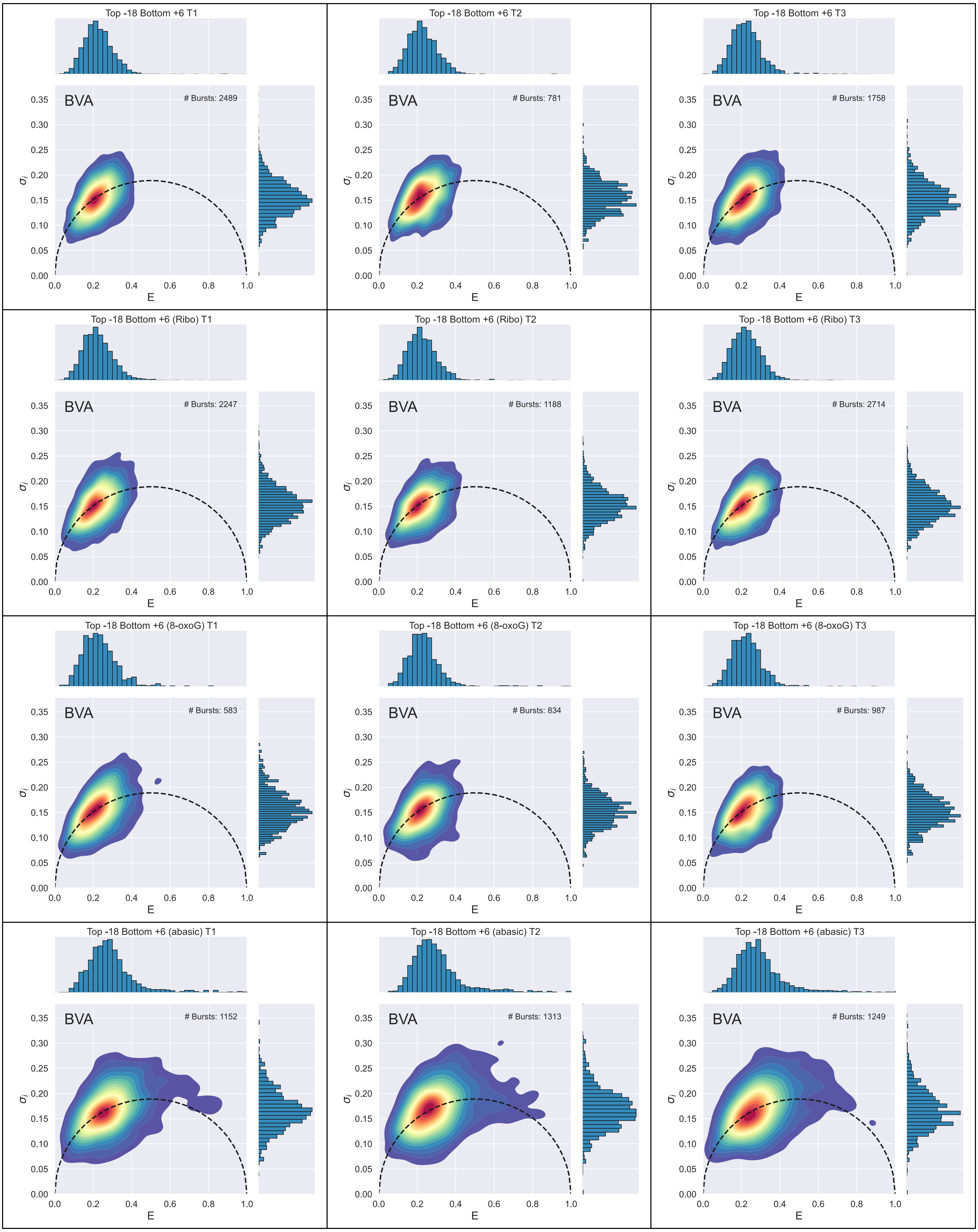
Top -18 Bottom +6 T1
BVA
### Bursts: 2489
Top -18 Bottom +6 T2
BVA
### Bursts: 781
Top -18 Bottom +6 T3
BVA
### Bursts: 1758
Top -18 Bottom +6 (Ribo) T1
BVA
### Bursts: 2247
Top -18 Bottom +6 (Ribo) T2
BVA
### Bursts: 1188
Top -18 Bottom +6 (Ribo) T3
BVA
### Bursts: 2714
Top -18 Bottom +6 (8-oxoG) T1
BVA
### Bursts: 583
Top -18 Bottom +6 (8-oxoG) T2
BVA
### Bursts: 834
Top -18 Bottom +6 (8-oxoG) T3
BVA
### Bursts: 987
Top -18 Bottom +6 (abasic) T1
BVA
### Bursts: 1152
Top -18 Bottom +6 (abasic) T2
BVA
### Bursts: 1313
Top -18 Bottom +6 (abasic) T3
BVA
### Bursts: 1249
$\sigma_i$
E
0.00
0.05
0.10
0.15
0.20
0.25
0.30
0.35
0.0
0.2
0.4
0.6
0.8
1.0

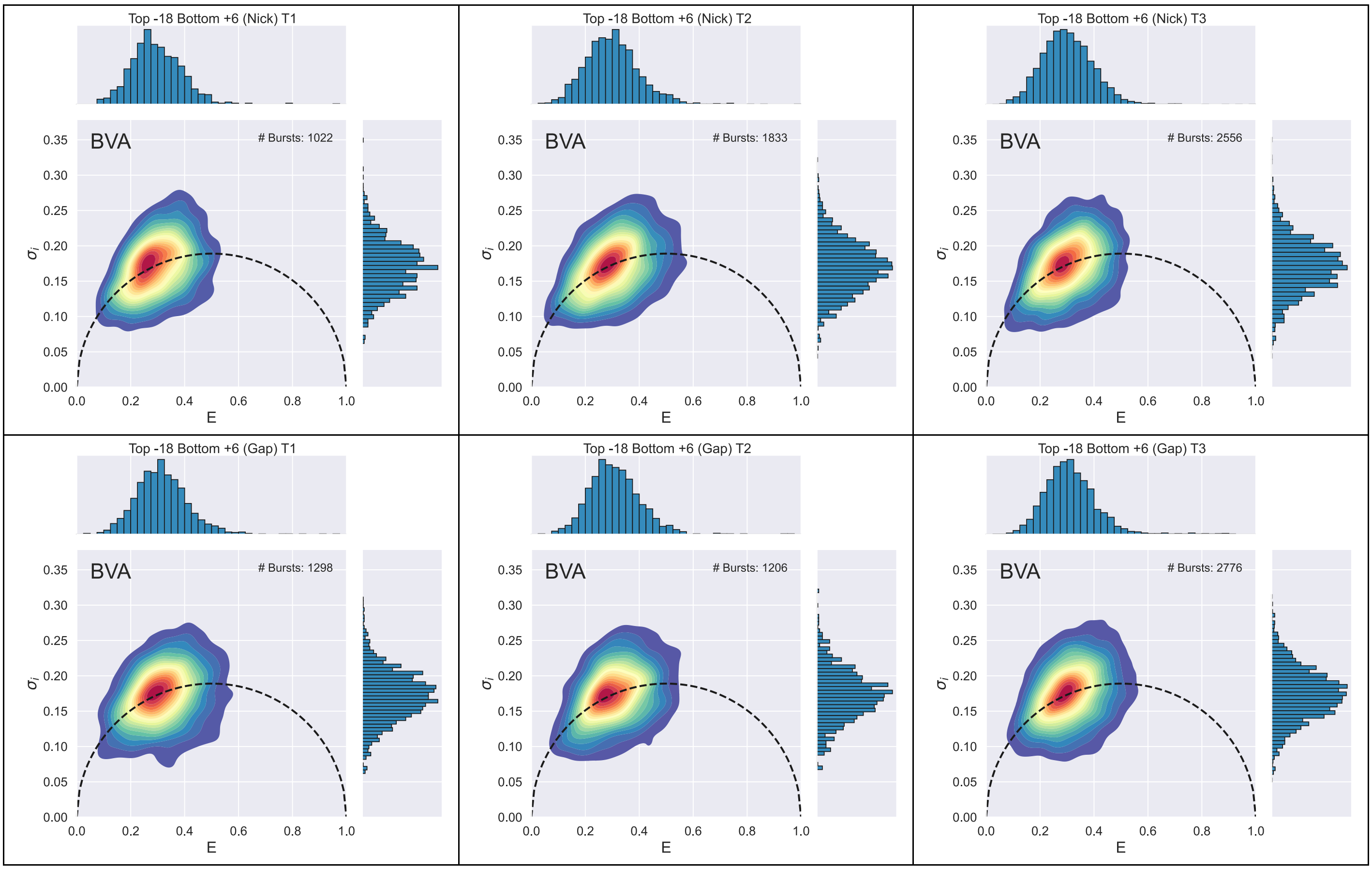

Supplementary Figure 45: BVA for each independent repeat of the seven duplex constructs for the dye-pair Top -18 Bottom +6.

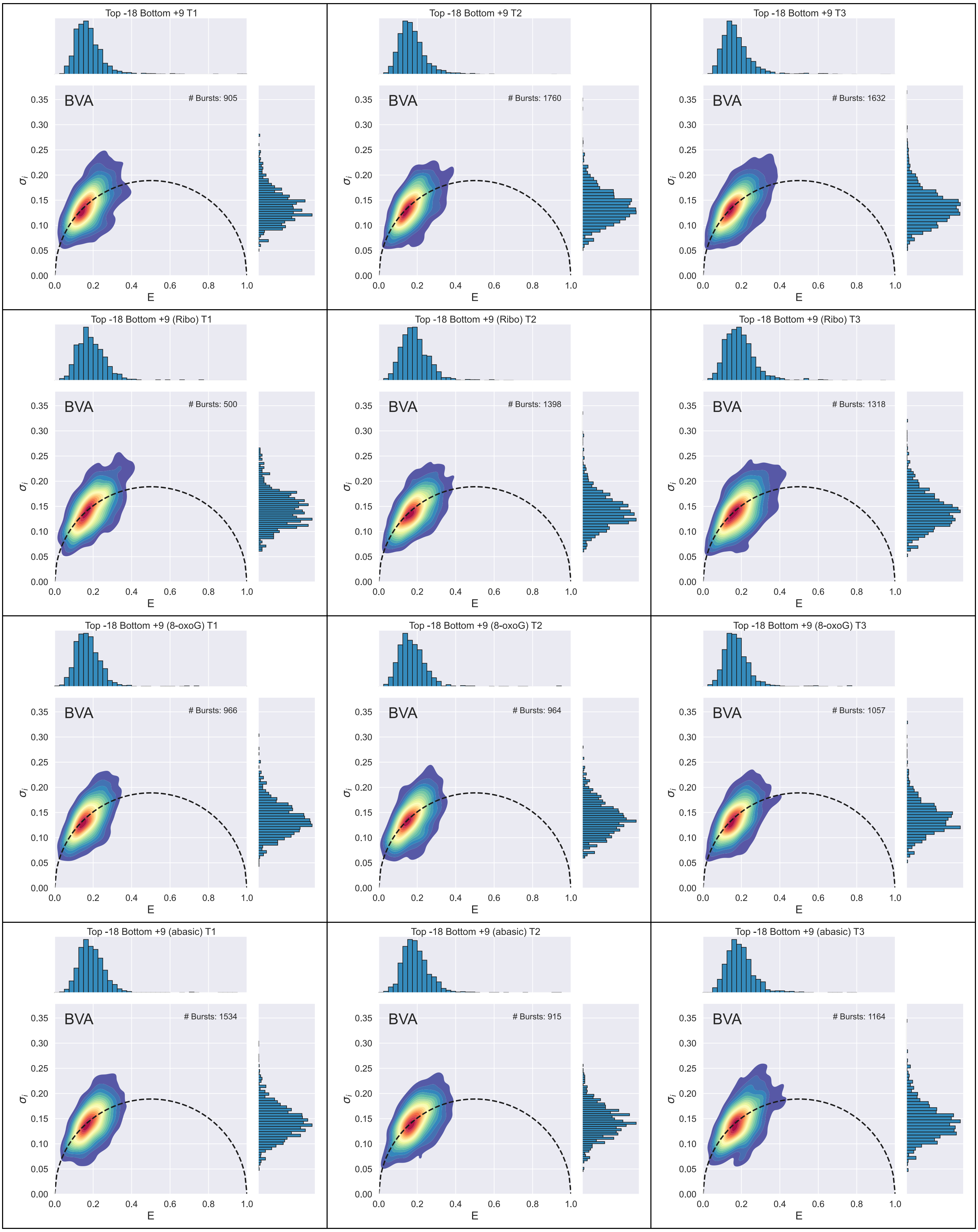
Top -18 Bottom +9 T1
BVA
### Bursts: 905
Top -18 Bottom +9 T2
BVA
### Bursts: 1760
Top -18 Bottom +9 T3
BVA
### Bursts: 1632
Top -18 Bottom +9 (Ribo) T1
BVA
### Bursts: 500
Top -18 Bottom +9 (Ribo) T2
BVA
### Bursts: 1398
Top -18 Bottom +9 (Ribo) T3
BVA
### Bursts: 1318
Top -18 Bottom +9 (8-oxoG) T1
BVA
### Bursts: 966
Top -18 Bottom +9 (8-oxoG) T2
BVA
### Bursts: 964
Top -18 Bottom +9 (8-oxoG) T3
BVA
### Bursts: 1057
Top -18 Bottom +9 (abasic) T1
BVA
### Bursts: 1534
Top -18 Bottom +9 (abasic) T2
BVA
### Bursts: 915
Top -18 Bottom +9 (abasic) T3
BVA
### Bursts: 1164
E
$\sigma_i$
0.0 0.2 0.4 0.6 0.8 1.0
0.00 0.05 0.10 0.15 0.20 0.25 0.30 0.35

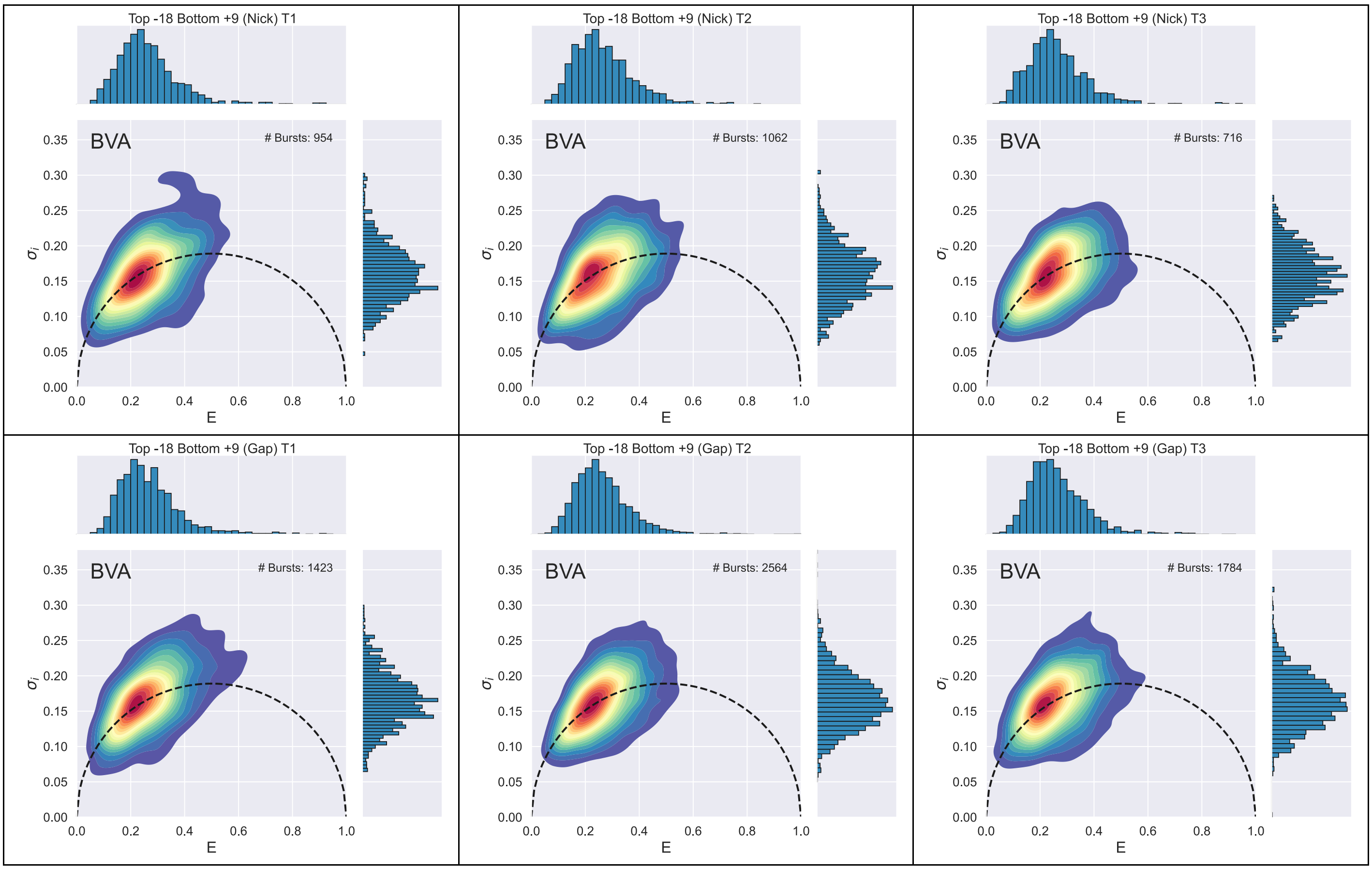

Supplementary Figure 46: BVA for each independent repeat of the seven duplex constructs for the dye-pair Top -18 Bottom +9.

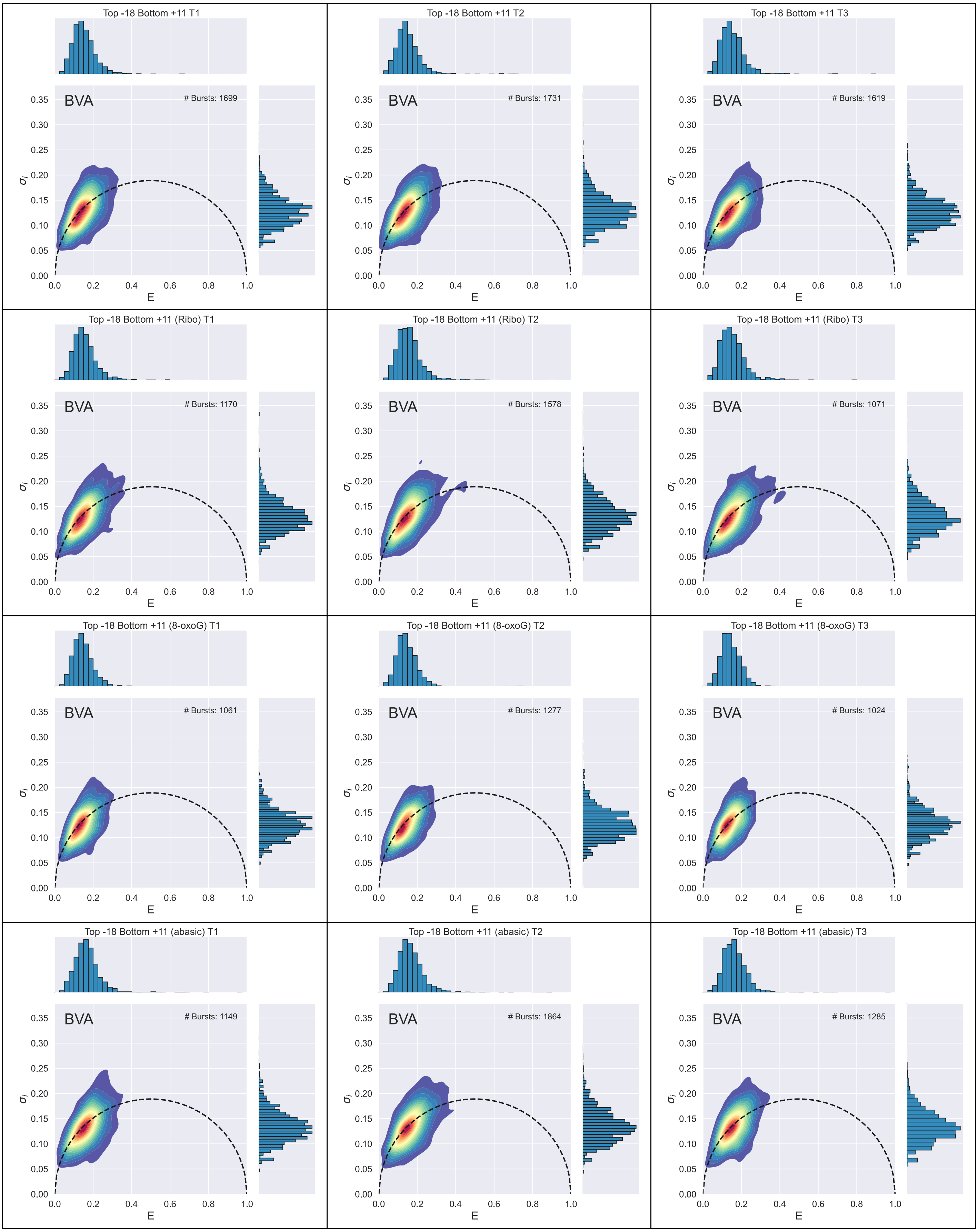
Top -18 Bottom +11 T1
BVA
### Bursts: 1699
Top -18 Bottom +11 T2
BVA
### Bursts: 1731
Top -18 Bottom +11 T3
BVA
### Bursts: 1619
Top -18 Bottom +11 (Ribo) T1
BVA
### Bursts: 1170
Top -18 Bottom +11 (Ribo) T2
BVA
### Bursts: 1578
Top -18 Bottom +11 (Ribo) T3
BVA
### Bursts: 1071
Top -18 Bottom +11 (8-oxoG) T1
BVA
### Bursts: 1061
Top -18 Bottom +11 (8-oxoG) T2
BVA
### Bursts: 1277
Top -18 Bottom +11 (8-oxoG) T3
BVA
### Bursts: 1024
Top -18 Bottom +11 (abasic) T1
BVA
### Bursts: 1149
Top -18 Bottom +11 (abasic) T2
BVA
### Bursts: 1864
Top -18 Bottom +11 (abasic) T3
BVA
### Bursts: 1285
E
$\sigma_i$

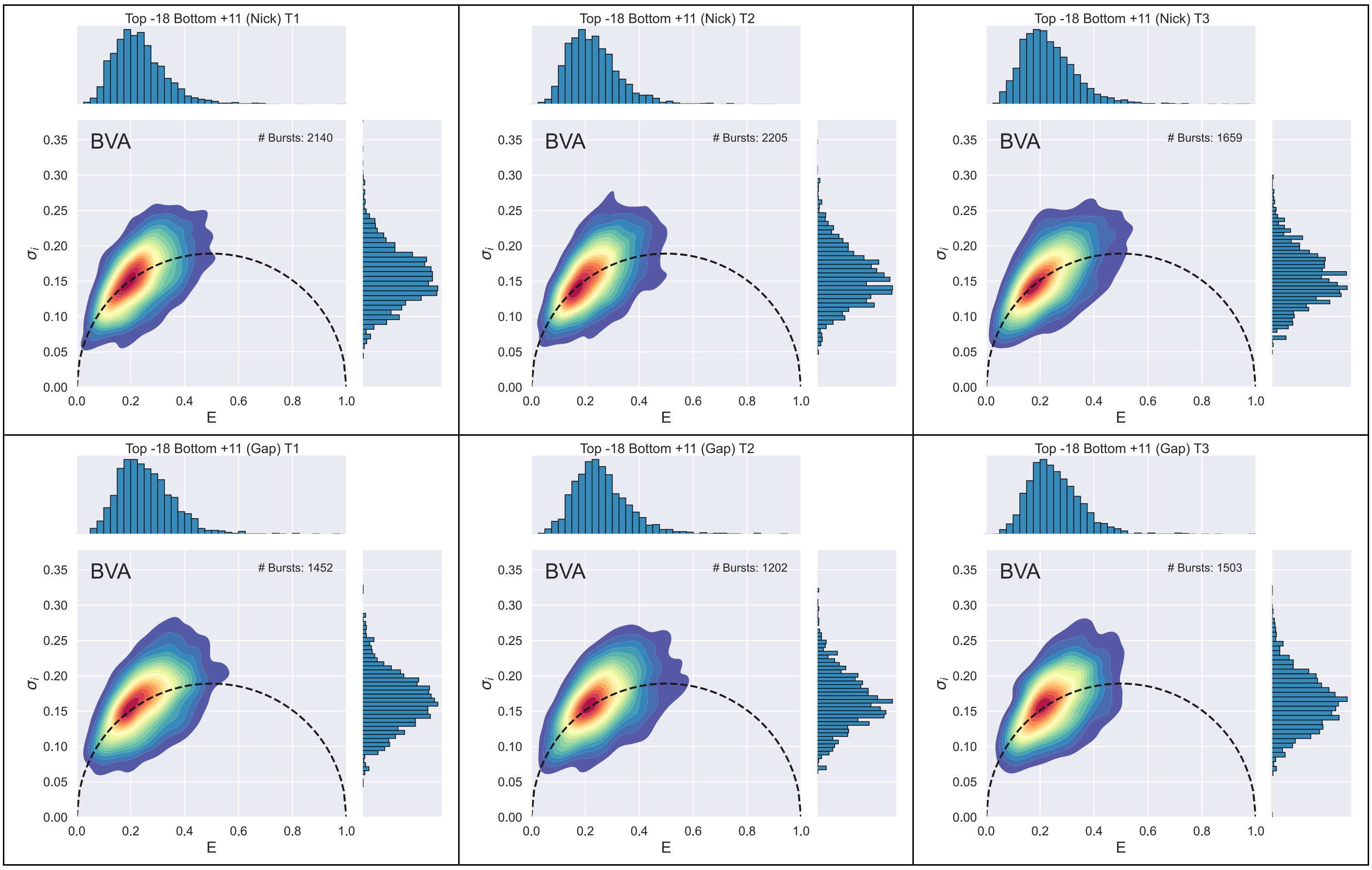

Supplementary Figure 47: BVA for each independent repeat of the seven duplex constructs for the dye-pair Top -18 Bottom +11.
